# Reinforcing Tunnel Network Exploration in Proteins Using Gaussian Accelerated Molecular Dynamics

**DOI:** 10.1101/2024.04.30.591887

**Authors:** Nishita Mandal, Bartlomiej Surpeta, Jan Brezovsky

**Author notes:** **Corresponding authors** Bartlomiej Surpeta, Jan Brezovsky.

## Abstract

Tunnels are structural conduits in biomolecules responsible for transporting chemical compounds and solvent molecules to and from the active site. They have been shown to be present in a wide variety of enzymes across all functional and structural classes. However, the study of such pathways is experimentally challenging because they are typically transient. Computational methods such as molecular dynamics (MD) simulations have been successfully proposed to explore tunnels. Conventional MD (cMD) provides structural details to characterize tunnels but suffers from sampling limitations to capture rare tunnel openings on longer timescales. Therefore, in this study, we explored the potential of Gaussian accelerated MD (GaMD) simulations to improve the exploration of complex tunnel networks in enzymes. We used the haloalkane dehalogenase LinB and its two variants with engineered transport pathways, which are not only well-known for their application potential but have also been extensively studied experimentally and computationally regarding their tunnel networks and their importance in multi-step catalytic reactions. Our study demonstrates that GaMD efficiently improves tunnel sampling and allows the identification of all known tunnels for LinB and its two mutants. Furthermore, the improved sampling provided insight into a previously unknown transient side tunnel (ST). The extensive conformational landscape explored by GaMD simulations allowed us to investigate in detail the mechanism of ST opening. We determined variant-specific dynamic properties of ST opening, which were previously inaccessible due to limited sampling of cMD. Our comprehensive analysis supports multiple indicators of the functional relevance of the ST, emphasizing its potential significance beyond structural considerations. In conclusion, our research proves that the GaMD method can overcome the sampling limitations of cMD for the effective study of tunnels in enzymes, providing further means for identifying rare tunnels in enzymes with potential for drug development, precision medicine, and rational protein engineering.

## Introduction

Enzymes function as natural catalysts to enable the execution of complex reactions within living cells. However, the structural basis of their efficiency and sensitivity remains incompletely understood due to the complexity of their specific biochemical reactions. Enzymatic reactions occur at the active site, which may be either on the surface or buried within the protein’s core. In enzymes with buried active sites, the transport of molecules from the bulk solvent to the active site and their release is carried out through transport pathways known as tunnels^1^. These tunnels are widespread, comprising over 50% of enzymes across all six enzyme commission (EC) classes^2^. To understand the structural basis of activity related to these tunnels, in-depth investigation is required to study the dynamic behavior of these pathways, which determines the exchange rate of substrates entering the active site or products being released into the bulk solvent^3^. The tunnels also regulate the movement of solvents and products^4^, providing additional control over enzymatic reactions.

The tunnels of certain enzymes may have narrow parts, known as bottlenecks, which determine the selectivity of compounds passing to or from the active site. Bottlenecks are primarily controlled through gates represented by residues, loops, secondary structures, or domains^5^. Due to the presence of these gates, the tunnels are transient in nature, making their characterization a non-trivial task and sometimes even impossible in a single static structure^3–5^. Gates are frequently involved in regulating transport, especially in multi-step reactions^4,5^. Gating residues often act as selectivity filters, allowing only certain-sized substrates or products to traverse the tunnel, thereby playing a crucial role in the catalytic activity of the enzyme^5^. Given that many enzymes containing molecular tunnels have been associated with various diseases and that the inhibitors binding to these tunnels can function as effective medications, we can comprehend the biological significance of tunnels^4^. Mutations in tunnel residues can result in variants with significantly altered properties^1,6^. Recent research shows that the catalytic functions of enzymes and their prospects can be altered by the dynamics, geometry, and physicochemical properties of the tunnel^4^. Therefore, the dynamics and flexibility of enzymes and their tunnels must be considered as important factors when studying structural biology in the context of structure-based drug design^4^.

Due to advancements in experimental methods such as X-ray crystallography^7^, cryo-EM^8^, various types of NMR spectroscopy^9^, and cutting-edge computational technologies such as deep learning-based protein structure prediction with AlphaFold2^10^, an increasing number of high-quality three-dimensional static structures are available for in-depth investigation, including the analysis of potential tunnels. Although experimental methods yield the structures of proteins, the study of tunnels within these structures requires dedicated geometry-based tools that explore free van der Waals volume, such as CAVER 3.0^11^, Mole 2.5^12^, and MolAxis^13^. Because static structures derived from experimental techniques or computational predictions usually do not consider multiple protein conformational states, these techniques are often followed by molecular dynamics (MD) simulations to explore the protein conformational landscape and capture tunnel dynamics, thereby studying their transient nature^4,7^. Additionally, the exploration of tunnels by geometry-based methods can be supplemented by ligand-tracking methods using tools such as Streamline tracing^14^, Visual abstraction of Solvent Pathlines^15^, Watergate^16^, AQUA-DUCT^17^, and trj_cavity^18^.

Due to the difficulty of biomolecules easily overcoming large energy barriers, milliseconds to seconds or even longer sampling is required to visit rare structural changes^7,19^. The opening of transient tunnels poses a significant challenge for conventional MD methods (cMD), which are typically limited to tens of microseconds^20^. To address this limitation, various biased enhanced sampling methods have been proposed, such as adaptive biasing force (ABF)^21^, umbrella sampling^22^, metadynamics^23^, and others. Their main limitation lies in defining the collective variables (CV), because this requires thorough knowledge of the system and often extensive optimization to find the optimal solution. For systems with complex tunnel networks, multiple CVs would be required for efficient exploration of the desired transport pathways. On the other hand, accelerated MD (aMD) and Gaussian accelerated MD (GaMD) have been proposed to enhance the sampling of conformational space without explicitly defined CVs^24^, which represent a promising solution for tunnel exploration. Furthermore, GaMD^25^ provides an opportunity to reconstruct the original free energy landscape due to appropriately controlled boosting potential. Although GaMD has been investigated for numerous enzymes, its effectiveness for reliably sampling tunnel dynamics remains unknown.

In this study, we examine the efficiency of GaMD to explore stable and transient tunnels in haloalkane dehalogenase’s (LinB) tunnel network. LinB belongs to the α/β hydrolase superfamily, which catalyzes the hydrolytic dehalogenation of halogenated compounds^26^, with high potential applications in bioremediation, biosensing, or biocatalysis^27,28^. Importantly, due to its multi-step reaction and utilization of water during the reaction, this class of enzymes requires multiple tunnels crucial for synchronizing particular steps, making it a perfect model for our investigation^29^. Structurally, LinB has a flexible cap domain (Figure 1A) and stable core domain^26^, and its active site consists of a catalytic pentad (E132 – a catalytic acid, D108 – a nucleophile, H272 – a base, W109, and N38 – two halide-stabilizing residues)^30^. LinB has three known tunnels: one permanent tunnel (p1) and two auxiliary tunnels (p2 and p3)^6^. Therefore, in the present study, we focused on its three variants to thoroughly test GaMD capabilities to explore the conformational space of the known LinB tunnel network. These variants included (i) LinB wild-type (LinB-Wt), in which p1 represents the main tunnel, whereas p2 tunnels have an auxiliary role (Figure 1B); (ii) designed LinB-Closed carrying the L177W mutation resulting in a drop of p1 tunnel occurrence (Figure 1C)^6^; and (iii) LinB-Open variant, in which the main p1 tunnel was closed by the same mutation L177W as seen in the former variant, resulting in a reduction in p1 functioning as the main tunnel. Introduced mutations W140A, F143L, and I211L result in the opening of an additional auxiliary p3 tunnel (Figure 1D)^6^. Given that the tunnel dynamics and function vary significantly across selected LinB variants, they represent suitable models for the exploration of GaMD capabilities to capture rare tunnel dynamics, validate its efficiency and sensitivity with the broad knowledge from the literature, as well as compare it with cMD simulations.

**Figure 1.**
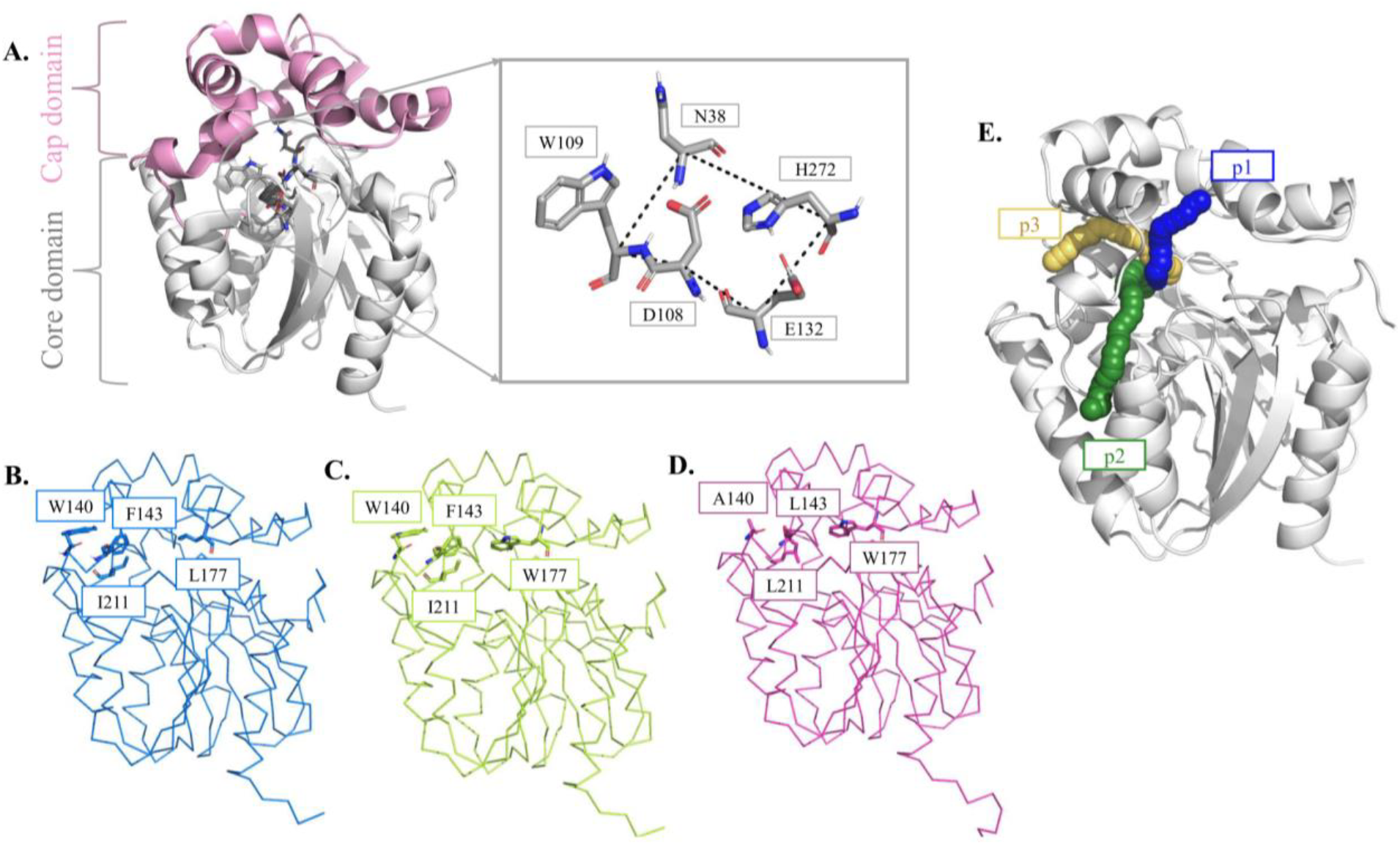
Structural representations of LinB and its variants. **A**. LinB has a more flexible cap domain (pink cartoon) and a more stable core domain (grey cartoon). The catalytic residues of LinB forming the catalytic pentad are shown as grey sticks. **B**. LinB-Wt. **C**. LinB-Closed mutant, carrying the L177W mutation. **D**. LinB-Open mutant with four mutations: L177W, F143L, W140A, & I211L. **E**. Known p1, p2, and p3 tunnels of LinB.

## Methods

### System setup and conventional molecular dynamics simulations

The cMD simulation was performed on three LinB variants: LinB-Wt (PDB code: 1MJ5)^26^, LinB-Closed (PDB code: 4WDQ)^6^, and LinB-Open (PDB code: 5LKA)^6^. First, the systems were protonated using the H++ server^31^ at pH 8.5^6^ and a salinity of 0.1 M. Subsequently, the systems were solvated using 4-point OPC water models^32^ and neutralized with counterions (Na^+^ and Cl^-^) to achieve a final NaCl concentration of 0.1 M. Initially, the systems were energy minimized using 500 steps of steepest descent followed by 500 steps of conjugate gradient with decreasing harmonic restraints in 5 rounds using the PMEMD^33^ module of AMBER18^34^ with the ff14SB force field^35^. The following restraints were applied to the system: 500 kcal mol^-1^Å^-2^on all heavy atoms of the enzyme, followed by 500, 125, 25, and 0 kcal mol^-1^Å^-2^on backbone atoms of the enzyme exclusively. Minimization was followed by a 2 ns equilibration MD simulation, with gradual heating to 310 K under a constant volume using the Langevin thermostat^36^ with a collision frequency of 1.0 ps^-1^ and harmonic restraints of 5.0 kcal mol^-1^Å^-2^on the positions of all enzyme atoms, using periodic boundary conditions with the particle mesh Ewald method^37,38^. The Berendsen barostat was used to control the pressure of the system. The simulations were run using a 4 fs time step enabled by SHAKE and the hydrogen mass repartitioning algorithm^39^. Finally, these simulations were continued with an unrestrained 200 ns production simulation performed with PMEMD.CUDA at constant pressure and temperature, with frames stored every 20 ps.

Clustering analysis was performed using the hierarchical agglomerative (HierAgglo) algorithm in cpptraj^40^ based on a 200 ns trajectory for each of the three systems. A cutoff of 4.5 was used, with nclusters set to 5 and the linkage method as average linkage, to obtain the five most diverse conformations of the system. These five most diverse conformations were used as seed structures for the production cMD, continued to an unrestrained 5 μs simulation, with frames stored every 200 ps, under constant pressure and temperature for all three LinB variants.

### Gaussian accelerated molecular dynamics

The five most diverse structures obtained using clustering analysis in cMD were used as seeds for GaMD in all three LinB variants to diversify the sampling space. After preparing the structures for GaMD simulation, a short 8 ns cMD was run to collect the potential statistics of the system, including maximum (*V*_max_), minimum (*V*_min_), average (*V*_av_), and standard deviation (σV)^25^. GaMD equilibration of 8 ns was performed after adding the boost potential^25^. The first 16 ns of each run were further considered as equilibration and excluded from the final analyses. Parameters governing the strength of the applied boosting potential were set as follows: σ0P, which is the standard deviation of the first potential boost on the total potential energy of the system, was set to 1.3 kcal/mol; and σ0D, which is the standard deviation of the second potential boost on the dihedral angle of the system, was set to 2.5 kcal/mol. These parameters for σ0P and σ0D were tested iteratively using a range of values to bring the coefficients *k*0D and *k*0P as close as possible to a maximum value of 1.0, which provides the highest acceleration to the system, or the highest possible value whenever the system was unstable before reaching the maximum. The parameters used for testing are described in detail in Table S1. The system threshold energy was set to the lower bound (E = *V*_*max*_). Simulations were run using a 4 fs time step, analogously to cMD, with hydrogen mass repartitioning applied^39^. Finally, these dual-boost GaMD simulations were continued for 5 μs of unconstrained production MD. The simulations were performed analogously to cMD settings, namely under constant pressure and temperature using AMBER18 with the ff14SB force field, with frames stored every 200 ps for subsequent analyses.

### Basic analysis

Trajectories generated from both methods, for all three LinB variants and corresponding replicates (totaling 30 repetitions of 5 μs sampling), were processed using cpptraj^40^ implemented in AmberTools17^34^. The root mean square deviation (RMSD) and root mean square fluctuation (RMSF) were calculated with the initial structure as a reference, considering the backbone heavy atoms of the protein (N, CA, C). RMSD calculations excluded the N-terminal tail (comprising the first 11 amino acids) due to its high overall flexibility. The protein’s radius of gyration (Rg)^41^ and solvent accessible surface area (SASA)^42^ were calculated, considering all heavy atoms of the protein. Distances between residues were calculated using the cpptraj, considering the alpha carbons (Cα) of respective residues.

### Tunnel analysis

The tunnels were calculated using CAVER 3.0.2^11^ software with a 6 Å shell radius and 4 Å shell depth. The starting point residues for tunnel calculation were specified as Asn38, Asp109, and His272, corresponding to the numbering from the crystal structure. A probe radius of 0.9 Å was used to calculate potential pathways in the enzymes, with a time sparsity of 1. Finally, the pathways were clustered using a clustering threshold of 3.0. Tunnel calculations were performed across 5 μs simulations of both cMD and GaMD, along with their initial fractions, namely, 2.5, 1, and 0.5 μs GaMD simulations for sampling evaluation and comparison between GaMD and cMD methods. Subsequently, TransportTools^43^ (TT) software v0.9.2 was used to generate a unified tunnel network across all variants. Transport tunnels from 5, 2.5, 1, and 0.5 μs GaMD and cMD simulations in the enzyme variants were compared using the comparative analysis module of TT. The clustering method was set to average, with a clustering cutoff of 1.

### Reweighting of GaMD tunnel profiles

A reweighting protocol was implemented to obtain properties of the investigated tunnels from the original GaMD trajectories, reweighted according to the boost potential applied during simulation. For this purpose, the output files of individual GaMD runs containing the boost potential were parsed along with corresponding TT profiles in CSV format for all superclusters, providing access to tunnel characteristics reweighted to the original free energy landscape without bias. The top-100 tunnels, selected based on reweighted TT throughput from each variant, were subjected to further analysis to study the migration of ligands through them.

### Cryptic pocket detection

To analyze the cryptic and allosteric pockets, three different tools were used. The DeepSite^44^ tool was used to identify viable druggable binding sites on the target protein and pockets likely to bind small molecules. PASSer 2.0^45^ was used to detect probable allosteric site pockets, and FTMove^46^ was used to search for important cryptic binding hotspots utilizing all known conformers of the protein. Web servers for all three tools used the LinB-Wt crystal structure.

### Migration of ligands through GaMD tunnels

To evaluate the efficacy of GaMD tunnels in transporting ligands, CaverDock software v1.1^47^ was used to perform molecular docking across the top tunnels. Four ligands, namely bromide ion (Br^-^), 2-bromoethanol (be), 1,2-dibromoethane (dbe), and water (H_2_O), were used to study the transport events in the tunnels, representing the substrate, products, and water important for the catalytic activity of LinB^6^. MGLTools v1.5.6^48^ was utilized to prepare the inputs using the prepare_receptor4.py and prepare_ligand4.py scripts with default settings. Finally, the upper-bound trajectories obtained from CaverDock were considered to study the energy barriers and the successful migration of ligands through particular tunnels using an in-house Python script^49^.

### Distance-based principal component analysis

Principal component analysis (PCA)^50^ was calculated based on the Cα distances from the side helix (residue IDs: 166–179 in the crystal structure) to the catalytic residue His272 (Figure S1). The Python Scikit-learn package^51^ was used for the calculation of PCA, whereas the distances were calculated using the distance module in cpptraj of AmberTools17. PCA was calculated separately for all three enzyme variants. Cluster analysis was conducted using HDBscan^51^, with the following parameters: min_cluster_size and min_samples were kept at 60, allow_single_cluster was set to True, and cluster_selection_epsilon was set to 0.5.

## Results and Discussion

### GaMD overcomes cMD sampling limitations and enables the exploration of a more diverse conformational space

To determine the optimal acceleration parameters, short GaMD simulations (50 ns) were tested on LinB-Wt, followed by extended simulations (1000 ns). For LinB-Wt, the total potential boost σ0P was set to 1.3, resulting in a *k*0P of 0.09, which avoided prohibitive instability in the system that was observed with higher σ0P values. Regarding the dihedral potential boost, σ0D was set to 2.5, resulting in a *k*0D of 1.0, the maximum possible boost. Consequently, the dual-boost GaMD on LinB-Wt with σ0P=1.3 and σ0D=2.5 was performed for all three variants of LinB (Table S1). The relatively low value of *k*0P for our model system LinB could be attributed to its two domains, namely a more flexible cap domain and a stable core domain. The presence of the flexible cap domain increases the likelihood of multiple tunnels opening in the protein, which can make the enzyme quite sensitive to boost potential energies in GaMD.

To analyze the stability of the enzymes across the simulations, we calculated the RMSD and RMSF for all replicates of simulations from the investigated LinB variants in both cMD and GaMD simulations. The time evolution of RMSD for the wild-type in cMD and GaMD oscillated below ∼2.5 Å, indicating sufficient equilibration and convergence of the simulations (Figure 2A). Similarly, the LinB mutants indicate even more stable behavior of the systems (Figures S2 & S3) and do not display significant increases in RMSD, suggesting more rigid internal dynamics due to the introduced mutations, even in the cap domain, regardless of the applied boosting potential. Furthermore, to verify the compactness of the protein during simulations, especially upon the boosting, we evaluated the Rg and SASA from GaMD trajectories and compared them with cMD, which did not elucidate any significant differences (Figures S4–S6).

**Figure 2.**
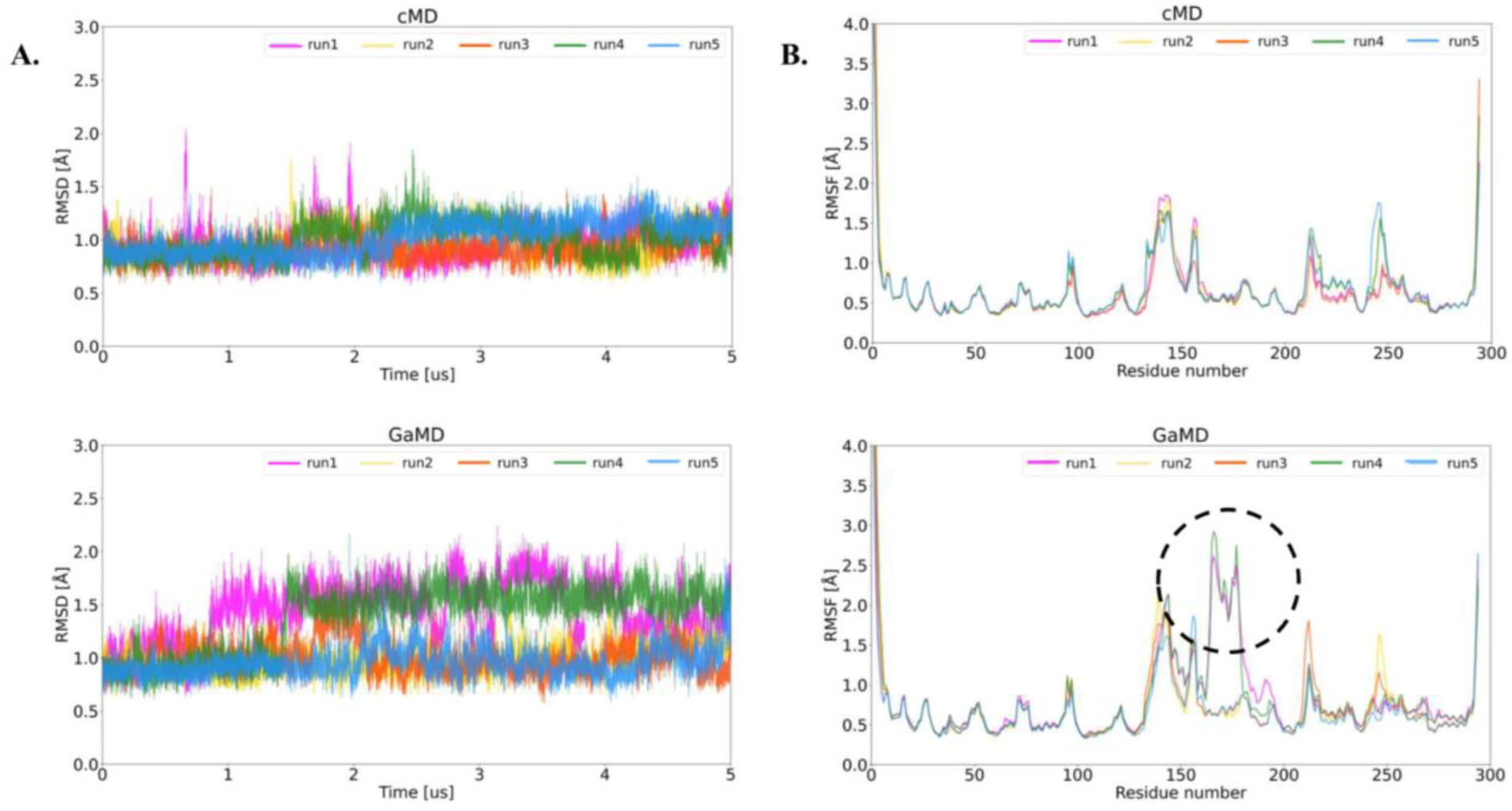
Analysis of protein stability. **A**. The RMSD time evolution of the LinB-Wt enzyme without the 11-residue long N-terminal tail during cMD and GaMD simulations. **B**. The RMSF of LinB-Wt from cMD and GaMD simulations. The most fluctuating residues from the GaMD simulation are highlighted by a black dashed circle.

After verifying the stability of the protein core, we further evaluated the behavior of the residues forming the catalytic machinery. Therefore, we examined the time-evolution RMSD of individual residues Asn38, Trp109, Asp108, Glu132, and His272 forming the catalytic pentad. Collectively, these residues may have been affected by the additional boost potential in GaMD, thus resulting in the sampling of inactive or unphysical conformations (Figures S7–S11). The most flips are observed in the case of residues Asp108, Glu132, and His272. Specifically, for His272 and Glu132, the His272 backbone forms a hydrogen bond with Glu132. However, during the simulation, this hydrogen bond breaks, leading to these residues adopting different conformational states before returning to the initial conformation, or sometimes, the His272 ring enters a different conformational state, causing fluctuations in the RMSD plot (Figures S7– S11). In the case of Asp108, the side chain frequently fluctuates between different conformations, with an RMSD range of approximately 1.25 to 1.50 Å. When considered separately and cumulatively in both cMD and GaMD, the values range between 2.0 to 2.5 Å, indicating that the enzymes’ catalytic machineries are not distorted due to the added boost potential in GaMD.

### GaMD enabled the accurate detection of the tunnel network known for LinB and led to the discovery of a new side tunnel (ST)

We calculated the tunnels to investigate the internal dynamics of ligand transport pathways due to enhanced sampling. The tunnel networks were detected using a combined approach, including CAVER 3.0.2 calculations and further tunnel unification across all simulated systems and replicates in TransportTools. This analysis revealed the presence of all known branches of the p1 and p2 tunnels, namely p1a, p1b, p2a, p2b, and p2c, respectively. Additionally, two rare tunnels – known as p3 and the newly discovered side tunnel (ST) – were found (Figure 3A & B). The newly discovered ST opening corresponds to a region with high RMSD fluctuations, displaying increased values in two replicates of GaMD (especially replicates 1 and 4), due to the boosting potential helping the enzyme to explore a broader sampling space. This observation is consistent with the RMSF profiles showing increased fluctuations in the cap region (residue numbers 166–179) for these corresponding replicates, suggesting improved sampling (Figure 2B). Furthermore, to compare the sampling efficiency between cMD and GaMD during the time evolution of the simulation, we considered various fractions of the trajectories, including the full length (5 μs), first half (2.5 μs), initial 20% (1 μs), and 10% (500 ns). We noted that at least 1 μs of GaMD is required to observe enhanced exploration of tunnels, which was particularly noticeable for the transient tunnel ST (Figures S12–S14). GaMD enhances the exploration of the sampling space to capture the rare tunnels more effectively than cMD (Figure 3A). Besides the tunnel occurrence as defined by the probe radius used for the CAVER calculations (0.9 Å), we also considered other, mostly geometric, properties of the tunnels, such as average length, average bottleneck radius, and maximum bottleneck radius. Interestingly, although tunnels exhibit consistent geometric properties within various groups, it was observed that GaMD simulations tend to open tunnels more, resulting in an increased maximum bottleneck radius exceeding 3 Å (Figures S12–S14).

**Figure 3.**
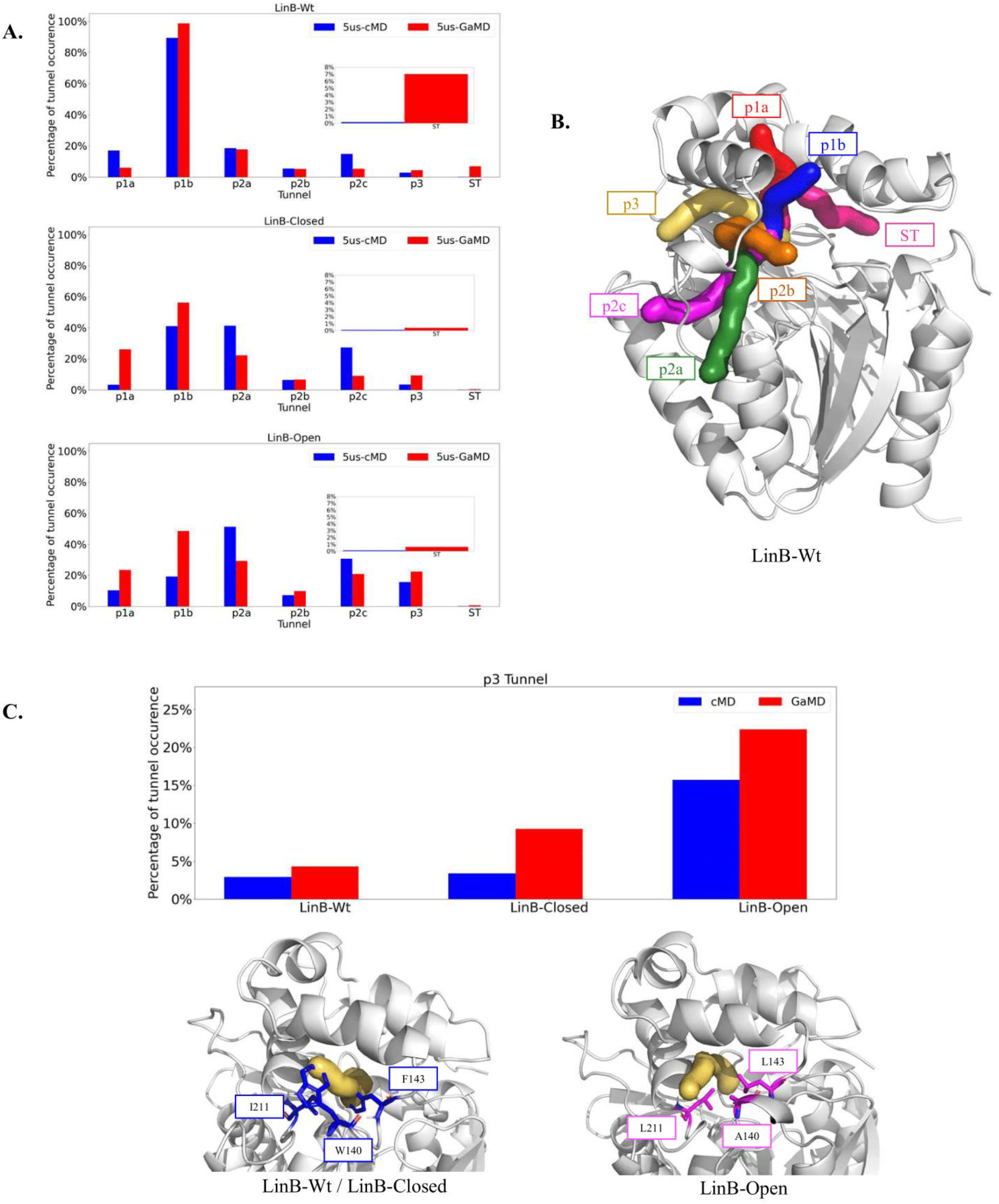
Occurrence of tunnels in LinB variants. **A**. Percentage of tunnels being open in the ensemble for LinB-Wt, LinB-Closed, and LinB-Open variants. Inset plots present the increased sampling of the ST captured by GaMD in LinB-Wt. **B**. Representation of the LinB-Wt crystal structure (grey cartoon) and the tunnel network in the LinB family (colored spheres). **C**. Plot of the p3 tunnel from TT analysis in respective Wt and mutants with cMD and GaMD methods (top) and representation of the protein structure region where the p3 tunnel opens in Wt and LinB-Closed mutant, containing bulky residues TRP, PHE, and ILE; also, the region of p3 tunnel opening in LinB-Open, containing the mutation W140A+F143L+I211L, which leads to the widening of this region (bottom).

The results obtained from GaMD simulations were compared with the tunnel network in three variants with already published data on *de novo* tunnel engineering^6^. We found that the p3 tunnel opening follows a similar trend as in previous studies, that is, the characteristic tunnel network known for each variant is reproduced in the GaMD trajectories. In LinB-Wt, the p1 tunnel branches serve as the primary conduits, which undergo changes in mutants due to the closure of p1 tunnels, resulting in a significant decrease in their occurrence. Additionally, we observed a significant increase in the presence of the p3 tunnel in the LinB-Open variant due to p3-opening mutations. Interestingly, the GaMD tunnels have a tendency for more frequent and broader tunnel openings (Figure 3C), particularly in the case of the p3 tunnel (as indicated by the high bottleneck radius shown in Figures S12–S14). Furthermore, we observed increased sampling of the ST in the wild-type protein, with its occurrence comparable to the rare p3 tunnel in GaMD simulations (Figure 3A), which was not as pronounced in mutants. Moreover, although it is sampled by cMD, we noted its increased sampling, mostly in GaMD for LinB-Wt, whereas the other two variants do not show such prominent ST openings (Figure 3A). Due to large structural perturbation in the helix caused by boosted GaMD simulation, the opening of a transient ST is enhanced. We monitored the RMSF of the protein throughout the simulation and found increased fluctuation in the cap domain in two out of five GaMD simulation replicas, indicating that GaMD visited a broader conformational space.

### Insights into the functional relevance of the discovered side tunnel (ST)

In addition to tunnels, the ligand interaction with enzyme surface pockets or cryptic pockets has been considered important because pocket residues can influence ligand transport through tunnels^52^. Interestingly, Raczyńska et al.^52^ published a study focused on identifying transient binding sites on the enzyme surface as potential sites for engineering enzyme activity. Their study, conducted on LinB-Wt, indicated the ability of the ST entrance pocket to bind 1-chlorohexane. Furthermore, they showed that the experimentally tested mutation of the ST entrance pocket residue A189F increased the enzyme activity by 21.4% for 1-chlorohexane and 26.2% for 1-bromocyclohexane^52^, which highlights the importance of the ST pocket and the newly discovered ST path, which connects the active site or active site pocket to the ST pocket for LinB (Figure 4). To study the opening site of the ST at the enzyme surface and its connectivity with the active site, we used cryptic and allosteric pocket detection tools such as FTMove, PASSer 2.0, and DeepSite. Our analysis revealed a cryptic pocket at the mouth of the ST as one of the major ligand-binding cryptic pockets, suggesting potential allosteric communication via the ST pocket. DeepSite predictions of cryptic pockets shown the ST pocket the second-highest score after the active site p1 tunnel pocket (Figure S15). Similar results were obtained using the PASSer 2.0 allosteric site prediction tool (Figure S16) and FTMove, where the p1 tunnel pocket was ranked third, the active site pocket fourth, and the ST pocket second among all the different cryptic pockets detected by the tool (Figure S17).

**Figure 4.**
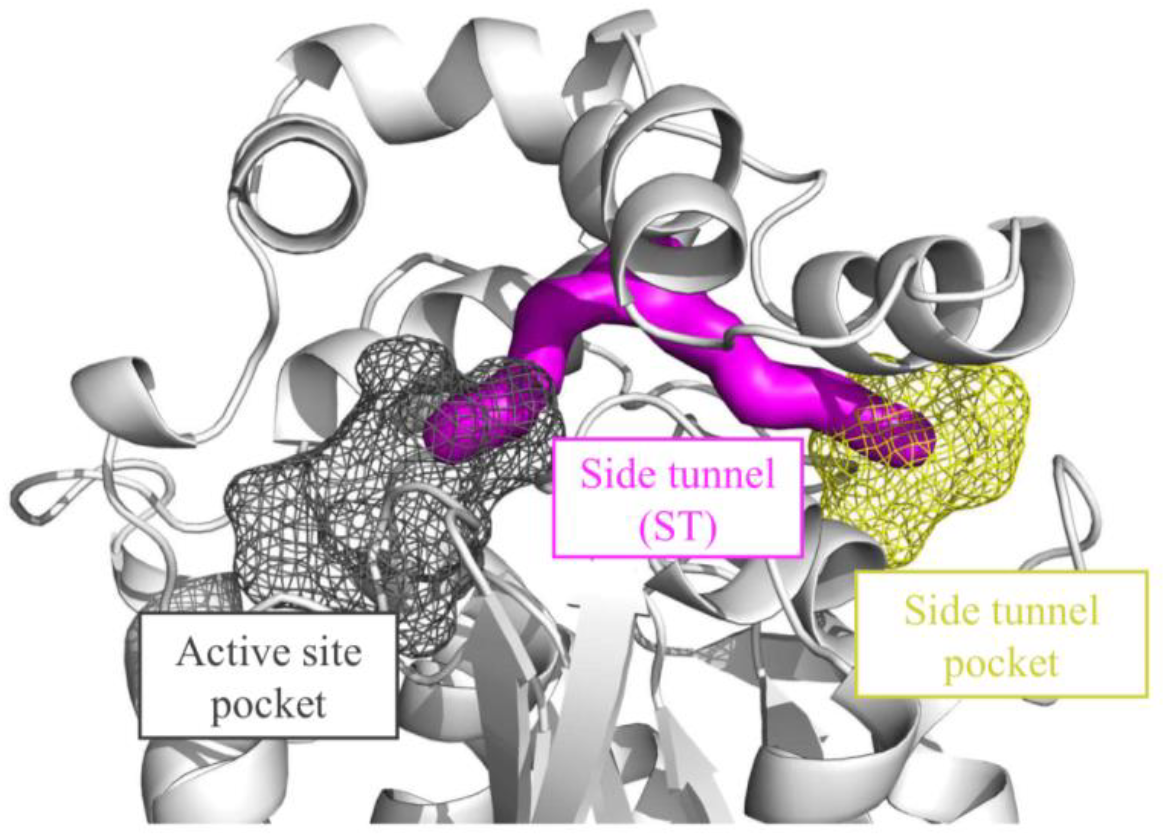
Cryptic pockets detected in the LinB-Wt structure by FTMove. The active site and ST pockets are shown in dark grey and yellow, respectively. The ST pocket is located at the mouth of the ST, which connects these two pockets.

### ST can transport ligands similarly to p3 tunnel

We observed that the ST is connected to cryptic pockets. To study the efficiency of the transient ST in transporting ligands between p3 and p1b, we performed CaverDock calculations using ensembles of the top-100 tunnels with the highest throughput with four substrate and product molecules (Figure 5A). The tunnels used for calculations were obtained from the GaMD trajectories, and using CaverDock, we calculated the energy profile by assessing the binding energy of the ligand to the protein along the entire tunnel length. Additionally, we calculated the energy barriers that the ligands must overcome for successful transport (Supplementary Information File 2). This energetic evaluation of transport demonstrated that the ST was able to transport all four ligands, and interestingly, the energetics of these transports were comparable to the energy barriers sampled for the auxiliary tunnel p3 (Figure 5B). Our evaluation indicated the importance of the ST in conjunction with the known p3 tunnel. Similar observations were obtained for haloalkane dehalogenase DhaA, where a comparable side tunnel was capable of transporting water molecules at levels comparable to p3^53^. Additionally, the energy barrier for water transport (H_2_O) was found to be lower (<5 kcal/mol) in all three tunnels, which shows that all the tunnels were effectively able to transport water through the enzyme. Furthermore, the best-performing tunnel was identified as p1b from the Open mutant, consistent with studies on *de novo* tunnel engineering^6^.

**Figure 5.**
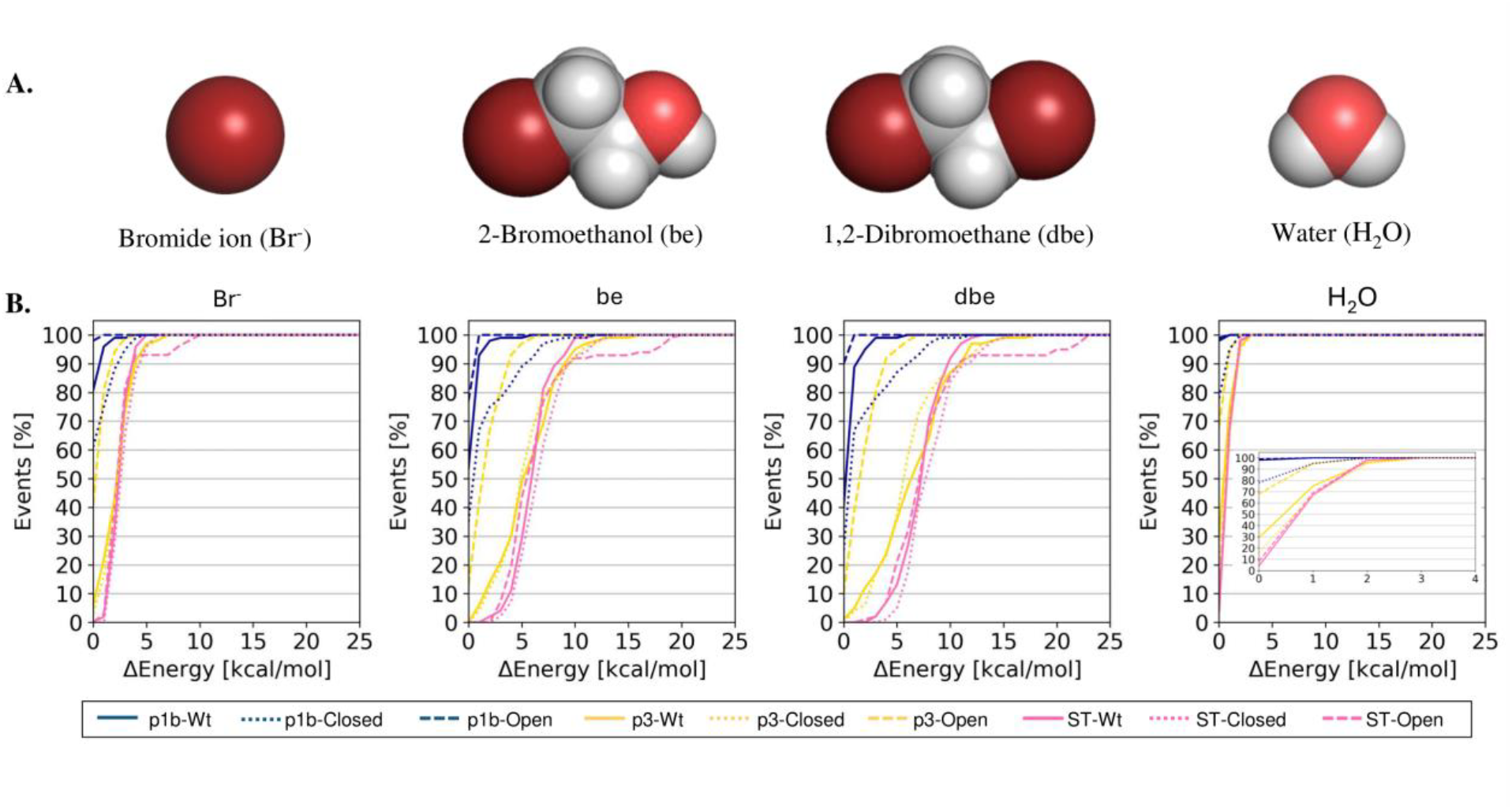
Energetic analysis of ligand transport through the transient ST. **A**. The structure of the four ligands studied: 1,2-dibromoethane (dbe), 2-bromoethanol (be), bromide ion (Br^-^), and water (H_2_O), used for transport analysis (top). **B**. Plots showing the energy costs of successful transport events in all three variants, LinB-Wt, LinB-Open, and LinB-Closed. Blue represents transport events of p1b, yellow represents transport events of p3, and pink shows transport events of the ST (bottom).

### ST opening mechanism is different in LinB-Wt and its mutants

The importance and validity of the ST is confirmed by our results. Therefore, to understand the conformational changes associated with ST opening, we conducted a distance-based PCA in LinB-Wt and its mutants, focusing on the most dynamic region of the protein as confirmed through RMSF. In LinB-Wt, we found that GaMD simulations sample two distinct conformational states in contrast to cMD simulations (Figure 6), indicating that this side helix region provides the protein with the possibility to visit different conformational states (Figure S18). In cMD simulations, we also detected the ST occurrence, but it was less frequent, which confirms that the opening is not forced by the biasing potential but highlights that the system requires broad sampling to capture it efficiently. Major conformational states in all variants were further investigated by HDBscan clustering of distance-based PCA data. We found two predominant states for LinB-Wt in GaMD (Figure 6), which were also partially observed for mutants but only sporadically (Figures S19–S21). Importantly, this analysis further highlights the improved sampling in GaMD trajectories compared to cMD.

**Figure 6.**
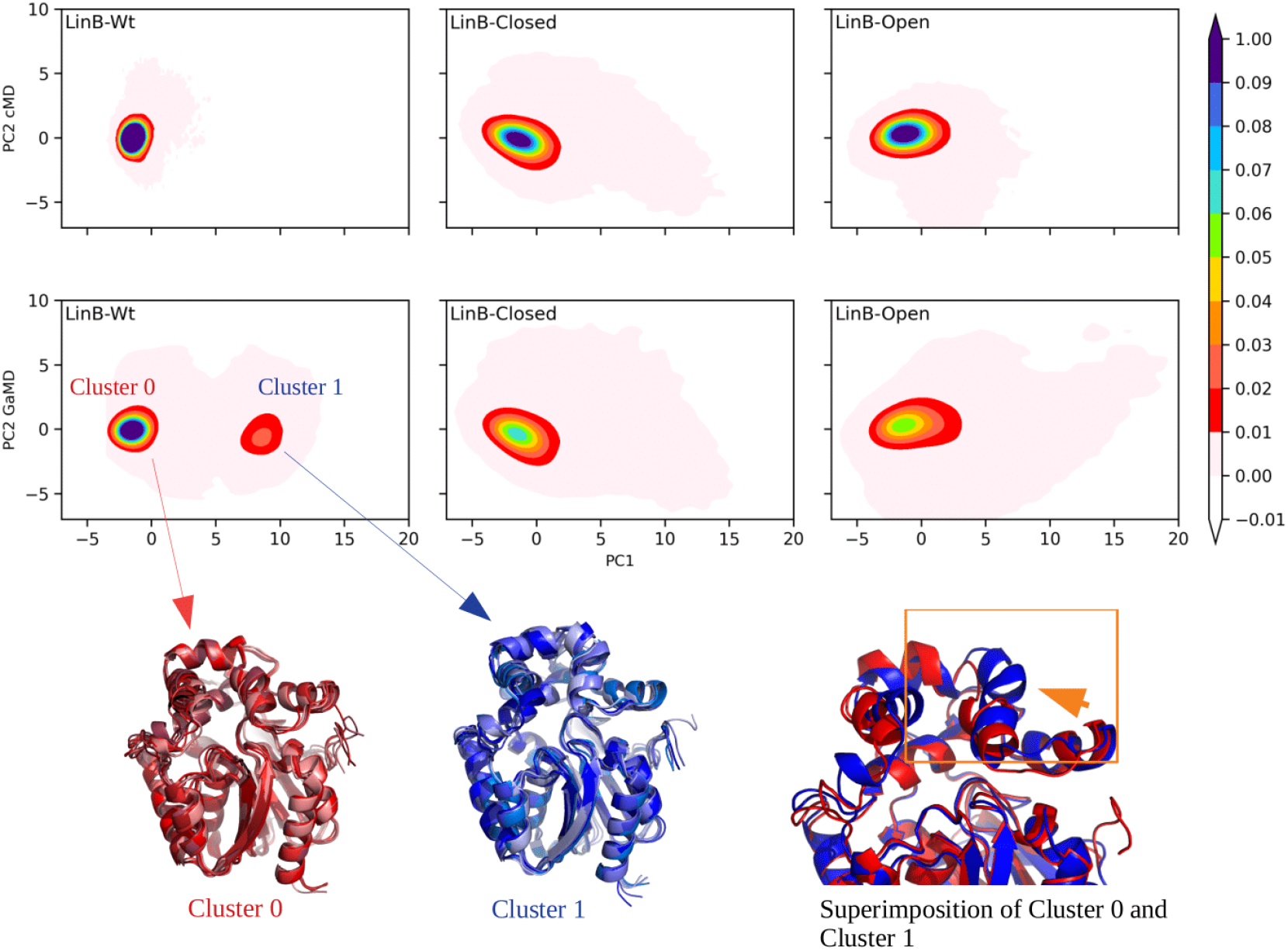
Detection of distinct conformational states sampled by LinB variants. Distance-based PCA analysis: PC1 and PC2 were plotted for all three variants of LinB and represented as Cluster 0 and Cluster 1 (all representative frames) from the LinB-Wt GaMD simulation, showing two different states of the protein. In the superimposed structures, red represents Cluster 0, and blue represents Cluster 1, showing the different conformational states captured by LinB-Wt.

We investigated in-depth the mechanism of the rare ST opening in LinB-Wt and mutants. In LinB-Wt, the prerequisite for the opening of ST is the movement of the side helix away from the protein cap domain. Due to the high fluctuation of the side helix of the cap domain, the ST opens in LinB-Wt frequently (Figure 7A & S22), especially in GaMD-boosted simulations. In the case of mutants (LinB-Closed and LinB-Open), a mutation at the bottleneck residue of the ST (L177W) introduces hydrogen bonding with D147, and the breakage of this hydrogen bond promotes the opening of the ST. However, this process is notably more energetically unfavorable, resulting in the ST being less frequently open in mutants compared to the wild-type enzyme (Figure 7B & S23–S24). We had already observed that LinB-Wt samples the ST more frequently than mutants because, in the case of LinB-Wt, the lack of a hydrogen bond donor makes it easier for the movement of the side helix away from the protein, contributing to the opening of the side tunnel. In contrast, in mutants, due to hydrogen bonding, the opening is less favorable and, therefore, less frequent.

**Figure 7.**
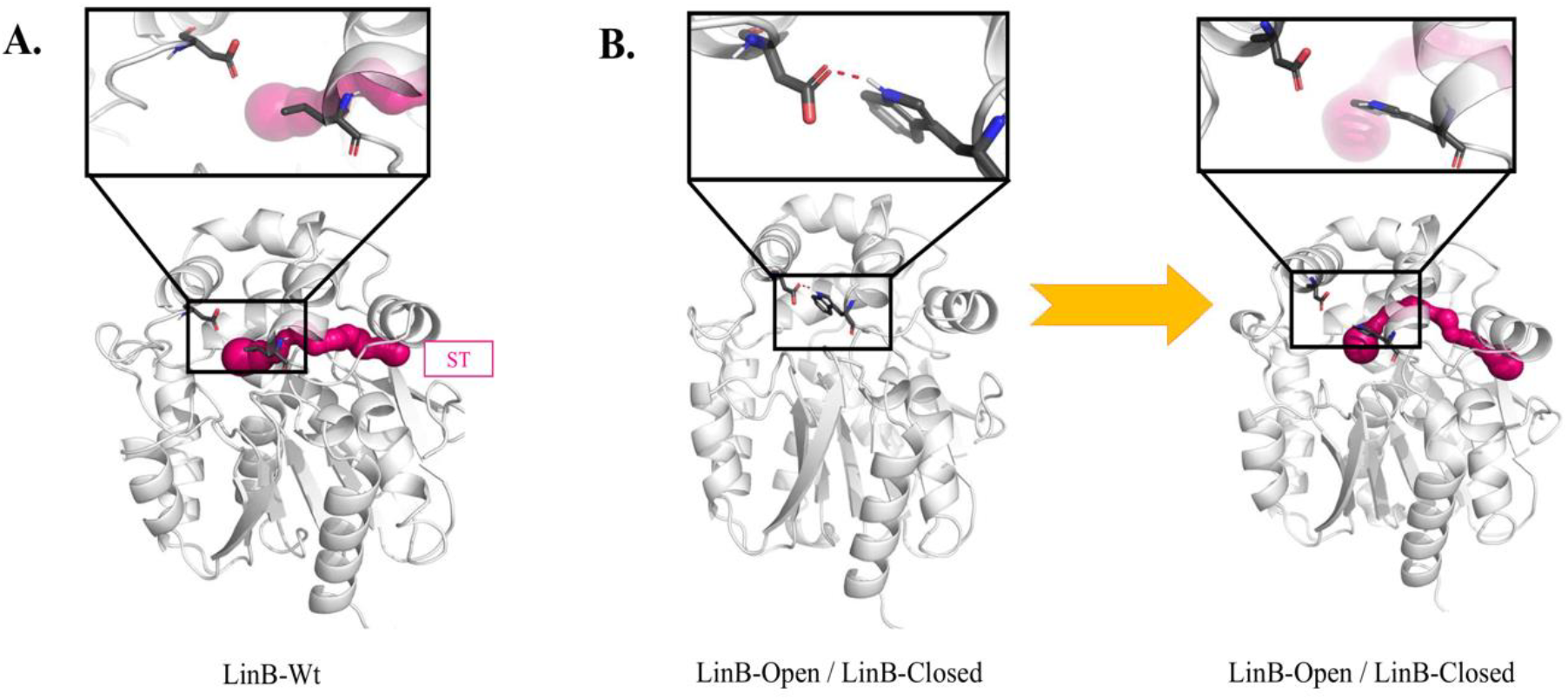
Mechanism of side tunnel opening. **A**. Representation of the ST in the LinB-Wt protein, showing the necessary movement of the side helix for the opening of the ST. **B**. Representation of the ST in mutants, showing the necessary movement of the side helix accompanied by the breakage of the hydrogen bond between Trp176 and Asp146.

## Discussion

In this study, we discussed the potential application of GaMD enhanced sampling method to investigate tunnels and ligand transport pathways in LinB dehalogenases. Our findings demonstrate that GaMD is not only able to identify all known tunnels (p1a, p1b, p2a, p2b, and p2c) comparable to cMD method but also enhances sampling of the enzyme conformational space and visits rare tunnels (ST and p3 tunnel). The improved sampling provided by GaMD enabled us to identify a previously unexplored ST and to further understand the mechanism of its opening in three LinB variants. This opening mechanism was found to be clear and straightforward in the case of the LinB-Wt enzyme, but more complex in mutants. Importantly, such a thorough investigation of the newly discovered ST pathway would not have been possible without the sufficient conformational exploration provided by GaMD. What is critically important is this method not only provided a better picture of conformational dynamics for the three LinB variants but also facilitated consistent exploration of known tunnels from broad computational and experimental data for all variants.

Our exploration revealed that in LinB-Wt, the movement of the side helix is the main contributing factor to the opening of the ST pathway. For mutants, the sampling of ST pathways is less pronounced because of the L177W mutation, resulting in a hydrogen bond between the introduced W177 and D147. This prohibits the opening of the helix observed in the wild-type enzyme and proves the ST is mostly unfavorable in mutants. Thus, it is rarely observed in standard and enhanced MD simulations. Furthermore, the functional importance of this ST pathway is supported by the exploration of druggable, cryptic, and allosteric pockets, all consistently pointing to the mouth of the ST pathway as the functional site. This is further corroborated by a recent experimental study demonstrating the ability of the ST pathway mouth to bind drugs^52^. Additionally, migration analysis using CaverDock with GaMD tunnels confirmed the capacity of the transient ST to transport the substrate and product molecules efficiently at levels comparable to the auxiliary tunnel p3, which supports the auxiliary role of the ST in LinB.

Regarding the p3 tunnel, we observed that GaMD provides better sampling compared to standard MD simulations. Furthermore, we noted the trend expected for LinB-Wt and mutant enzymes from previous literature data. The p3 tunnel is detected more frequently in the LinB-Open mutant due to the mutations W140A+F143L+I211L, which replace bulky residues with smaller ones, aiming to provide more space for the opening of the p3 tunnel. Conversely, in LinB-Wt and the LinB-Closed mutant, the amino acids at the mouth of the p3 tunnel are not modified, resulting in a more restricted opening compared to the LinB-Open variant.

## Conclusions

Our study demonstrates GaMD as a practically useful approach for investigating tunnels in proteins. By overcoming the sampling limitations of standard MD simulations, GaMD can more effectively explore the conformational dynamics of the protein under study while limiting the computational costs. This opens up new possibilities for identifying tunnels by efficiently sampling rare events and overcoming unfavorable energy barriers. Therefore, as shown for the LinB enzyme in our study, this method is suitable for effectively exploring new pathways that can become a druggable site to target, in addition to the conventional targeting of the active site itself.

### Associated Content

## Supporting Information

The following files are available free of charge:

- Testing parameters for GaMD simulations; region used for distance-based PCA calculation; RMSD and RMSF of mutant enzymes; Rg and SASA of LinB-Wt and its mutants; RMSD of catalytic residues; comparison between cMD and GaMD tunnel properties from TransportTools; the results of cryptic and allosteric pocket detecting tools; plots showing PCA comparison between cMD and GaMD; HDBscan cluster analysis of LinB-Wt and its mutants; mechanism of side tunnel opening using helix movement in LinB-Wt; mechanism of side tunnel opening using helix movement and hydrogen bond analysis in the mutant enzymes (PDF).
- Supporting Information File 2 with upper bound energy profiles using CaverDock from individual 100 tunnels in all variants with all four ligands (PDF).
- Underlying data are available on the Zenodo repository: (i) https://zenodo.org/doi/10.5281/zenodo.11092891 containing input, output, and analysis files; and (ii) https://zenodo.org/doi/10.5281/zenodo.11093856 containing the parameter files and stripped MD trajectory files.

## Author Information

### Corresponding Authors

* (J.B.) & (B.S.)

#### Author Contributions

N.M. performed all MD simulations, calculations of tunnels, cryptic pocket analysis, PCA, and migration analysis and drafted the manuscript. B.S. devised the tunnel reweighting, PCA, and clustering analysis protocol. J.B. and B.S. devised and supervised the project. N.M. and B.S. analyzed the data. J.B., N.M., and B.S. interpreted the results. The manuscript was written through the contributions of all authors. All authors have approved the final version of the manuscript.

#### Funding Sources

This work was supported by the National Science Centre, Poland (Grant Number 2017/26/E/NZ1/00548).

#### Notes

The authors declare no competing financial interest.

## Acknowledgments

The computations were performed at the Poznan Supercomputing and Networking Center.

## Abbreviations

MD: Molecular Dynamics
cMD: Conventional MDs;
GaMD: Gaussian accelerated MD;
RMSD: Root Mean Square Deviation;
RMSF: Root Mean Square Fluctuation;
TT: TransportTools;
ST: Side Tunnel;
EC: Enzyme Commission;
ABF: Adaptive Biasing Force
CV: Collective Variable;
NMR: Nuclear Magnetic Resonance;
aMD: Accelerated MD;
PC1: Principal Component 1
PC2: Principal Component 2
cryo-EM: Cryogenic Electron Microscopy.

## Table of Contents Graphic

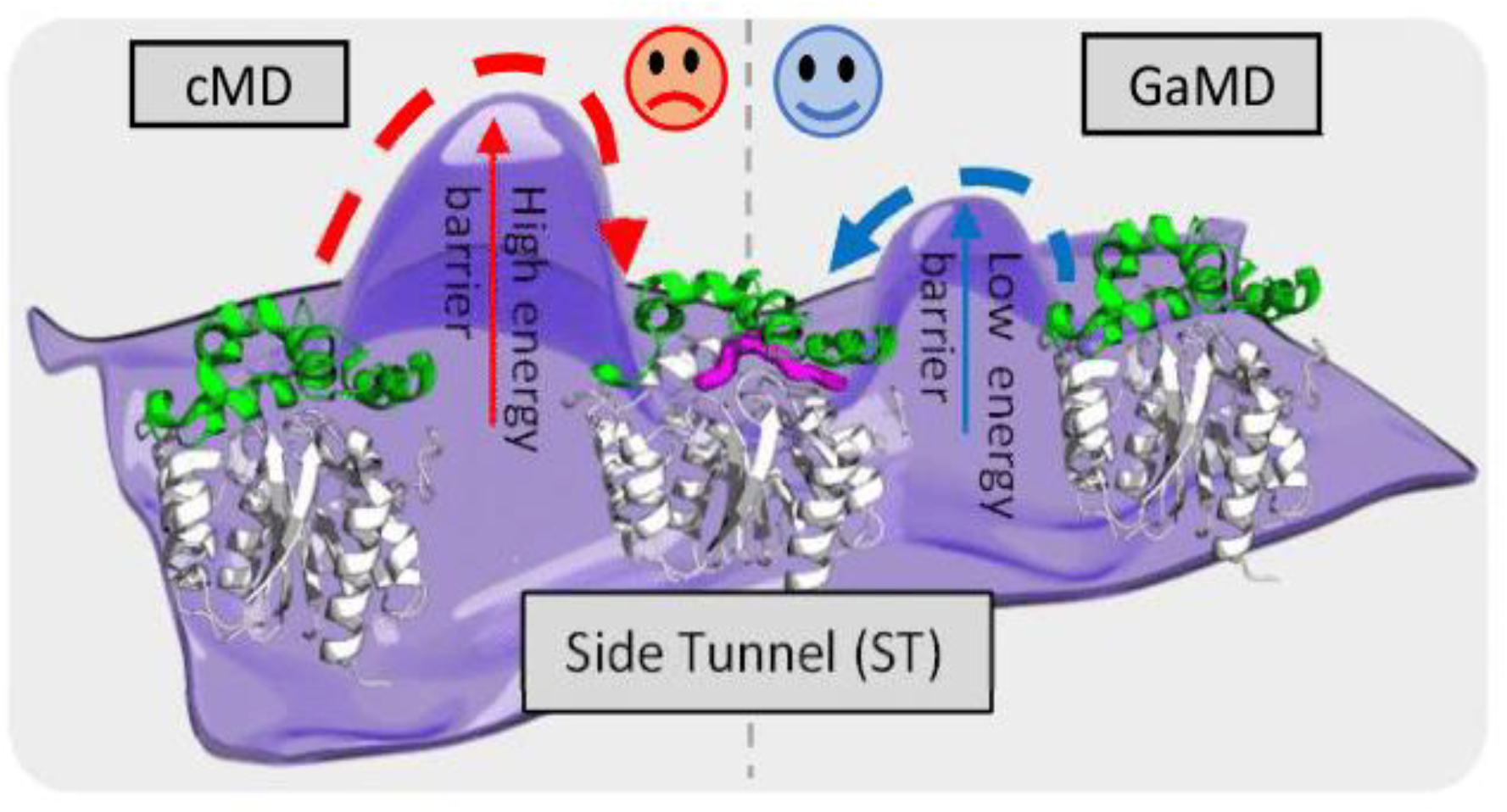

