## Supplementary figures and tables for "Reinforcing Tunnel Network Exploration in Proteins Using Gaussian Accelerated Molecular Dynamics"

**Corresponding authors**

transport tunnel, molecular dynamics, haloalkane dehalogenase, enhanced sampling methods, mechanism

**Table S1.** Testing σ0P and σ0D parameters for dual boost GaMD setup.

| **σ0P/σ0D (kcal/mol)** | $k_0$P/$k_0D$ |
|---|---|
| **GaMD simulations – 66ns** | |
| **Test 1** | |
| 6.0/6.0 | simulation crashed |
| 5.0/5.0 | simulation crashed |
| 4.0/4.0 | simulation crashed |
| 3.0/3.0 | simulation crashed |
| 2.0/2.0 | simulation crashed |
| 1.0/1.0 | 0.388/0.1296 |
| **Test 2** | |
| 1.0/1.1 | 0.0425/0.1389 |
| 1.0/1.2 | 0.0423/0.1650 |
| 1.0/1.3 | 0.0356/0.2133 |
| 1.0/1.4 | 0.0379/0.2196 |
| 1.0/1.5 | 0.0411/0.2718 |
| 1.0/1.6 | 0.0382/0.2731 |
| 1.0/1.7 | 0.0376/0.3417 |
| 1.0/1.8 | 0.0386/0.4394 |
| 1.0/1.9 | 0.0318/0.4541 |
| **Test 3** | |
| 2.0/1.2 | simulation crashed |
| 3.0/1.2 | simulation crashed |
| 4.0/1.2 | simulation crashed |
| 2.0/1.4 | simulation crashed |
| 3.0/1.4 | simulation crashed |
| 4.0/1.4 | simulation crashed |
| 2.0/1.6 | simulation crashed |
| 3.0/1.6 | simulation crashed |
| 4.0/1.6 | simulation crashed |
| 2.0/1.8 | simulation crashed |
| 3.0/1.8 | simulation crashed |
| 4.0/1.8 | simulation crashed |
| **Test 4** | |
| 1.5/1.0 | 0.1013/0.1326 |
| 2.0/1.0 | simulation crashed |
| 2.5/1.0 | simulation crashed |
| 3.0/1.0 | simulation crashed |
| 3.5/1.0 | simulation crashed |
| 4.0/1.0 | simulation crashed |

| | |
|---|---|
| 1.5/0.8 | 0.0986/0.1029 |
| 2.0/0.8 | simulation crashed |
| 2.5/0.8 | simulation crashed |
| 3.0/0.8 | simulation crashed |
| 3.5/0.8 | simulation crashed |
| 4.0/0.8 | simulation crashed |
| 1.5/0.6 | 0.0908/0.0685 |
| 2.0/0.6 | simulation crashed |
| 2.5/0.6 | simulation crashed |
| 3.0/0.6 | simulation crashed |
| 3.5/0.6 | simulation crashed |
| 4.0/0.6 | simulation crashed |
| 1.5/0.4 | 0.0820/0.0430 |
| 2.0/0.4 | simulation crashed |
| 2.5/0.4 | simulation crashed |
| 3.0/0.4 | simulation crashed |
| 3.5/0.4 | simulation crashed |
| 4.0/0.4 | simulation crashed |
| **Test 5** | |
| 1.0/2.5 | 0.0392/1.0000 |
| 1.1/2.5 | 0.0454/1.0000 |
| 1.2/2.5 | 0.0596/1.0000 |
| 1.3/2.5 | 0.0698/1.0000 |
| 1.4/2.5 | 0.0938/1.0000 |
| 1.5/2.5 | 0.1013/1.0000 |
| 1.6/2.5 | 0.1133/1.0000 |
| 1.7/2.5 | 0.1415/1.0000 |
| 1.8/2.5 | simulation crashed |
| 1.9/2.5 | simulation crashed |
| 2.0/2.5 | simulation crashed |
| **Extended GaMD simulations – 516ns** | |
| **Test 6** | |
| 1.5/2.5 | 0.0937/1.0000 |
| 1.7/2.5 | Simulation crashed |
| **Restart simulation – 500ns** | |
| 1.5/2.5 | 0.0937/1.0000 |
| **Further Extended GaMD simulation – 1016ns** | |
| **Test 7** | |
| 1.5/2.5 | Simulation crashed |
| 1.5/2.5 | 0.1001/1.0000 |
| 1.5/2.5 | Simulation crashed |

| | |
|---|---|
| 1.5/2.5 | 0.0928/1.0000 |
| 1.5/2.5 | 0.0955/1.0000 |
| 1.4/2.5 | 0.0776/1.0000 |
| 1.4/2.5 | 0.0926/1.0000 |
| 1.4/2.5 | Simulation crashed |
| 1.4/2.5 | 0.0825/0.9838 |
| 1.4/2.5 | 0.0867/1.0000 |
| 1.3/2.5 | 0.0803/1.0000 |
| 1.3/2.5 | 0.0759/1.0000 |
| 1.3/2.5 | 0.0665/1.0000 |
| 1.3/2.5 | 0.0804/1.0000 |
| 1.3/2.5 | 0.0734/1.0000 |

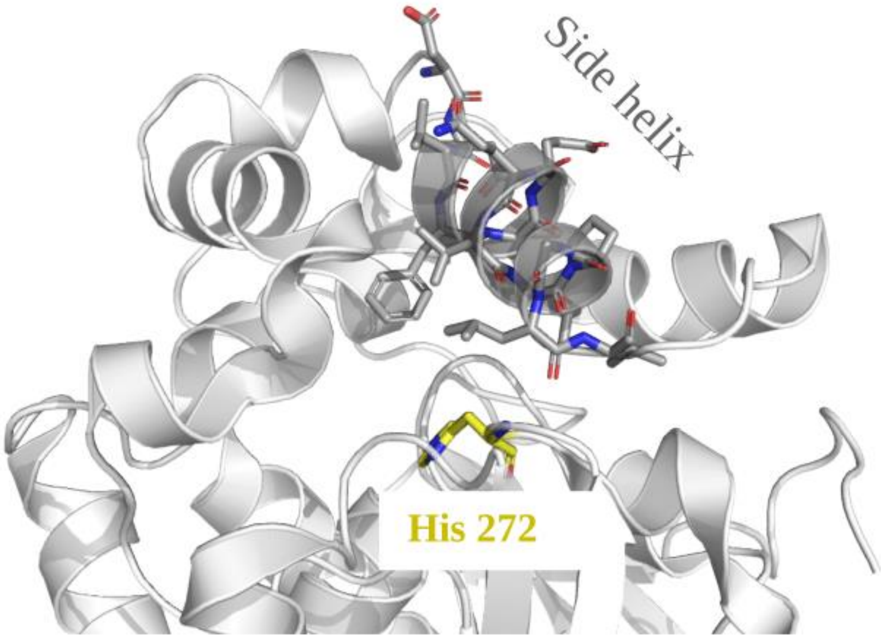

**Figure S1.** Region from the side helix from where (Cα) distance based PCA is calculated to His 272.

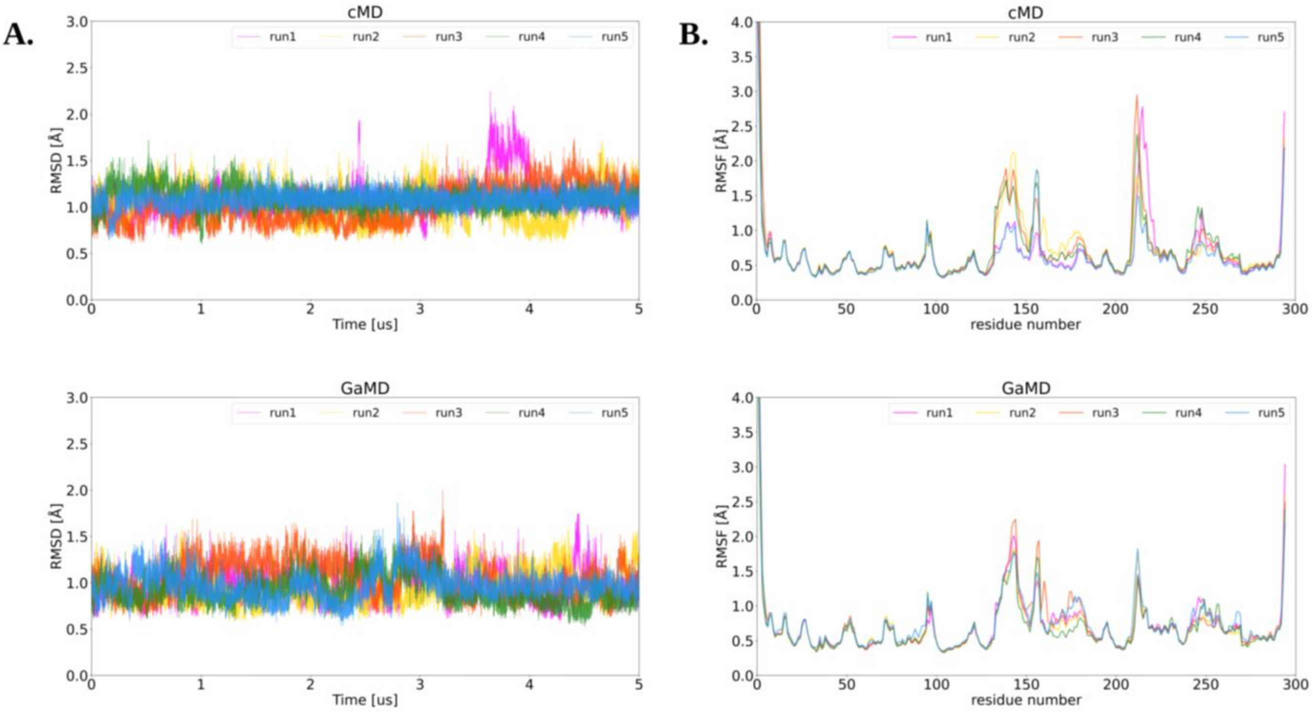

**Figure S2.** RMSD and RMSF of mutant LinB-Closed: (A) RMSD calculated on protein after removal of N-terminal tail (11 amino acids) using cMD and GaMD, (B) RMSF calculated on whole protein residues.

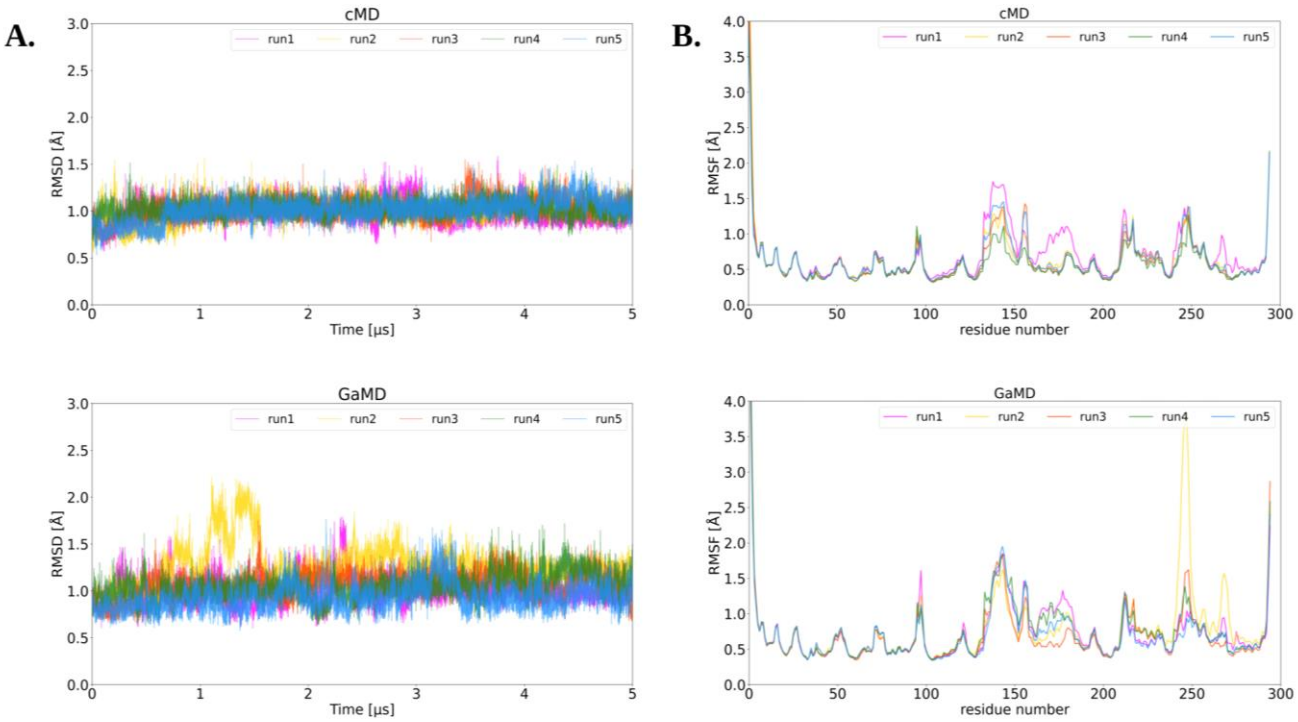

**Figure S3.** RMSD and RMSF of mutant LinB-Open: (A) RMSD calculated on protein after removal of N-terminal tail (11 amino acids) using cMD and GaMD, (B) RMSF calculated on whole protein residues. The high fluctuation in the second replicate's (run2) values in RMSD and RMSF plots of GaMD are caused by residues of loop domain (245GLY – 249GLY). This can occur due to boost potential of GaMD, which increased the fluctuations of the loop and the residues in the vicinity. Importantly, the RMSD remains in the range of 2Å and the loop region returns close to its initial, stable conformation after second μs indicating that observed movement represent reversible fluctuation without serious effect on the overall dynamics of protein.

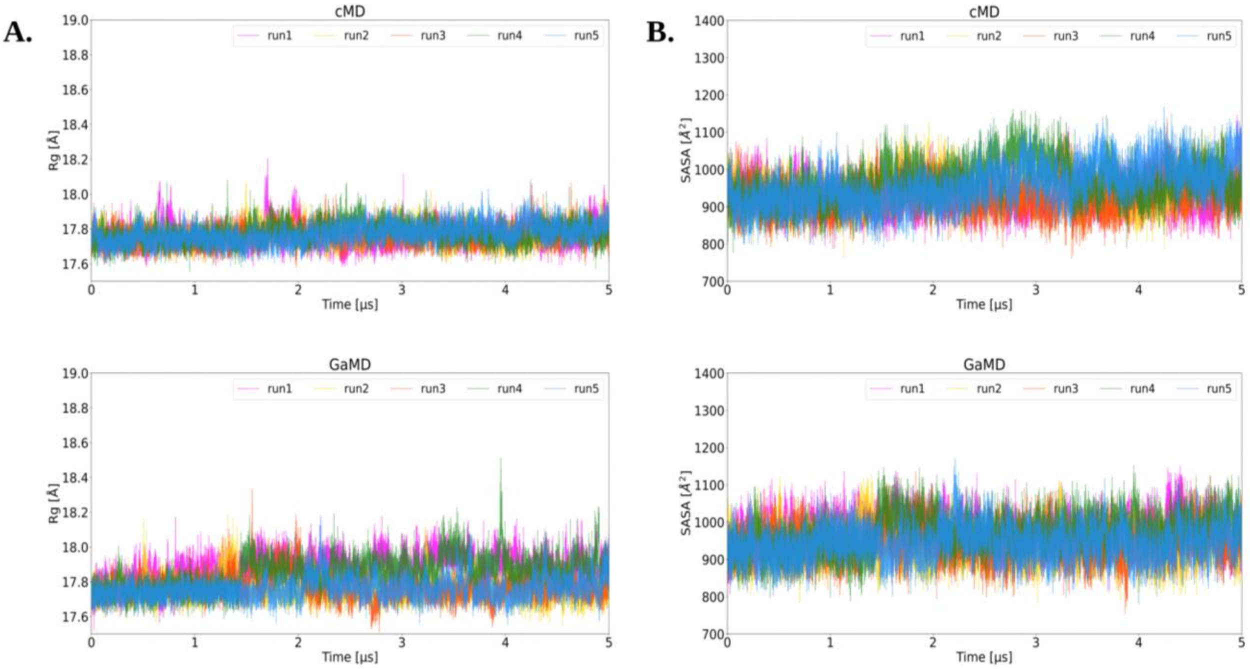

**Figure S4.** Rg and SASA of LinB-Wt: (A) Rg calculated on whole protein using cMD and GaMD, (B) SASA calculated on whole protein using cMD and GaMD.

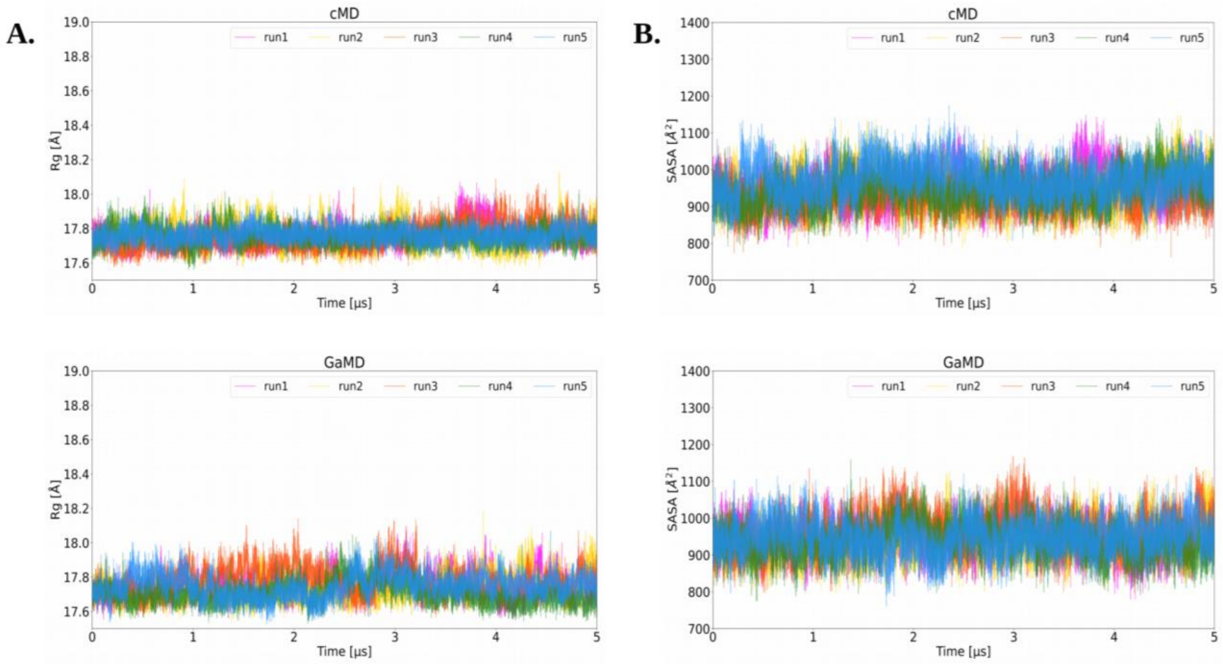

**Figure S5.** Rg and SASA of LinB-Closed: (A) Rg calculated on whole protein using cMD and GaMD, (B) SASA calculated on whole protein using cMD and GaMD.

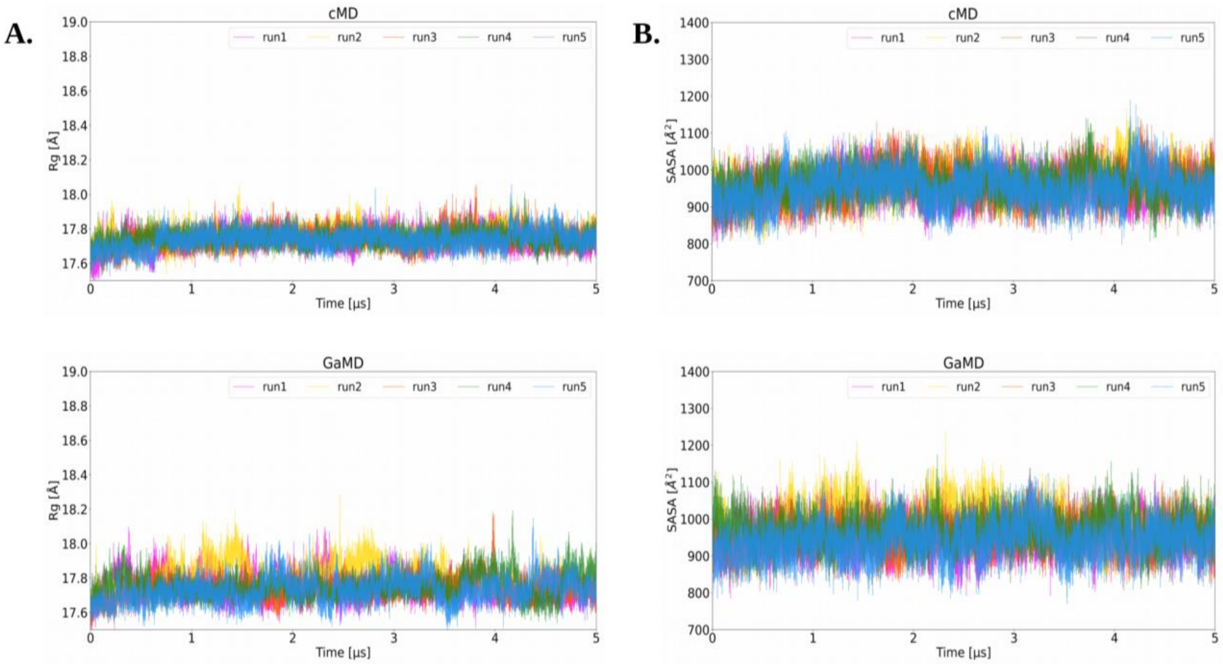

**Figure S6.** Rg and SASA of LinB-Open: (A) Rg calculated on whole protein using cMD and GaMD, (B) SASA calculated on whole protein using cMD and GaMD.

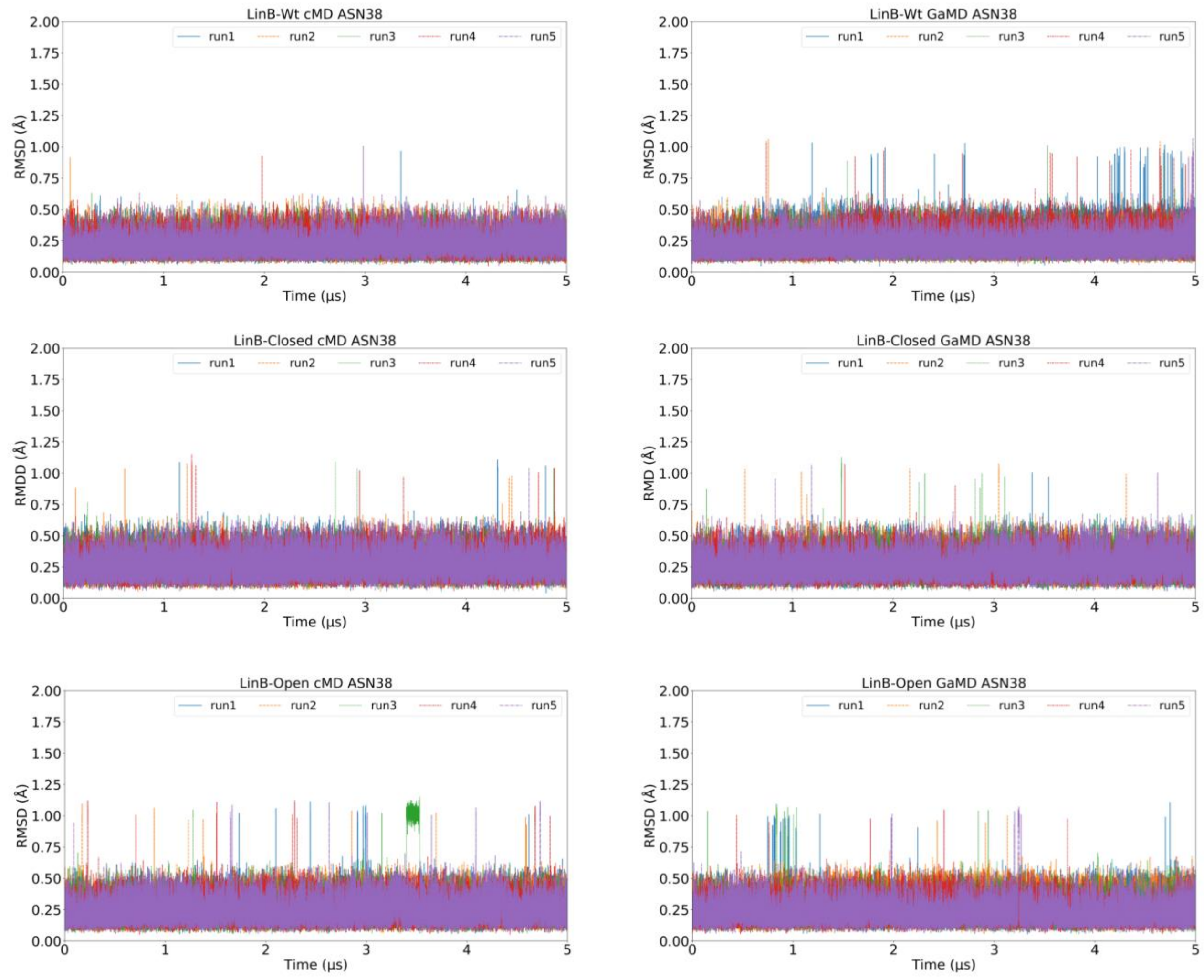

**Figure S7.** RMSD of catalytic residues ASN38 in all variants of LinB, to investigate its stability in the cMD and GaMD.

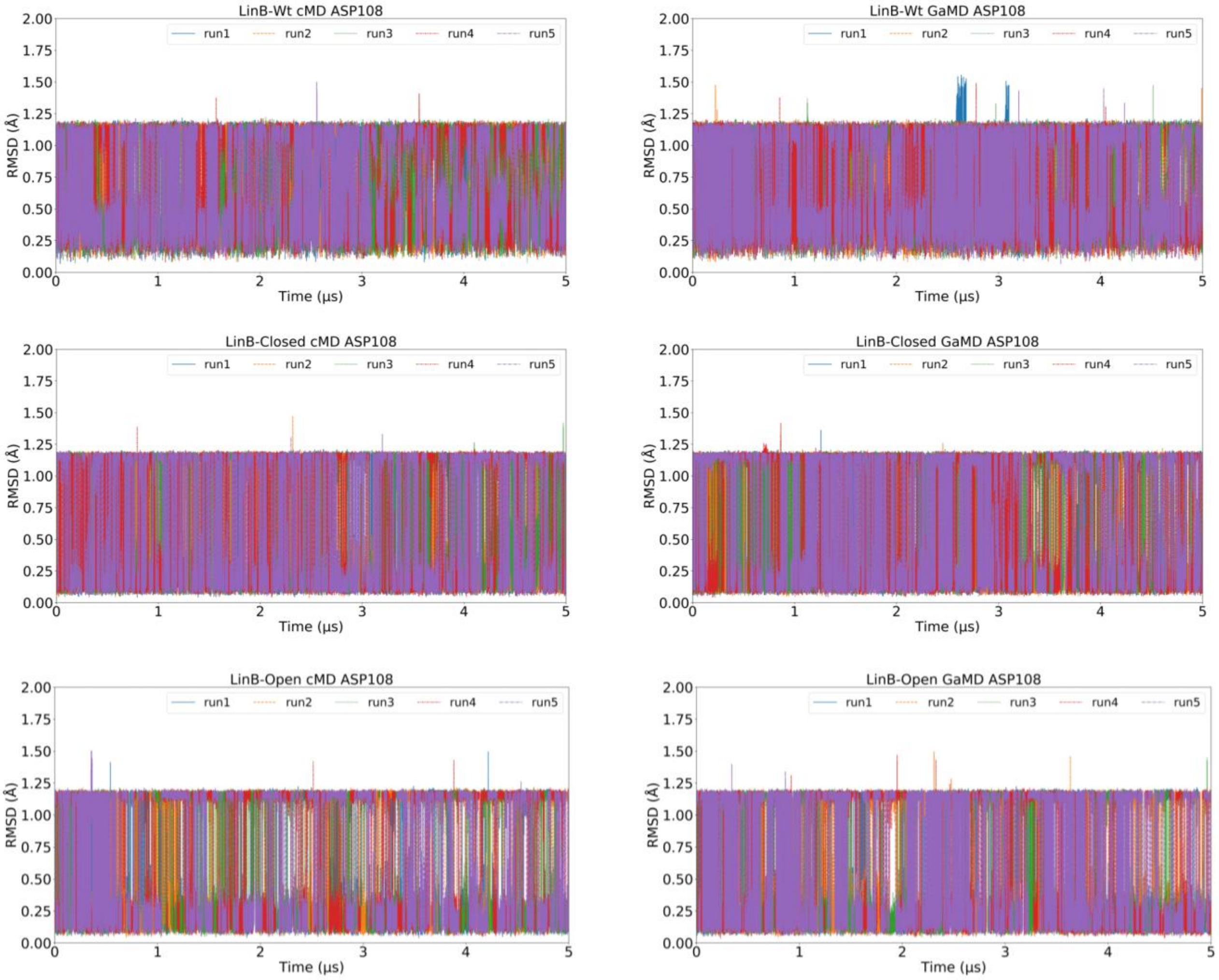

**Figure S8.** RMSD of catalytic residues ASP108 in all variants of LinB, to investigate its stability in the cMD and GaMD.

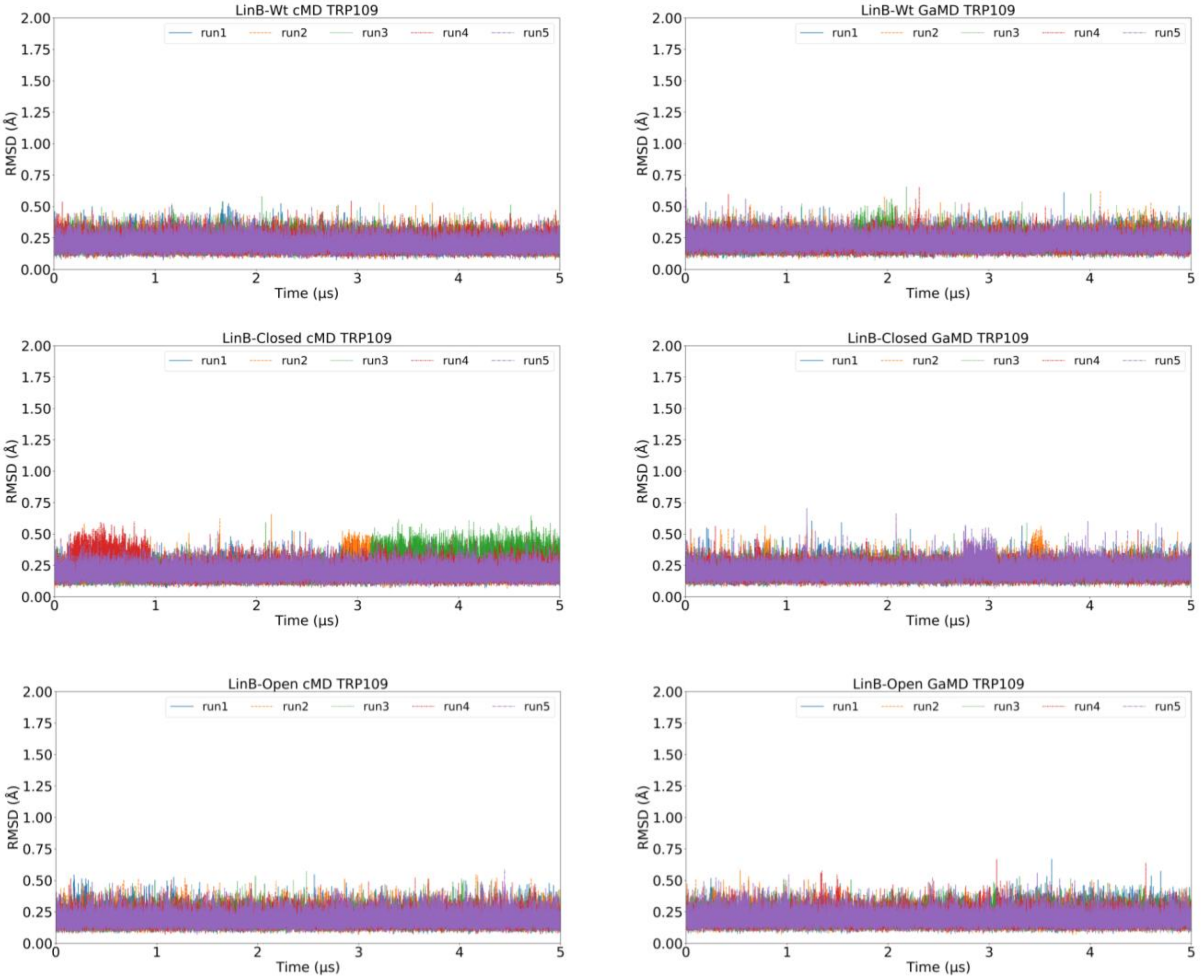

**Figure S9.** RMSD of catalytic residues TRP109 in all variants of LinB, to investigate its stability in the cMD and GaMD.

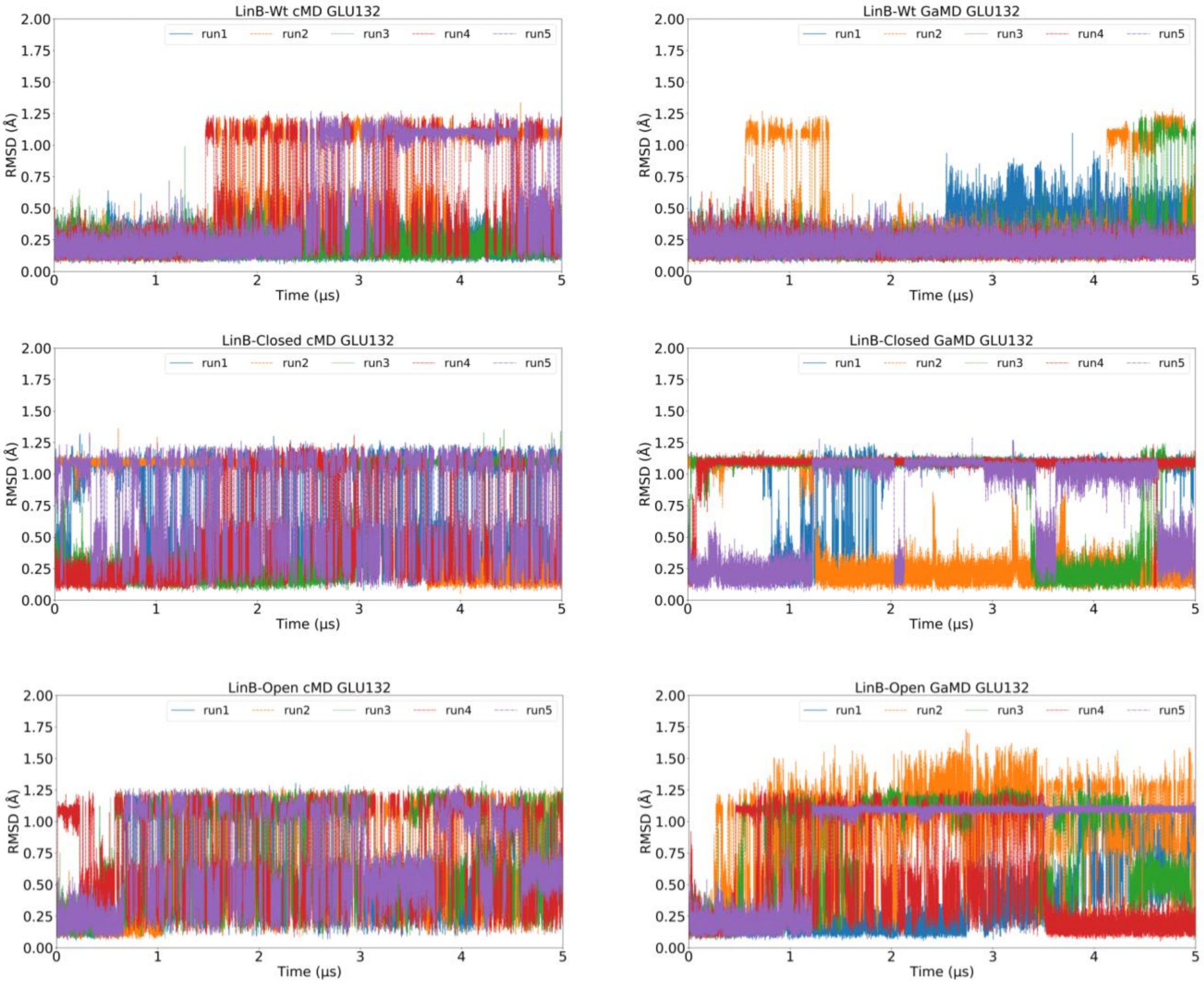

**Figure S10.** RMSD of catalytic residues GLU132 in all variants of LinB, to investigate its stability in the cMD and GaMD.

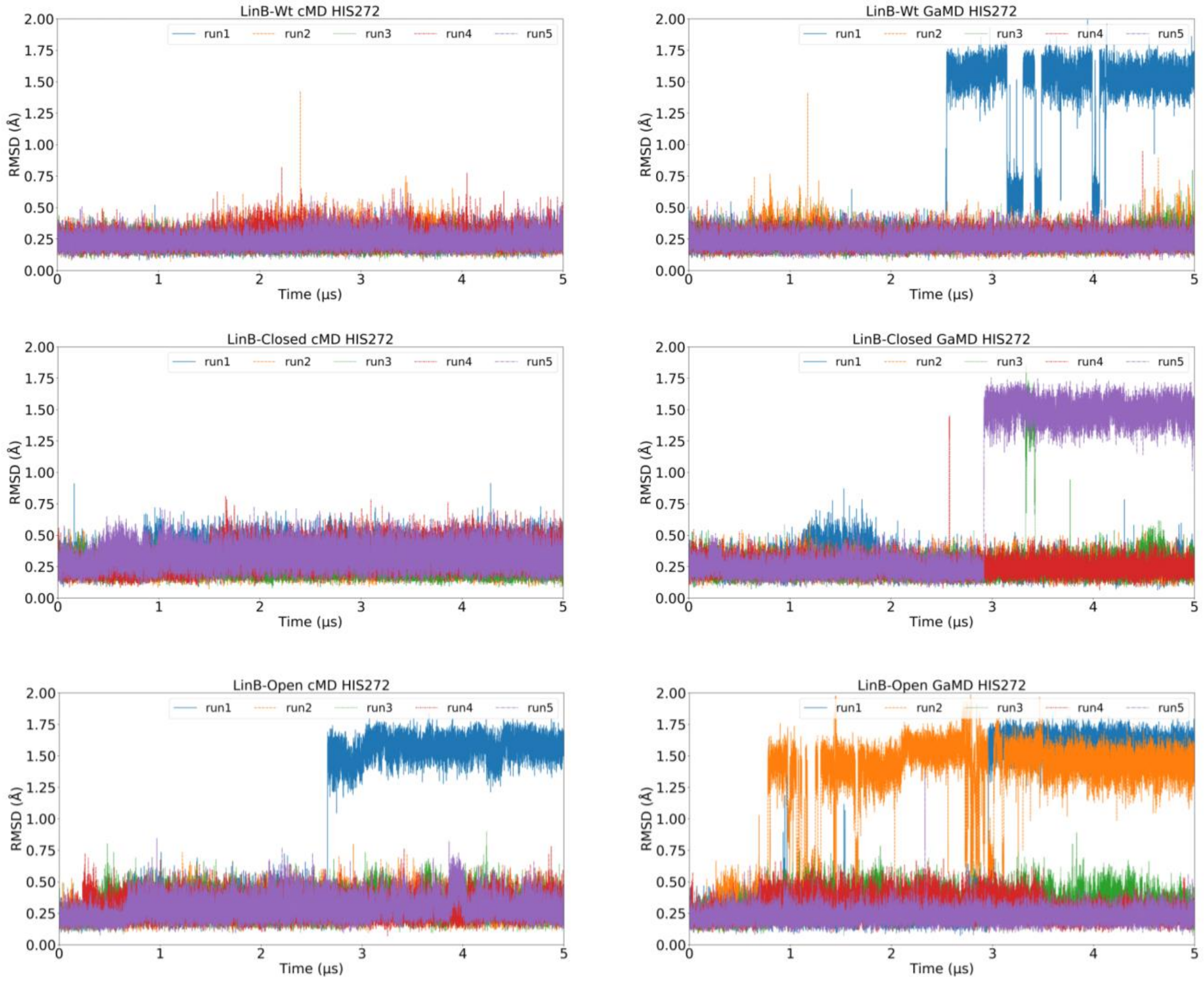

**Figure S11.** RMSD of catalytic residues HIS272 in all variants of LinB, to investigate its stability in the cMD and GaMD.

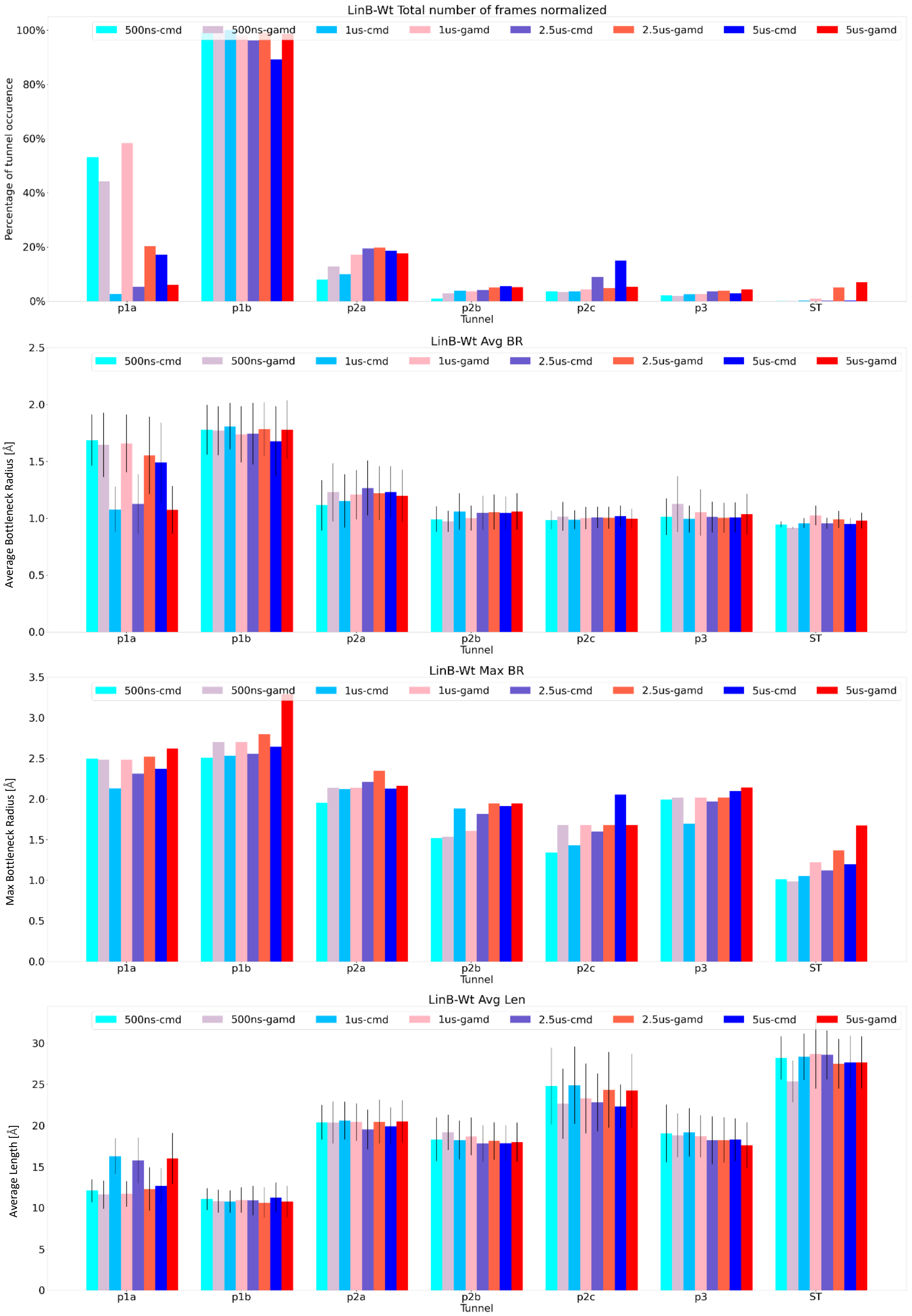

**Figure S12.** TransportTools results of LinB-Wt, comparison between cMD and reweighted GaMD: bar plots for total number of frames, average bottleneck radius (BR), average length and maximum BR for LinB-Wt obtained from corresponding fractions of 5, 2.5, 1 µs and 500 ns of cMD and GaMD simulations.

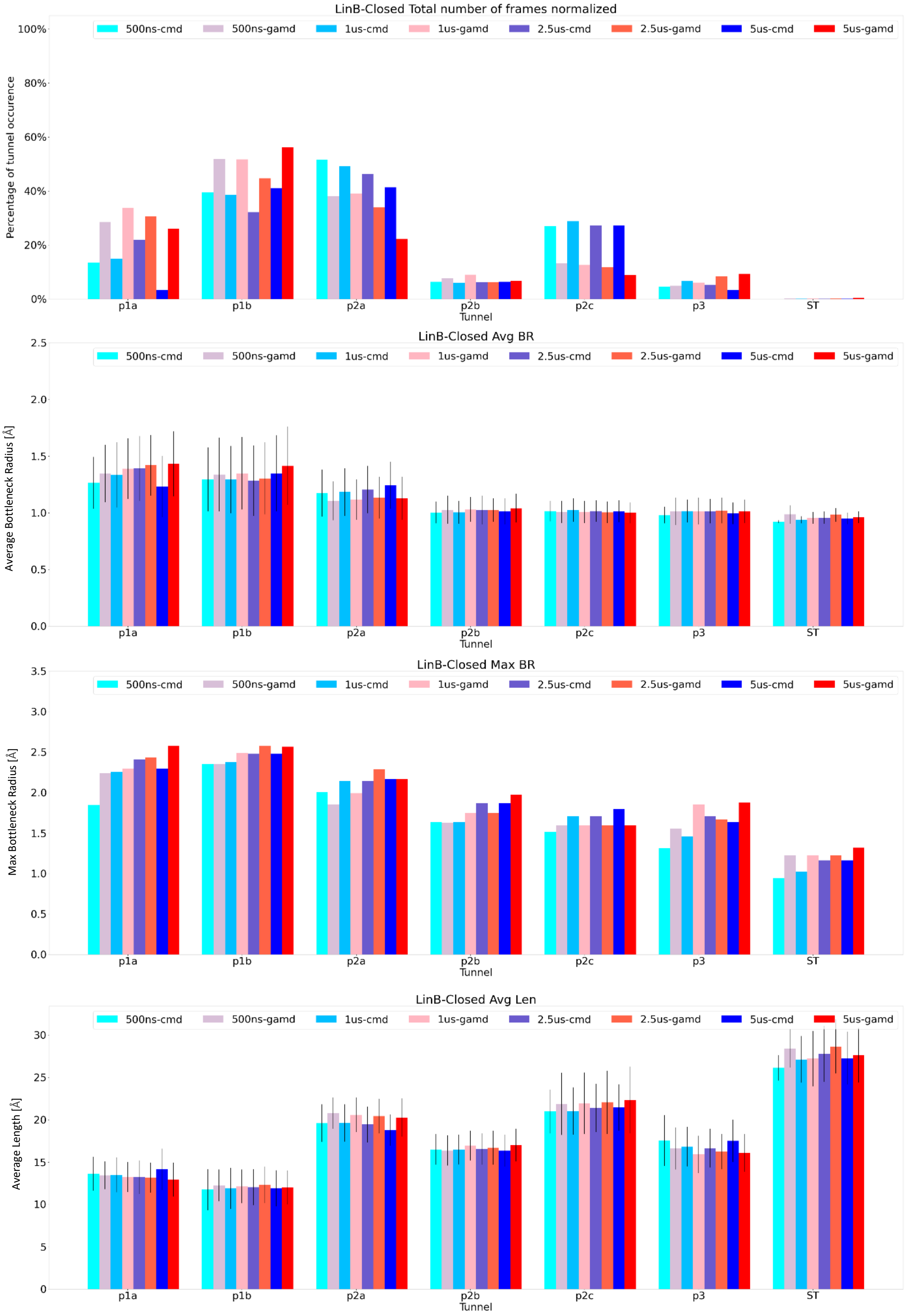

**Figure S13.**TransportTools results of LinB-Closed, comparison between cMD and reweighted GaMD: bar plots for total number of frames, Average bottleneck radius (BR), average length and maximum BR for LinB-Closed mutant obtained from corresponding fractions of 5, 2.5, 1 μs and 500 ns of cMD and GaMD simulations.

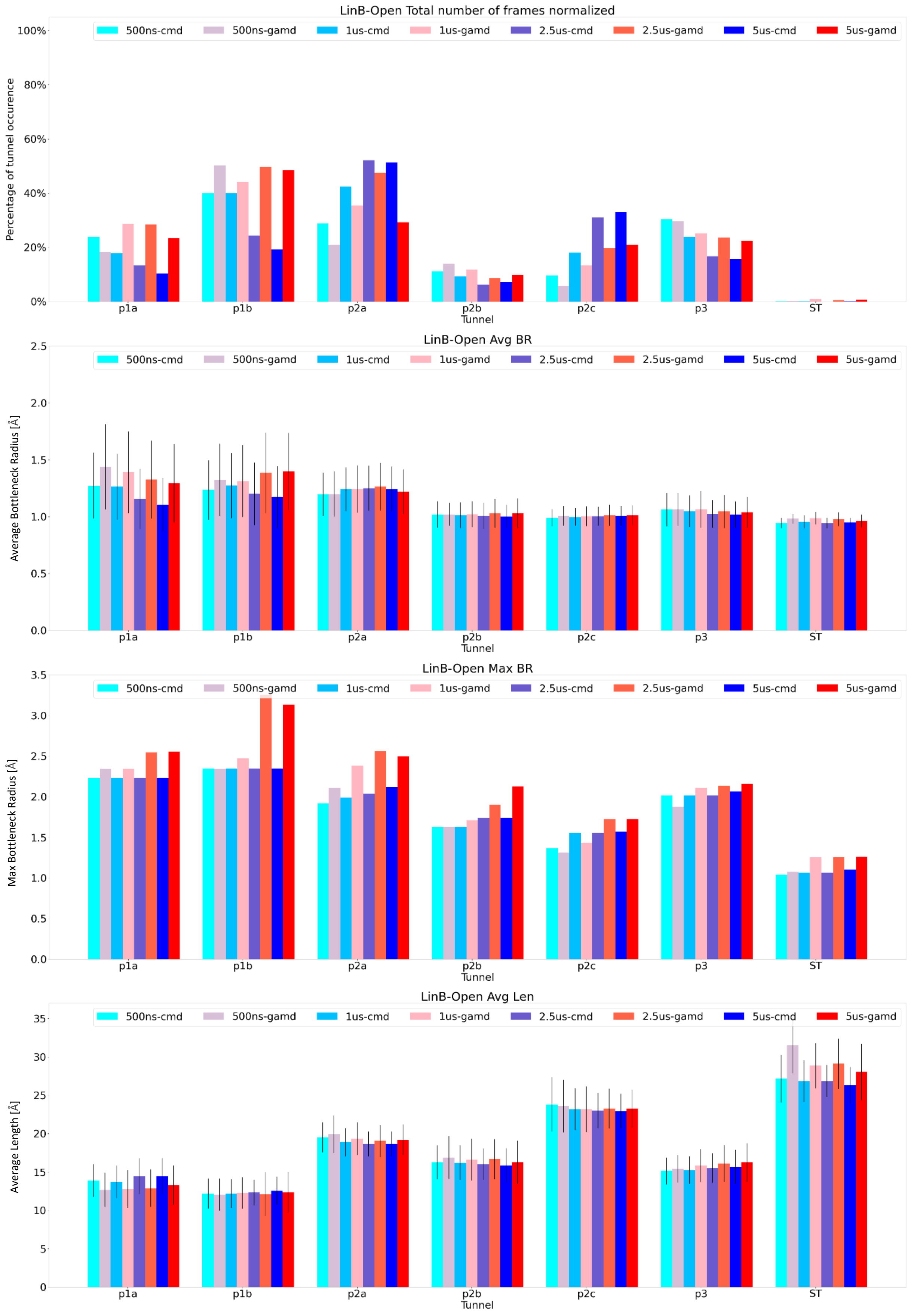

**Figure S14.** TransportTools results of LinB-Open, comparison between cMD and reweighted GaMD: bar plots for total number of frames, Average bottleneck radius (BR), average length and maximum BR for LinB-Open mutant obtained from corresponding fractions of 5, 2.5, 1 μs and 500 ns of cMD and GaMD simulations.

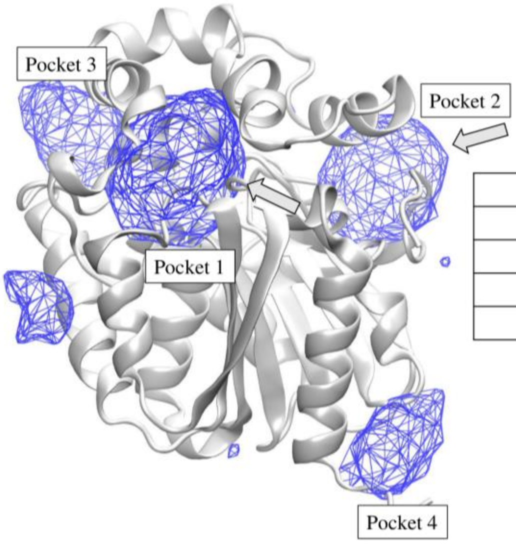

| Id | Centers | Score |
|---|---|---|
| 1 | [20.0, 35.3, 3.8] | 0.99 |
| 2 | [-2.0, 33.3, 1.8] | 0.92 |
| 3 | [20.0, 49.3, 13.8] | 0.77 |
| 4 | [-0.0, 11.3, 11.8] | 0.67 |

**Figure S15.** *Deepsite* tool prediction of viable druggable protein-ligand-binding sites using machine learning approach. Top two viable binding site represents p1 tunnel pocket as pocket1 and ST pocket as pocket2.

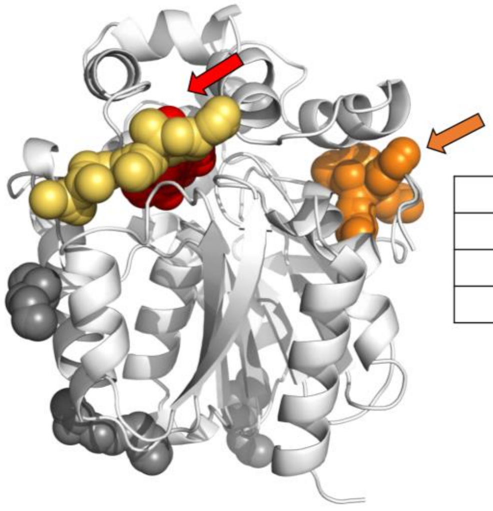

| Pocket Number | Probability |
|---|---|
| 1 | 83.61% |
| 2 | 21.61% |
| 3 | 16.79% |

**Figure S16.** *PASSer 2.0* tool prediction results for top three pockets as viable allosteric or cryptic site to bind ligand. Here, p1 tunnel pocket is shown in red colored spheres and arrow and ST pocket is shown as orange colored spheres and arrow.

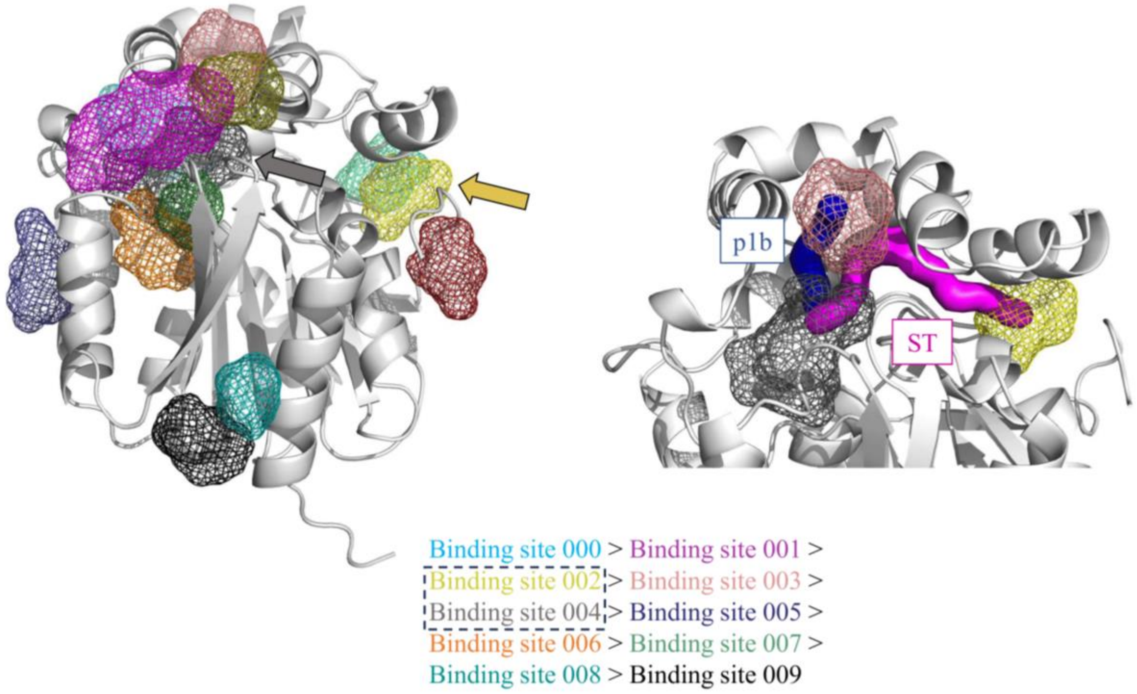

**Figure S17.** *FTMove* tool prediction result of transient binding sites on conformers of protein. This tool predicts cryptic allosteric site in the protein and probability is depicted by color in the figure. Here we can also see, ST pocket (yellow color - Binding site 002) is predicted as high probable cryptic site along with active site pocket (dark grey color - Binding site 004) (left). The ST connects the two cryptic pockets (Binding site 002 - 004) and also we can see main p1b tunnel connecting p1b tunnel pocket (salmon color - Binding site 003) and active site (Binding site 004) pocket (right).

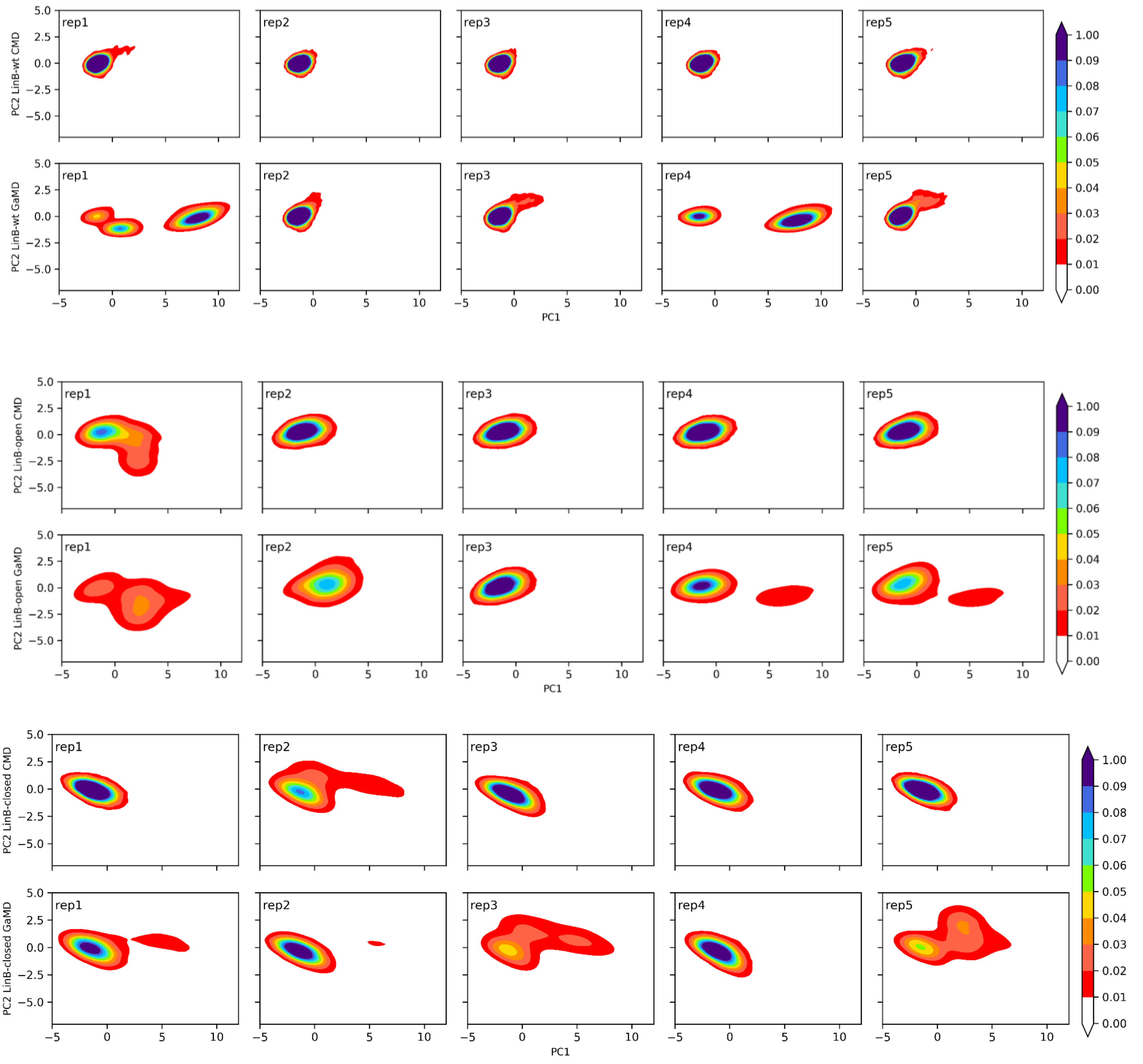

**Figure S18.** Distance based PCA of LinB-Wt, LinB-Closed and LinB-Open mutant, showing comparison between cMD and GaMD.

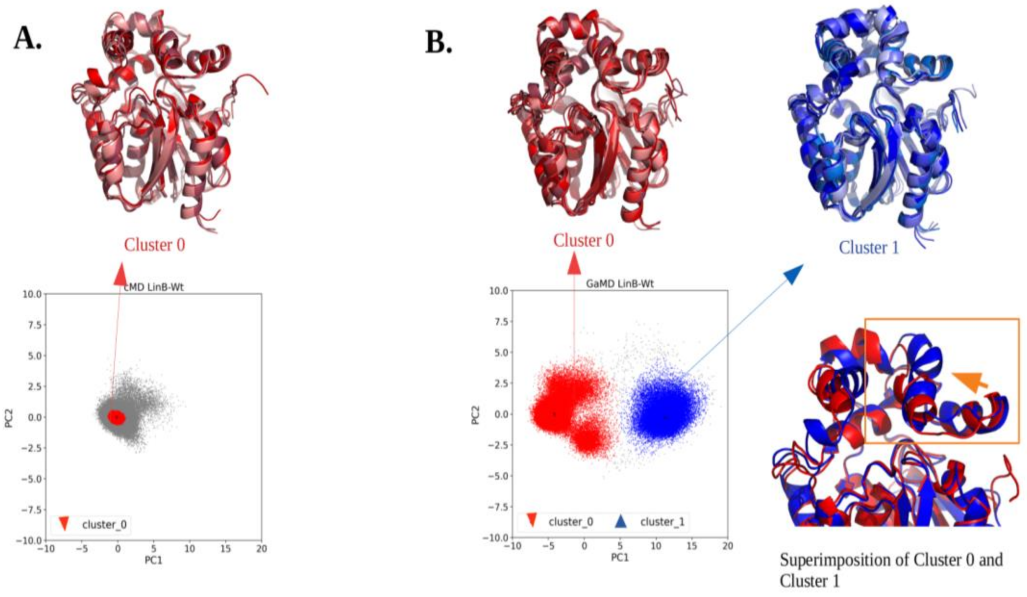

**Figure S19**. HDBscan Cluster analysis: LinB-Wt, (A) Cluster 0 with all representative frames from LinB-Wt cMD, (B) Cluster 0, Cluster 1 and overlap of Cluster 0 and Cluster 1 showing different conformations of representative structure of LinB-Wt captured by GaMD,

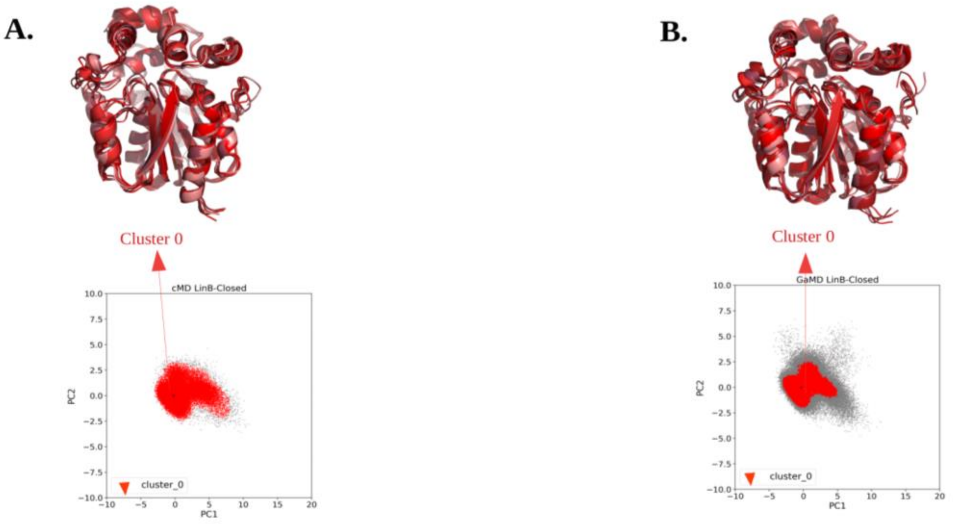

**Figure S20.** HDBscan Cluster analysis: LinB-Closed, (A) Cluster 0 with all representative frames of LinB-Closed mutant and (B) Cluster 0 with representative structures from LinB-Closed mutant,

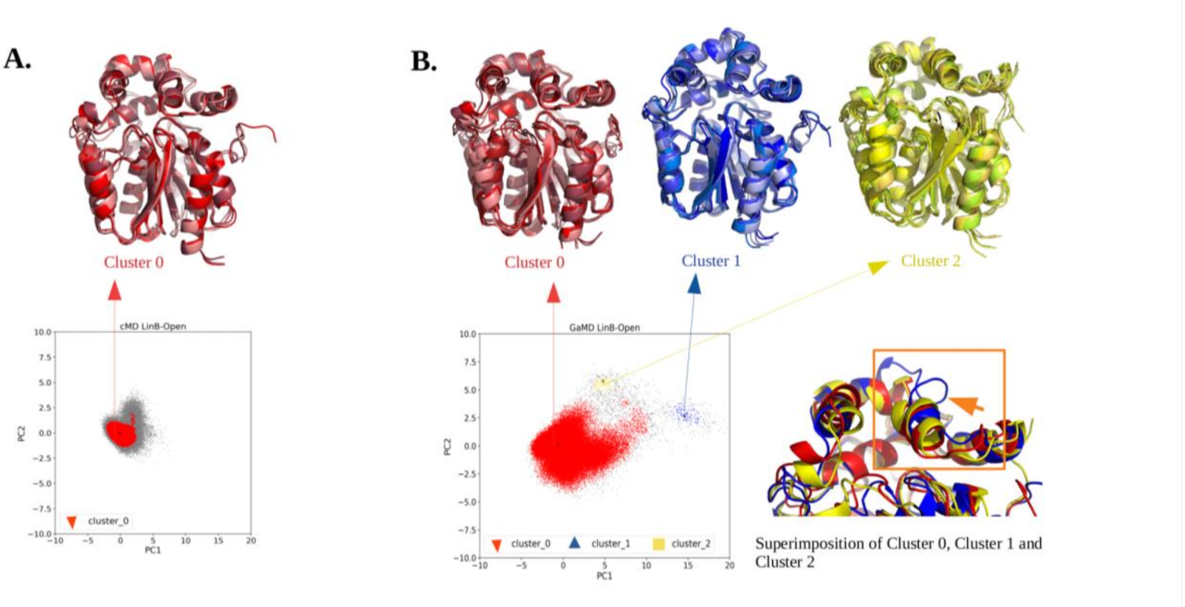

**Figure S21.** HDBscan cluster analysis: LinB-Open (A) Cluster 0 with all representative frames from LinB-Open mutant and (B) Cluster 0, Cluster 1 and Cluster 2 shown with all representative frames from LinB-Open mutant, superposition of all three clusters shown by orange box, where we can visualize different states of enzymes captured by GaMD method.

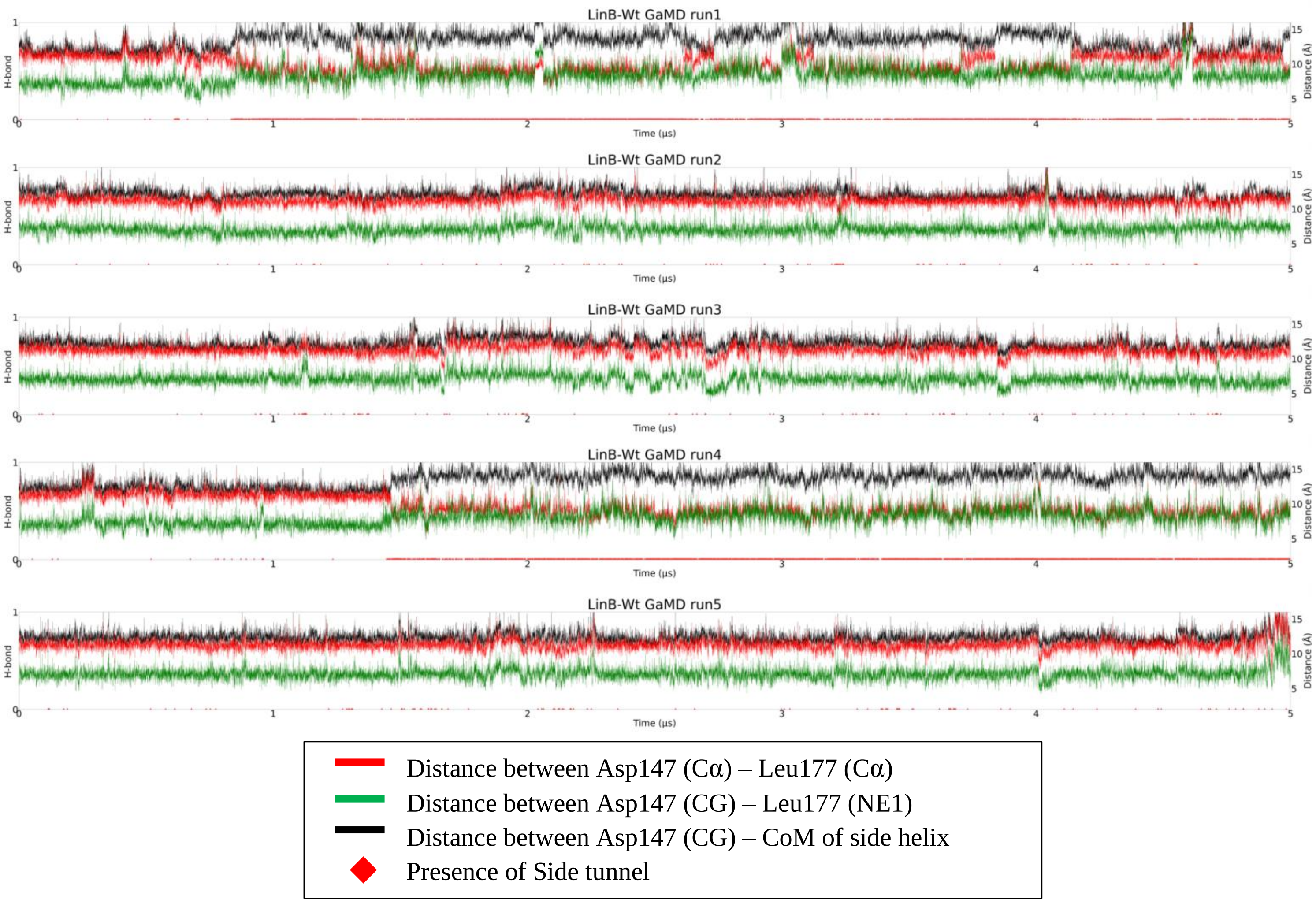

**Figure S22.** Mechanism of side tunnel (ST) opening in LinB-Wt using GaMD method. Here, Because of Leu177, it cannot make hydrogen bond with Asp147 and hence the hydrogen bonds presence is at 0 and we will observe the opening of ST in absence of h-bond. The movement of side helix was investigated by distance analysis between atoms (Cα) of Leu, which is present on side helix in respective to Asp which is shown in red color and distance between same residue but different atom Asp (CG) and Leu (NE1) to analyze the side chain behavior is shown by green color. Distance between Asp side chain atom (CG) and center of mass (CoM) of side helix to understand the mechanism of opening of ST is shown in black color and finally opening of ST are shown in red diamonds in the figure. In result, we observed and mostly can be seen in run1 and run4 that the distance increases between Asp147 (CG) and CoM of side helix also distance increases between Asp147 – Leu177 (both backbone Cα and side chain atoms) and the ST opens. We can observe around 1 μs and after that in run1 and run4 of GaMD, there is increase of red diamond appearance, which indicated that movement of side helix farther from the protein is required for the opening of ST in LinB-Wt.

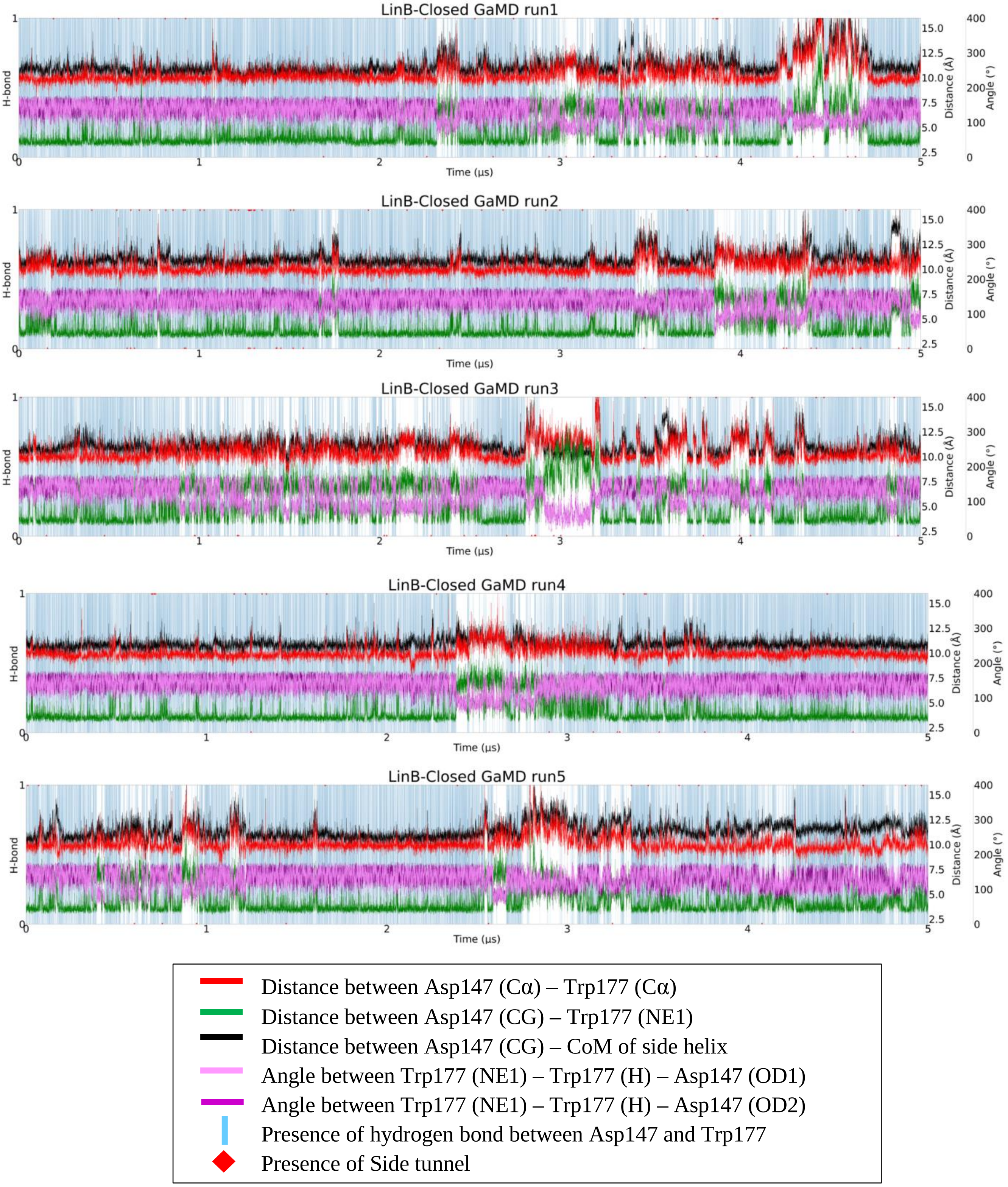

**Figure S23.** Mechanism of side tunnel opening in LinB-Closed mutant using GaMD method. Here because of the mutation at Leu177 to Trp177, it forms hydrogen bond with Asp147. Analogously to previous plot, the presence and absence of hydrogen bond is shown by 0 and 1 by blue vertical line. To understand the mechanism of ST opening in Closed mutant, we have analyzed distance between Trp177 backbone atom (Cα), side chain (CG) which is on the side helix to Asp147 (Cα) and side chain atom (NE1) by red and green color. Further we have analyzed the movement of side helix with respective of protein by analyzing distance between Asp147 and CoM of side helix by black color. To understand the movement of phi and psi angle, the angle between the Trp and Asp were analyzed by pink and magenta color, and presence or opening of ST is shown by red diamond. In result, we can observe that the breakage of hydrogen bond is accompanied by the opening of ST and during such event, the distance increases between the Asp and Trp as well as the distance increases between CoM of side helix and Asp147. Also we can observe the fluctuation in psi and phi angle during the event of hydrogen bond breakage. So, in mutant the ST opens with the breakage of H-bond and movement of side helix farther from the protein.

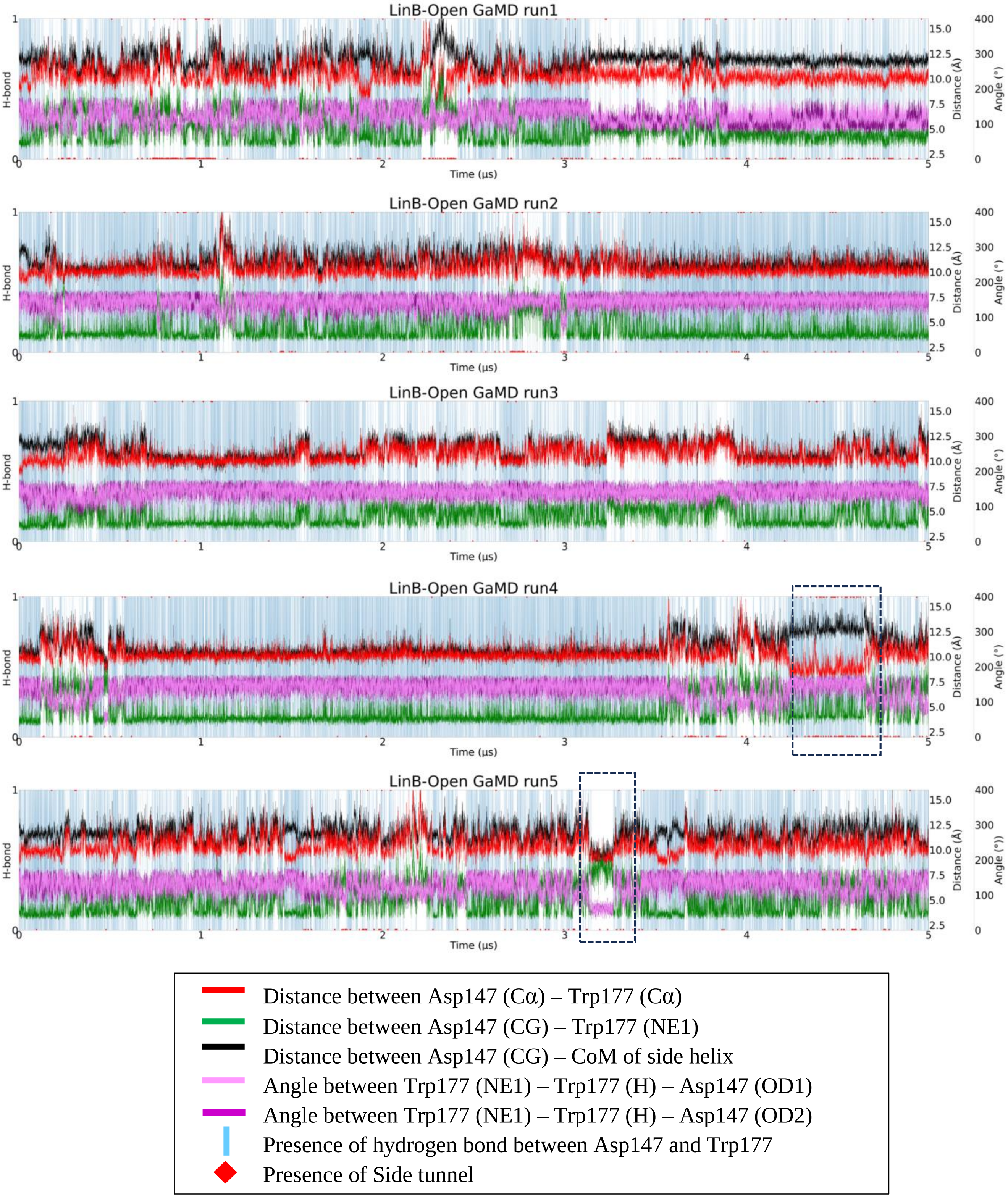

**Figure S24.** Mechanism of side tunnel opening in LinB-Open mutant using GaMD method. Analogously to Figure S23. Some interesting cases indicated by dashed lines, in few cases, there is hydrogen bond formation and still opening of ST, we investigated that in these cases the hydrogen bond is formed by the residues but still the movement of side helix can be seen farther away from the protein (black color), and due to the movement of side helix away from the protein, it helps with the opening of the ST. In another case there is no hydrogen bond formation despite the residues Asp147 and Trp177 backbone are close enough, in this case the backbone is close enough but the side chain is far investigated by distance between Asp147 (CG) and Trp177 (NE1) in green color, hence no formation of hydrogen bond.
