## Supplementary File 2 for "Reinforcing Tunnel Network Exploration in Proteins Using Gaussian Accelerated Molecular Dynamics"

**Corresponding authors**

transport tunnel, molecular dynamics, haloalkane dehalogenase, enhanced sampling methods, mechanism

Caverdock energy profiles of individual 100 tunnels in all variants.

1. Upper Bound energy profile of p1b-Wt with Bromide ion ($Br^-$) ligand. The X-axis represents upper bound energy (kcal/mol) and Y-axis represents length of the trajectory [Å], along the disc of tunnel.

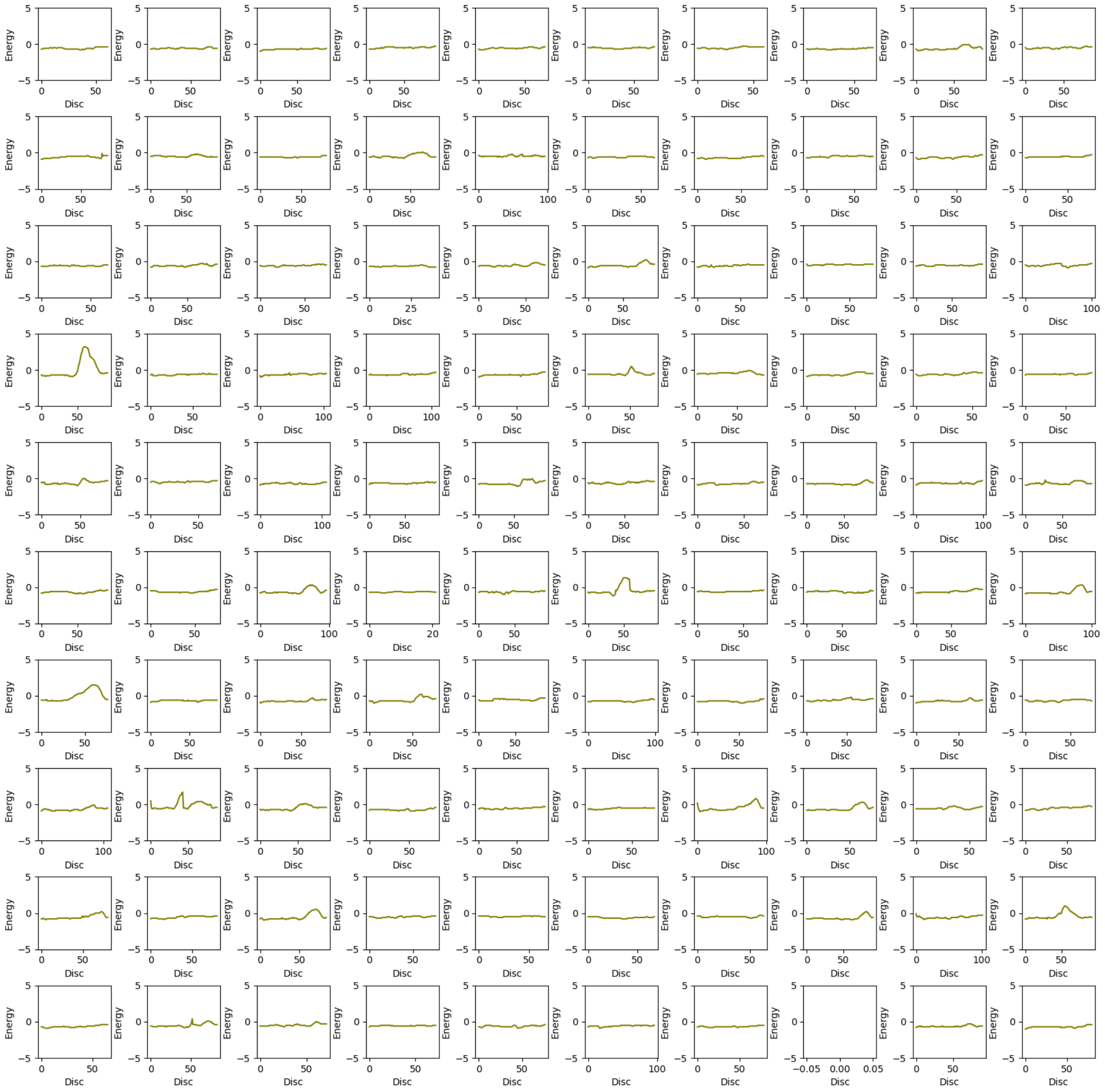

2. Upper Bound energy profile of p1b-Wt with 2-Bromoethanol (be) ligand. The X-axis represents upper bound energy (kcal/mol) and Y-axis represents length of the trajectory [Å], along the disc of tunnel.

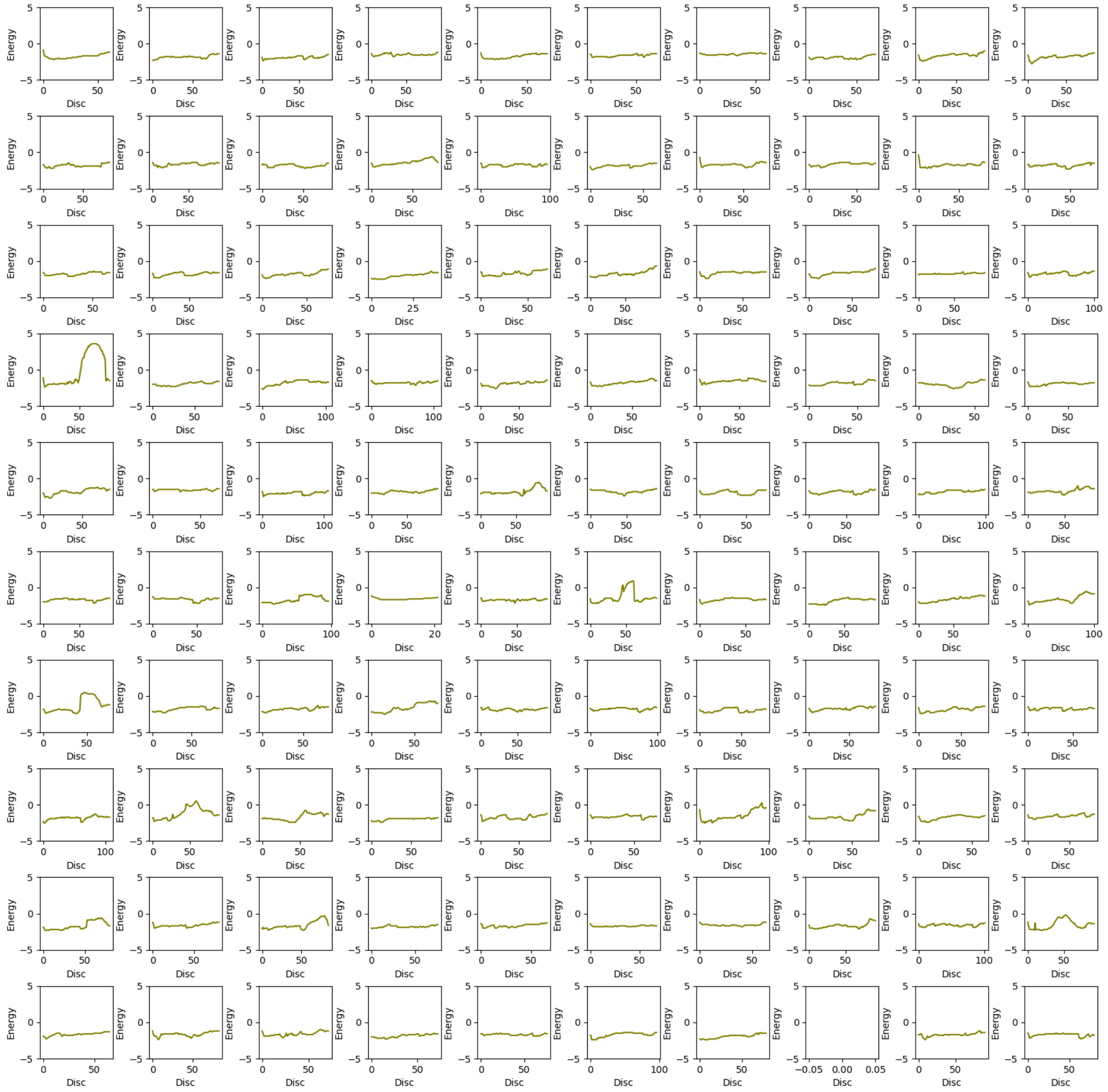

3. Upper Bound energy profile of p1b-Wt with 1,2- Dibromoethane (dbe) ligand. The X-axis represents upper bound energy (kcal/mol) and Y-axis represents length of the trajectory [Å], along the disc of tunnel.

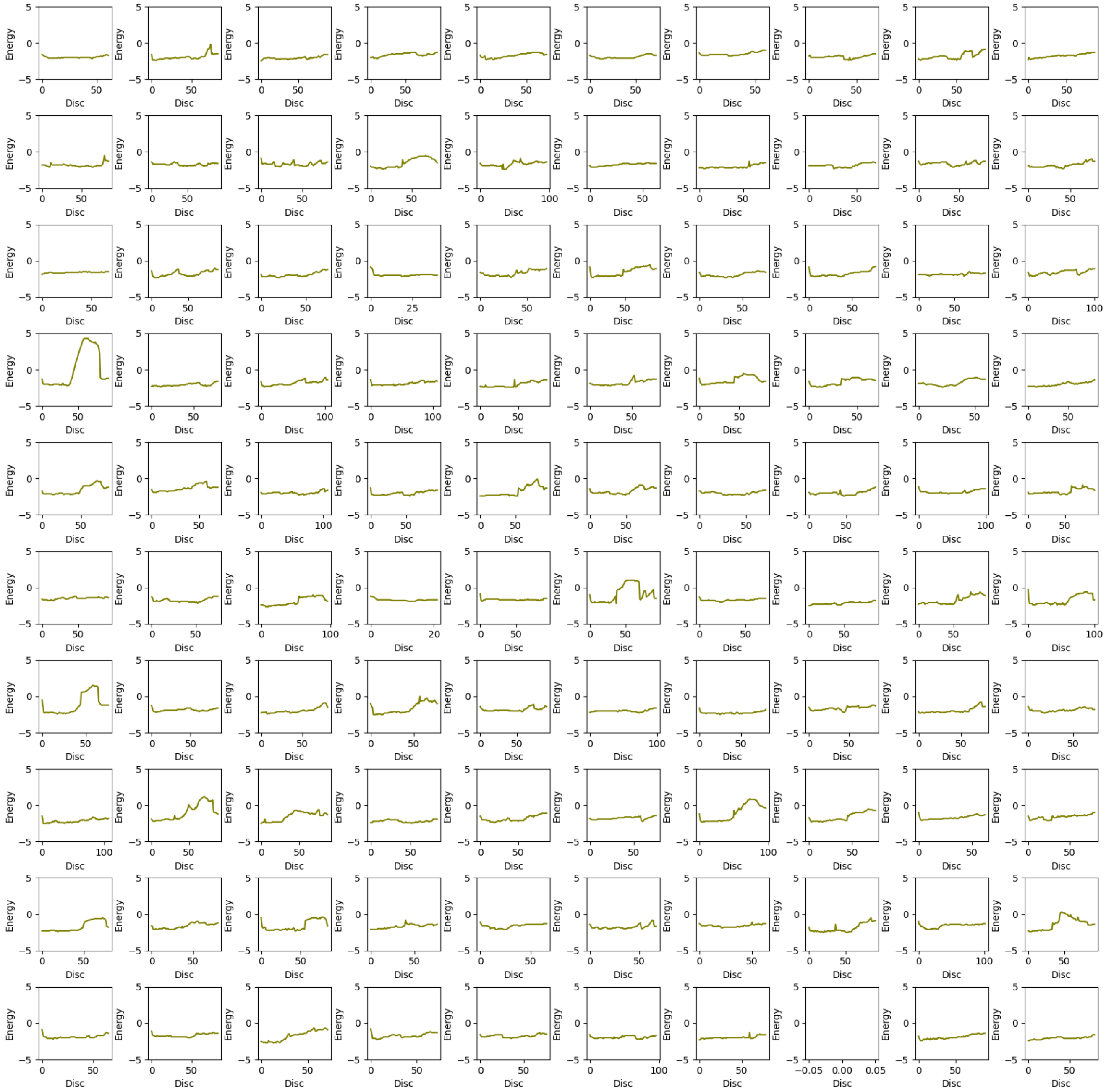

4. Upper Bound energy profile of p1b-Wt with Water ($H_2O$) ligand. The X-axis represents upper bound energy (kcal/mol) and Y-axis represents length of the trajectory [Å], along the disc of tunnel.

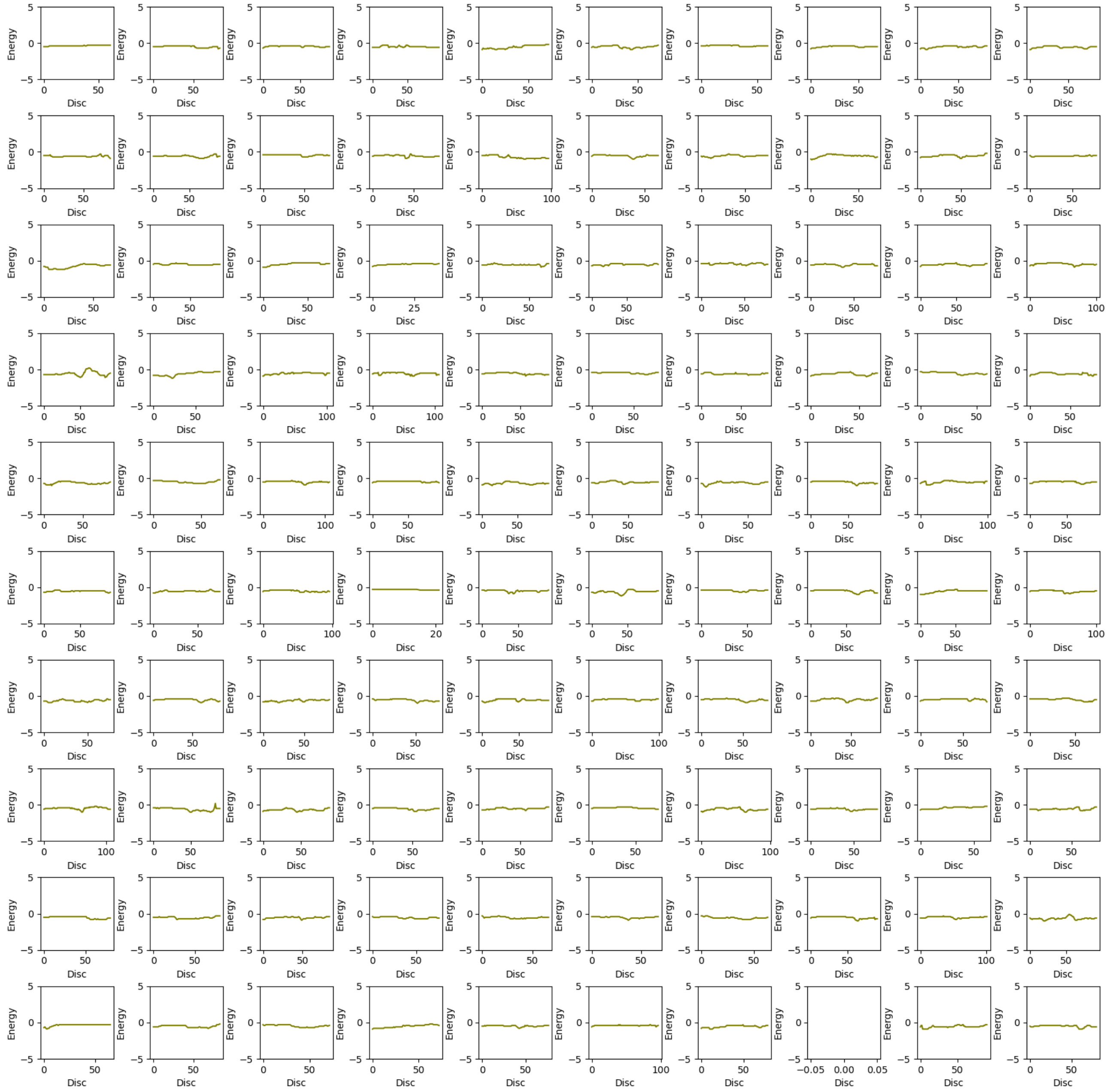

5. Upper Bound energy profile of p1b-Closed with Bromide ion ($Br^-$) ligand. The X-axis represents upper bound energy (kcal/mol) and Y-axis represents length of the trajectory [Å], along the disc of tunnel.

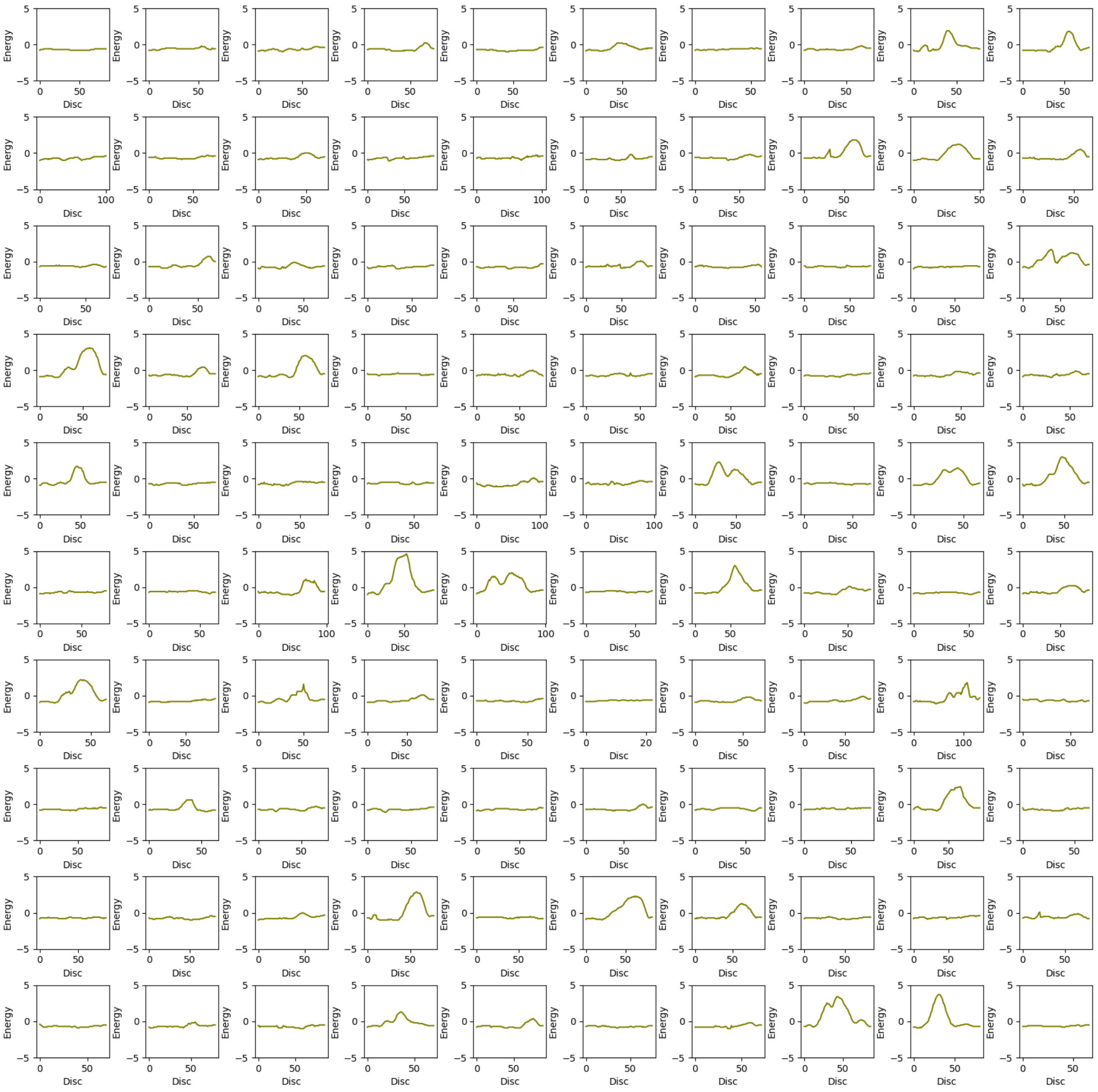

6. Upper Bound energy profile of p1b-Closed with 2-Bromoethanol (be) ligand. The X-axis represents upper bound energy (kcal/mol) and Y-axis represents length of the trajectory [Å], along the disc of tunnel.

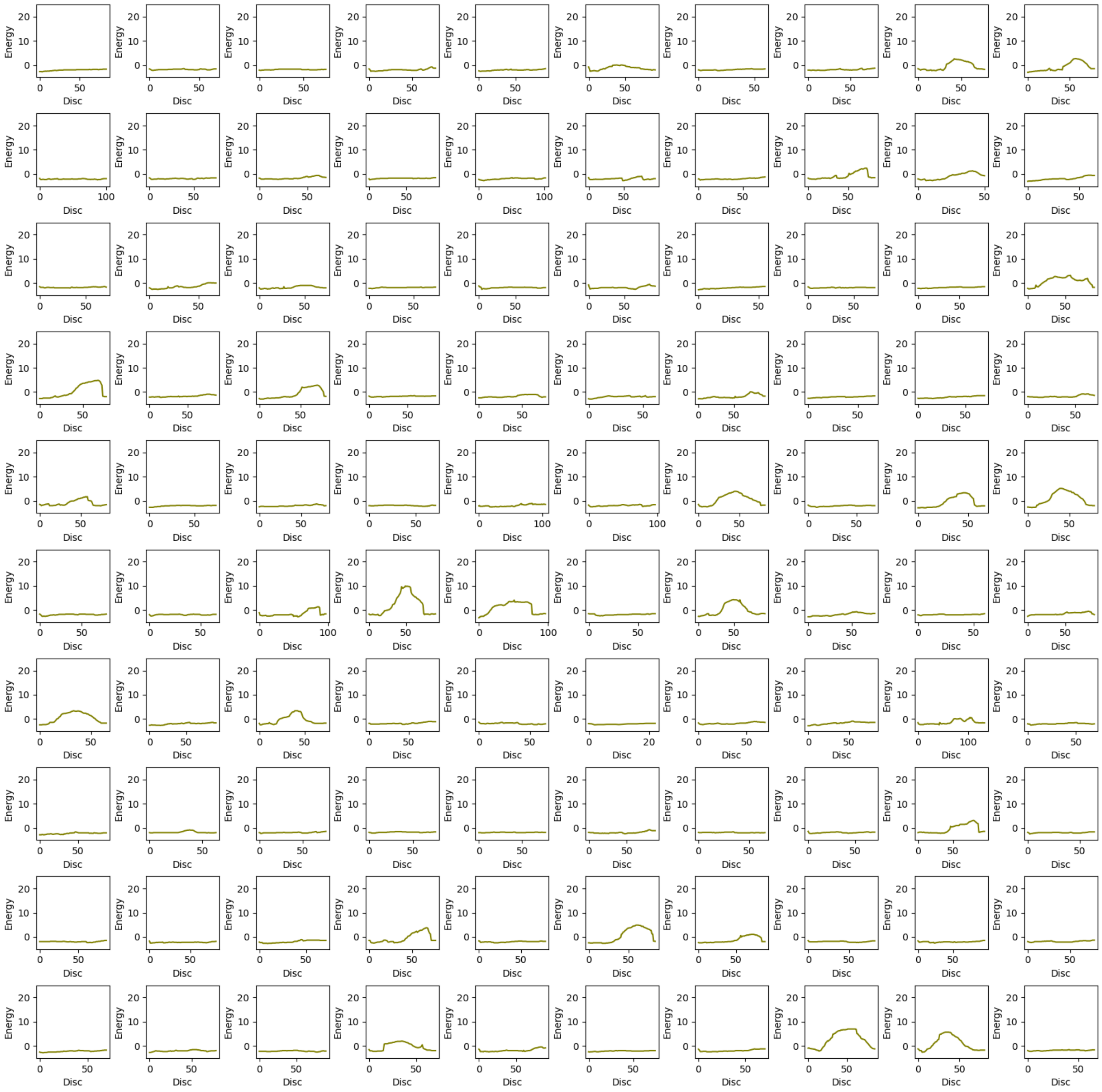

7. Upper Bound energy profile of p1b-Closed with 1,2- Dibromoethane (dbe) ligand. The X-axis represents upper bound energy (kcal/mol) and Y-axis represents length of the trajectory [Å], along the disc of tunnel.

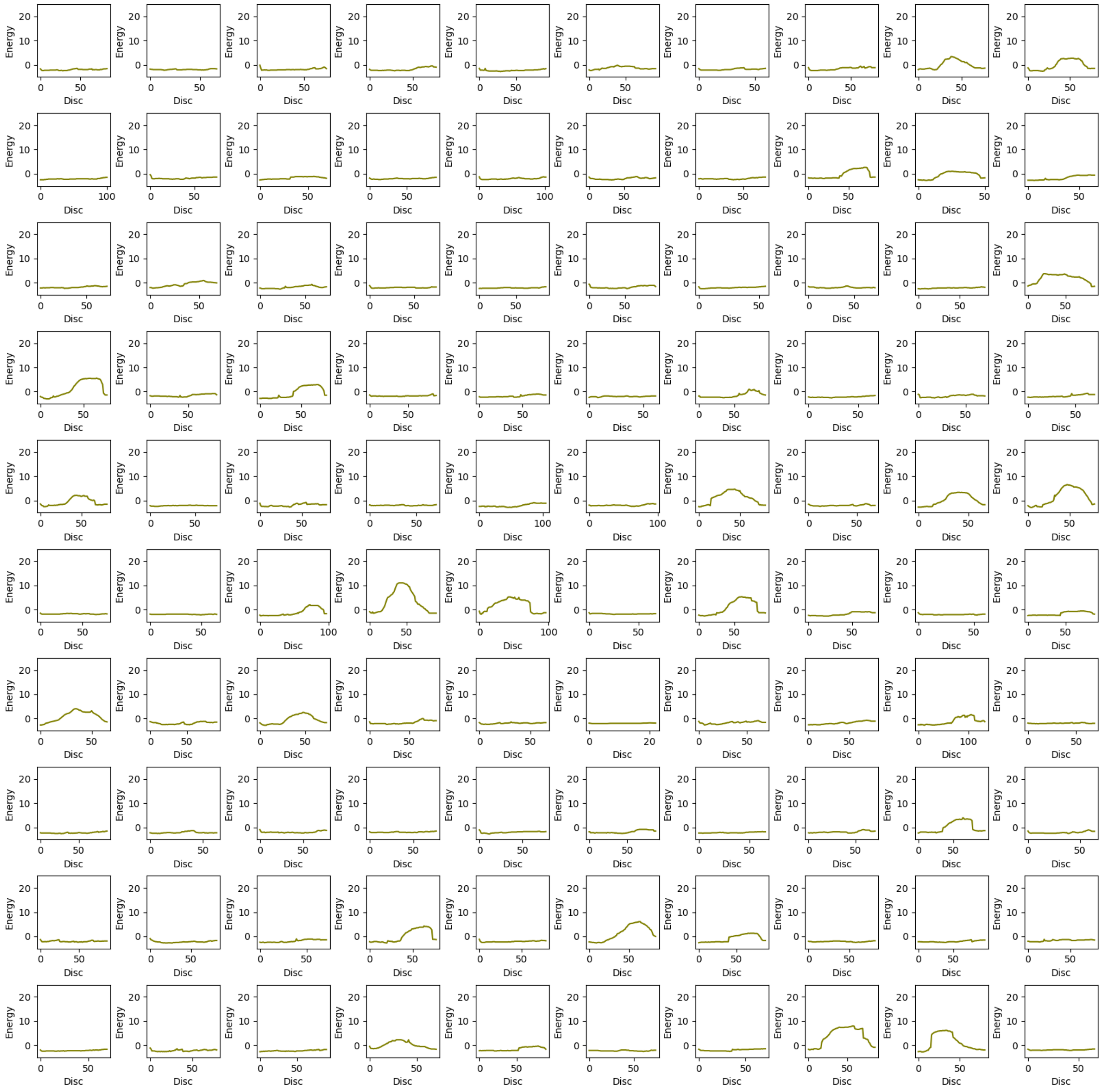

8. Upper Bound energy profile of p1b-Closed with Water ($H_2O$) ligand. The X-axis represents upper bound energy (kcal/mol) and Y-axis represents length of the trajectory [Å], along the disc of tunnel.

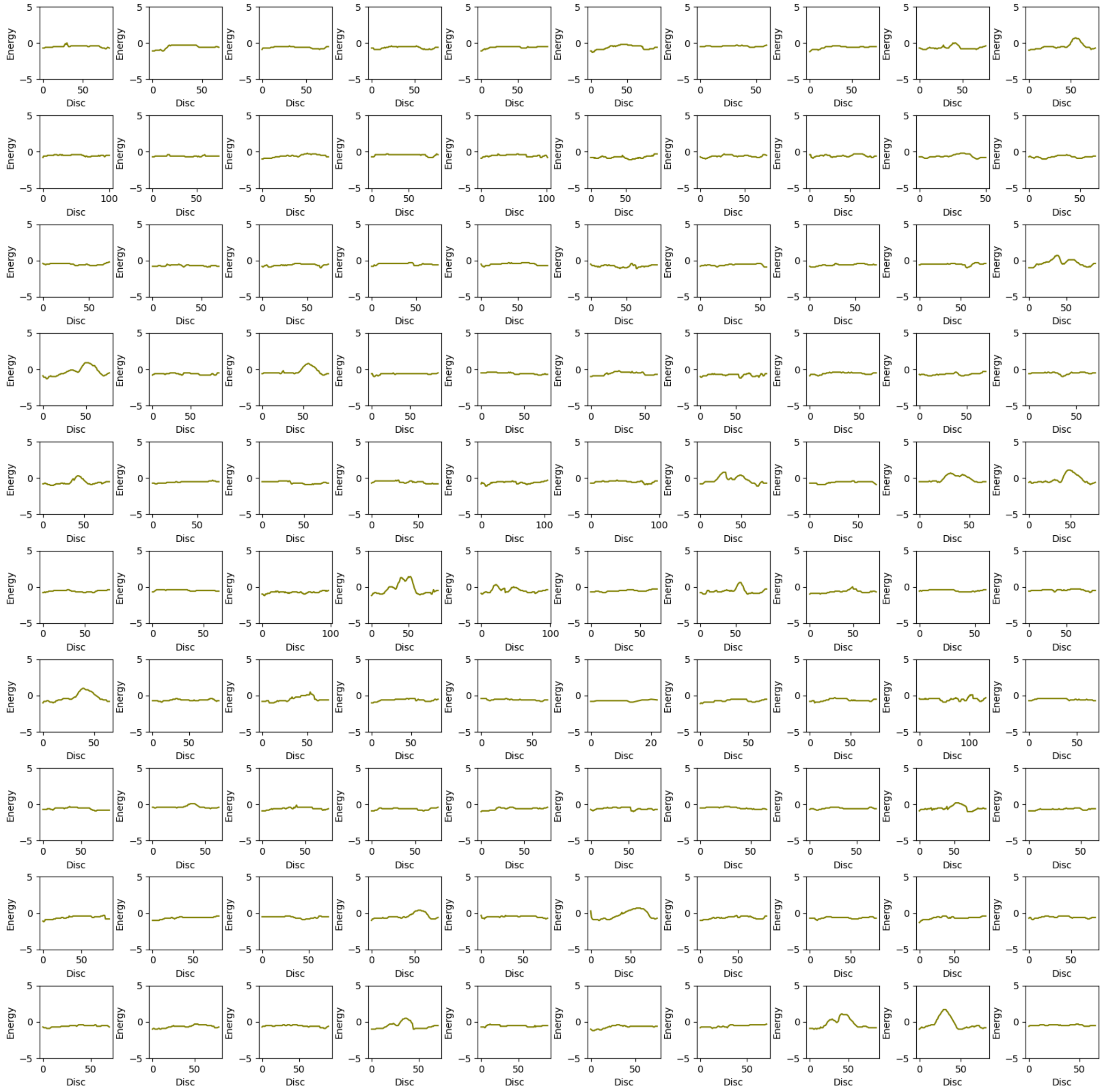

9. Upper Bound energy profile of p1b-Open with Bromide ion ($Br^-$) ligand. The X-axis represents upper bound energy (kcal/mol) and Y-axis represents length of the trajectory [Å], along the disc of tunnel.

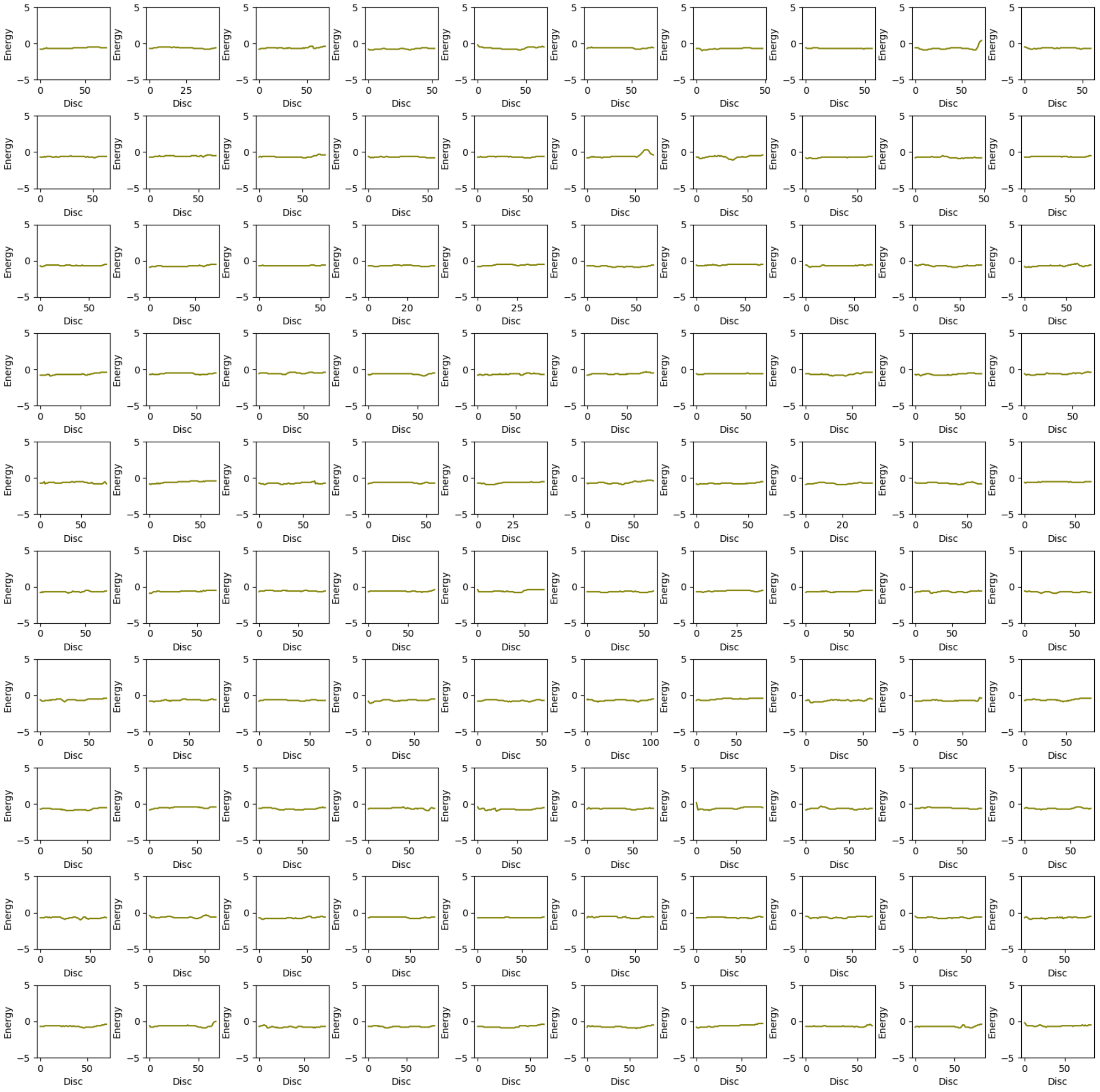

10. Upper Bound energy profile of p1b-Open with 2-Bromoethanol (be) ligand. The X-axis represents upper bound energy (kcal/mol) and Y-axis represents length of the trajectory [Å], along the disc of tunnel.

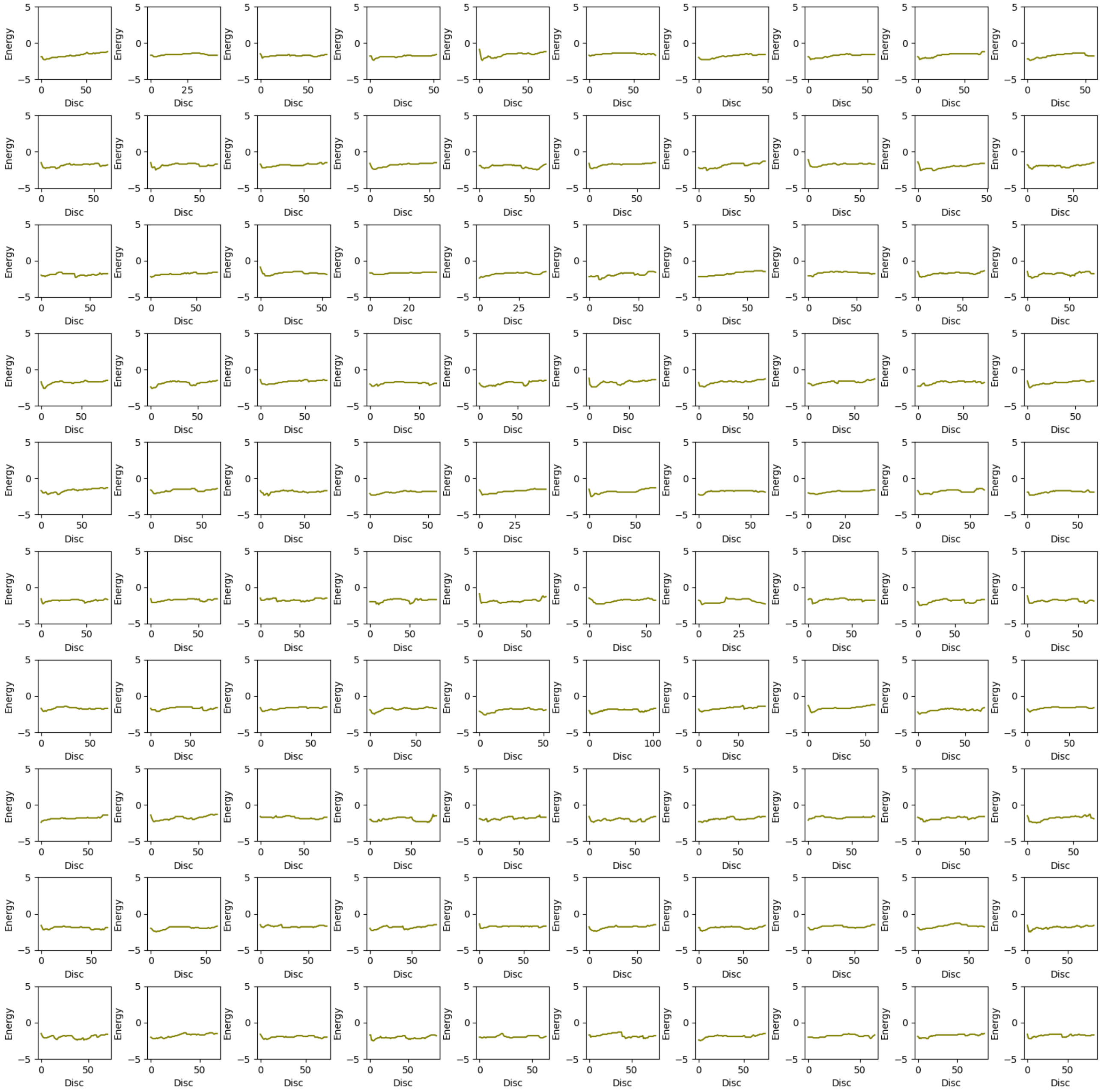

11. Upper Bound energy profile of p1b-Open with 1,2- Dibromoethane (dbe) ligand. The X-axis represents upper bound energy (kcal/mol) and Y-axis represents length of the trajectory [Å], along the disc of tunnel.

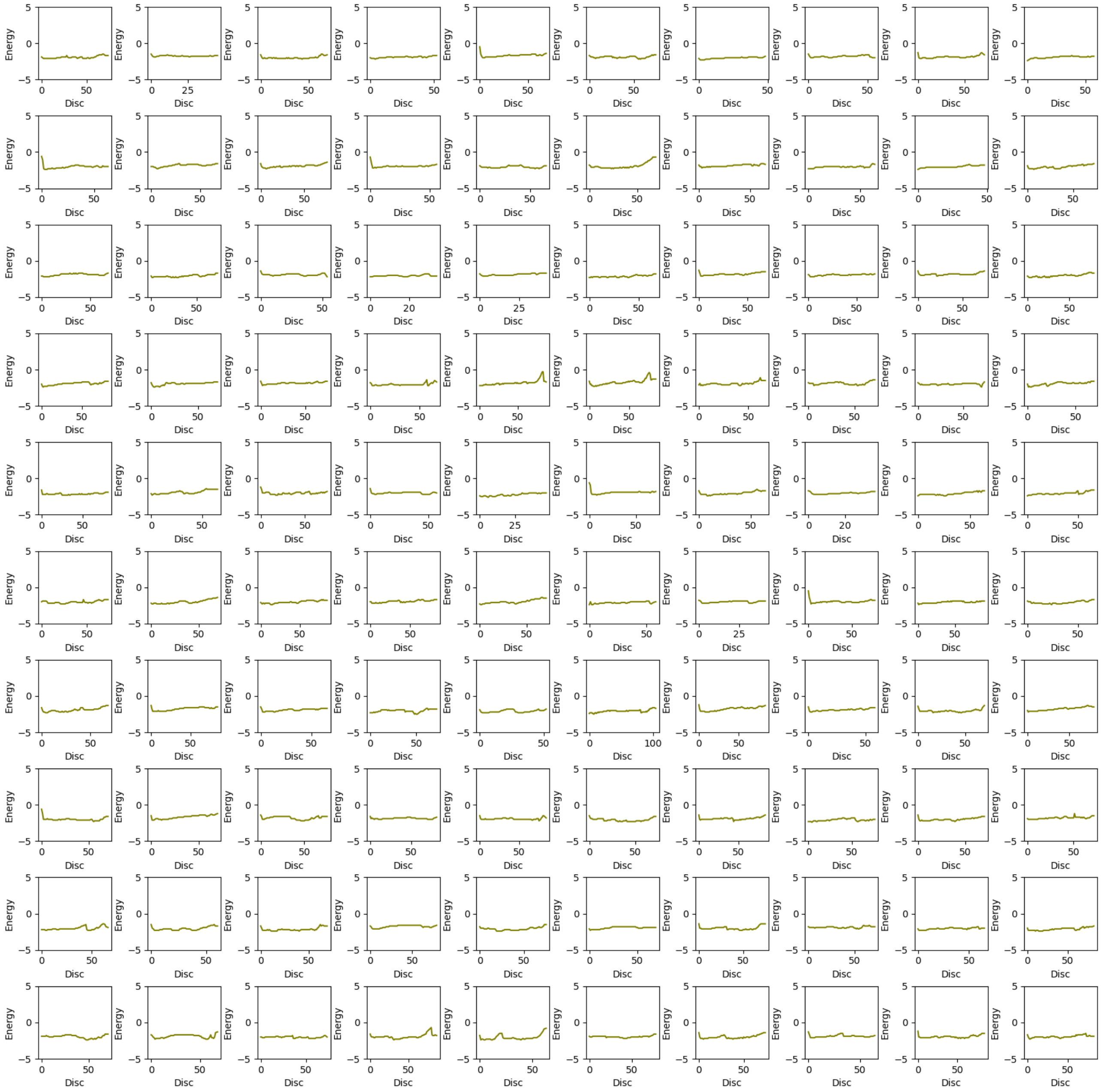

12. Upper Bound energy profile of p1b-Open with Water ($H_2O$) ligand. The X-axis represents upper bound energy (kcal/mol) and Y-axis represents length of the trajectory [Å], along the disc of tunnel.

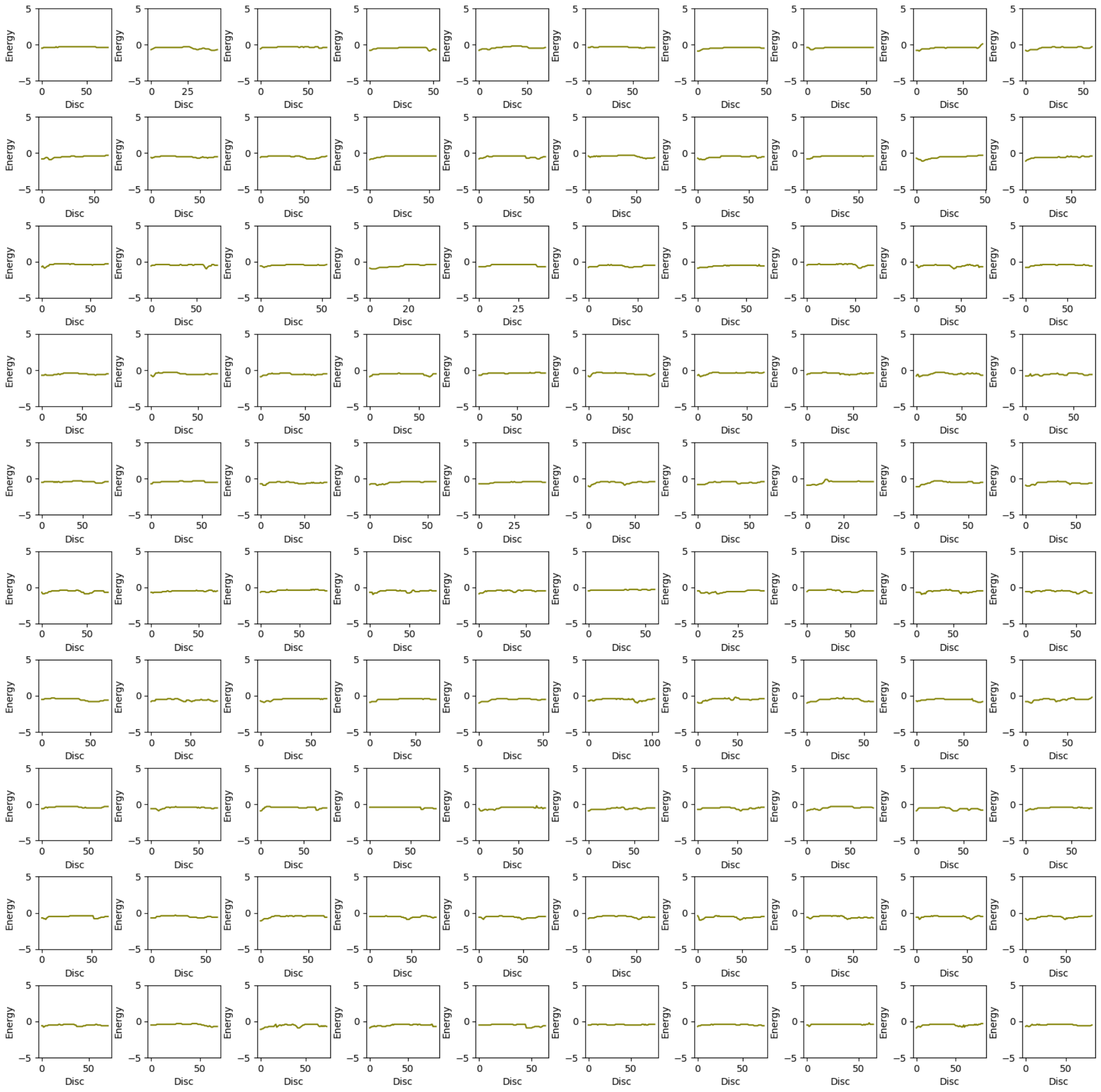

13. Upper Bound energy profile of p3-Wt with Bromide ion ($Br^-$) ligand. The X-axis represents upper bound energy (kcal/mol) and Y-axis represents length of the trajectory [Å], along the disc of tunnel.

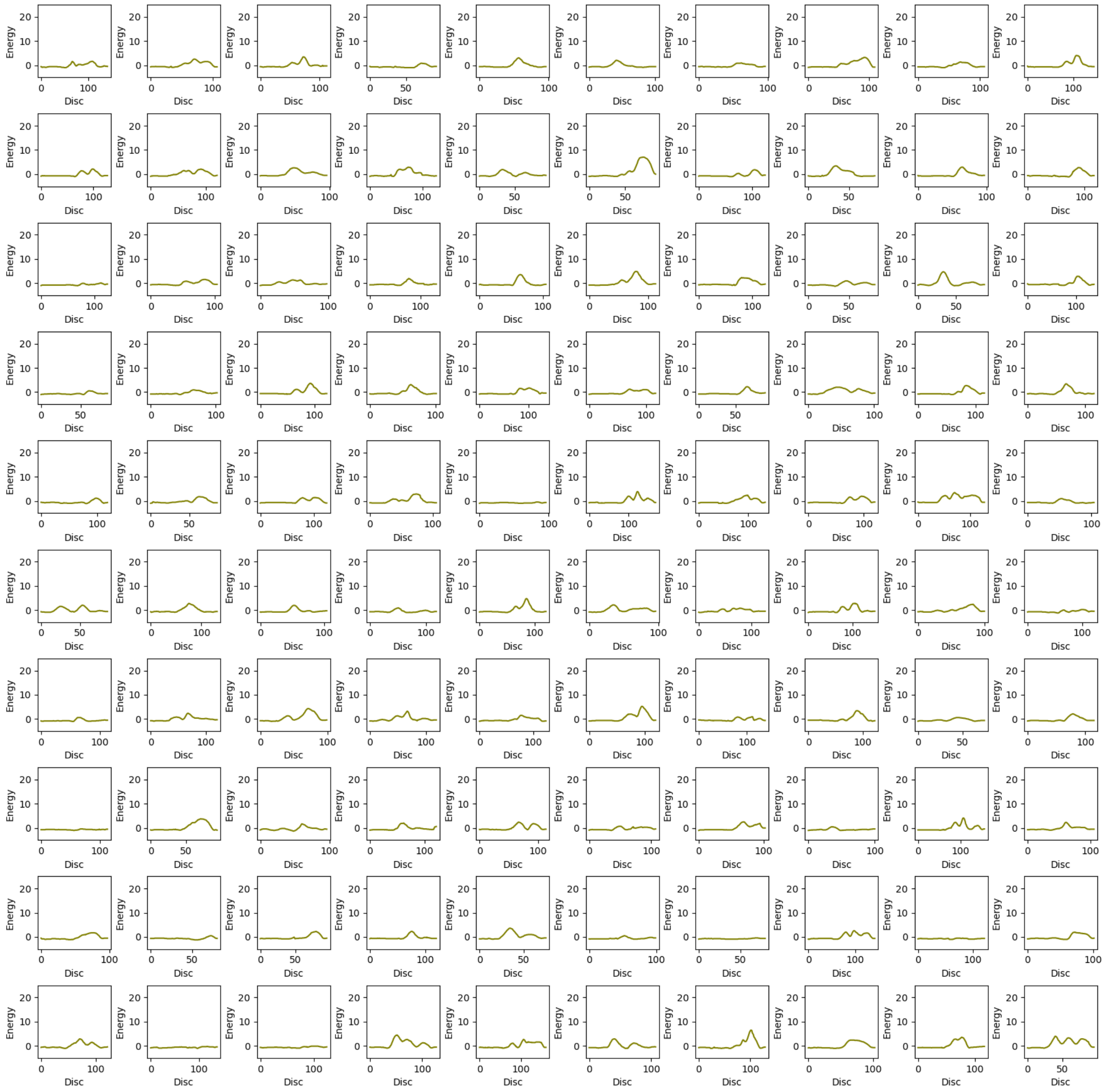

14. Upper Bound energy profile of p3-Wt with 2-Bromoethanol (be) ligand. The X-axis represents upper bound energy (kcal/mol) and Y-axis represents length of the trajectory [Å], along the disc of tunnel.

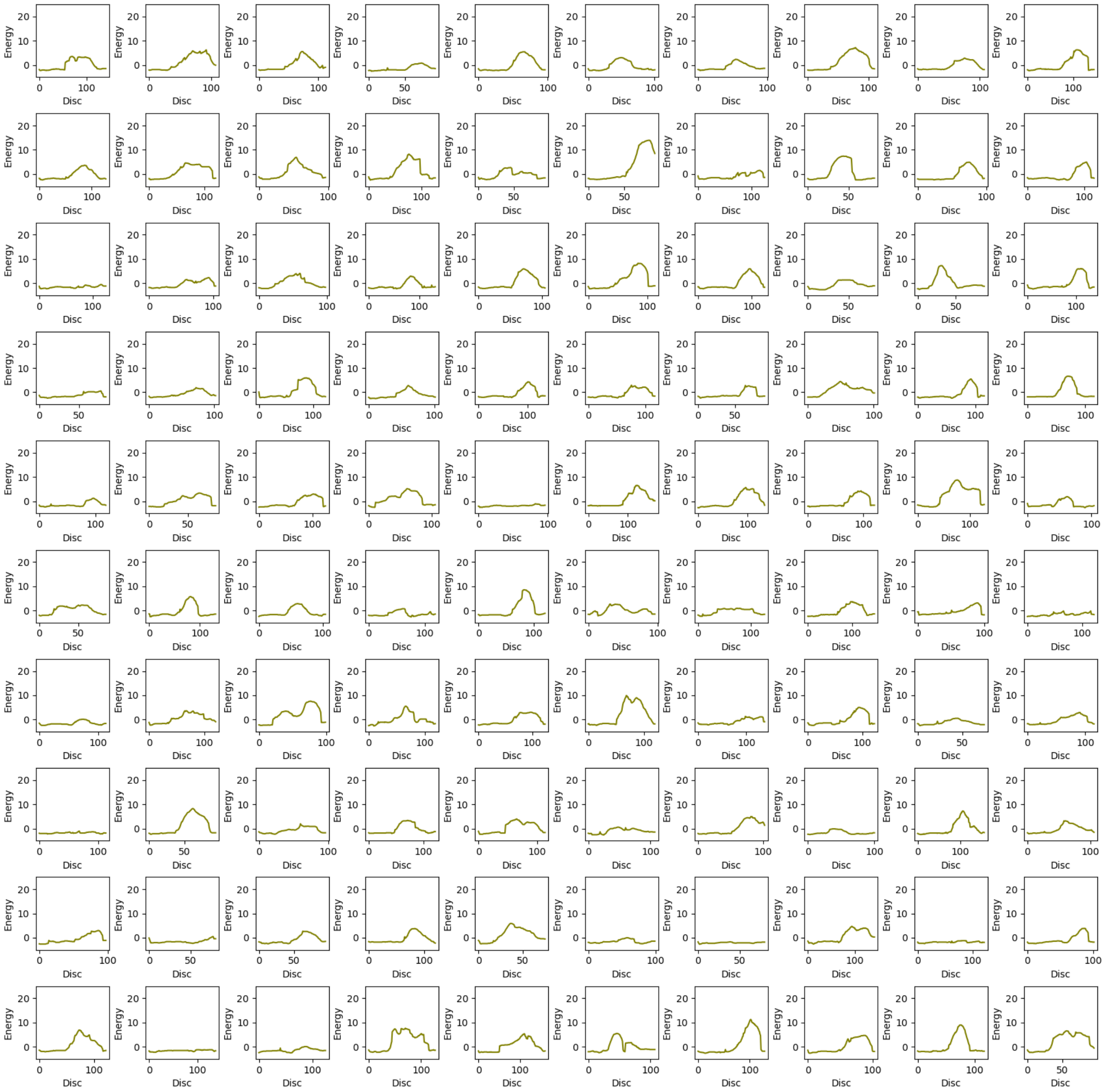

15. Upper Bound energy profile of p3-Wt with 1,2- Dibromoethane (dbe) ligand. The X-axis represents upper bound energy (kcal/mol) and Y-axis represents length of the trajectory [Å], along the disc of tunnel.

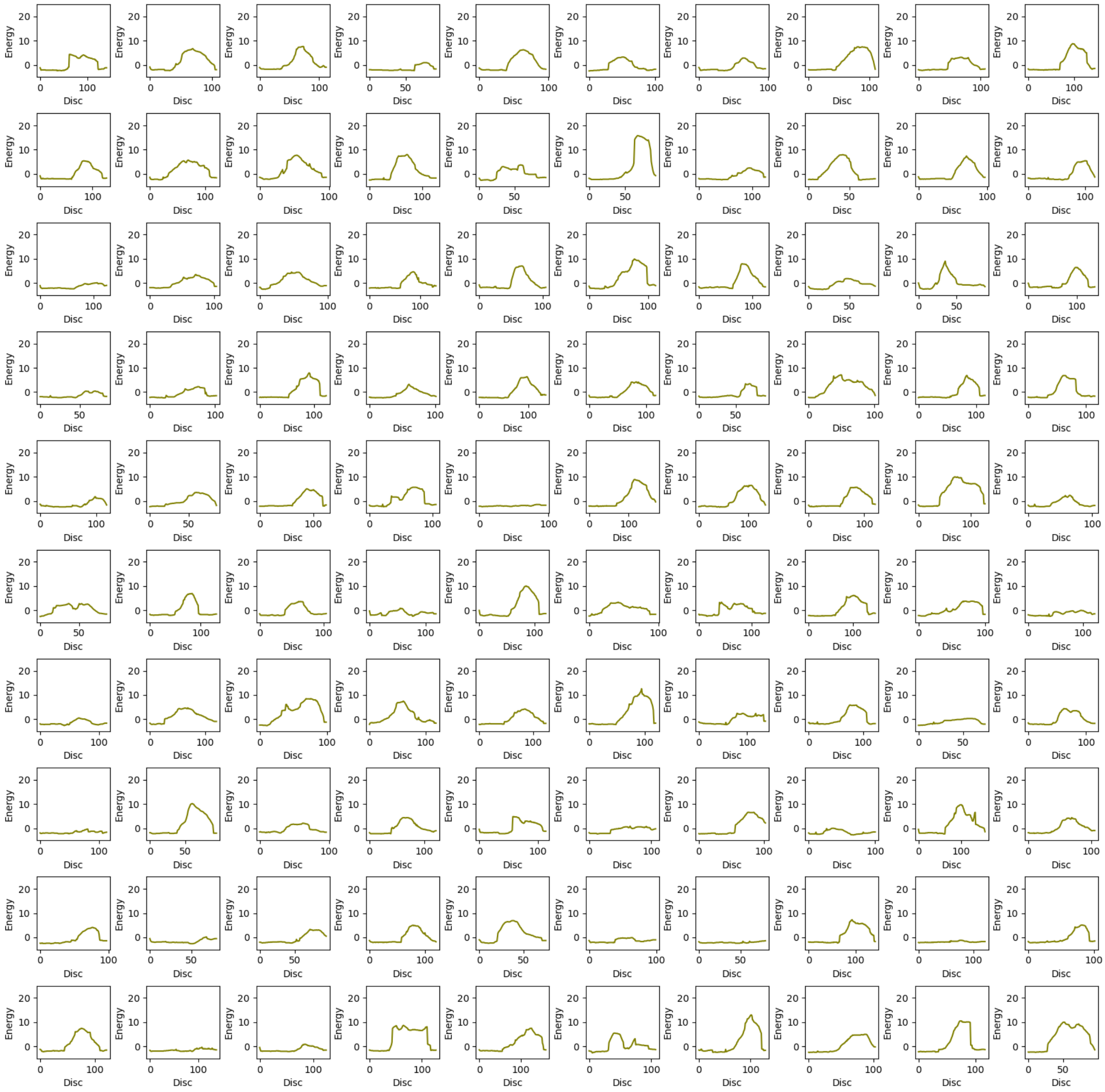

16. Upper Bound energy profile of p3-Wt with Water ($H_2O$) ligand. The X-axis represents upper bound energy (kcal/mol) and Y-axis represents length of the trajectory [Å], along the disc of tunnel.

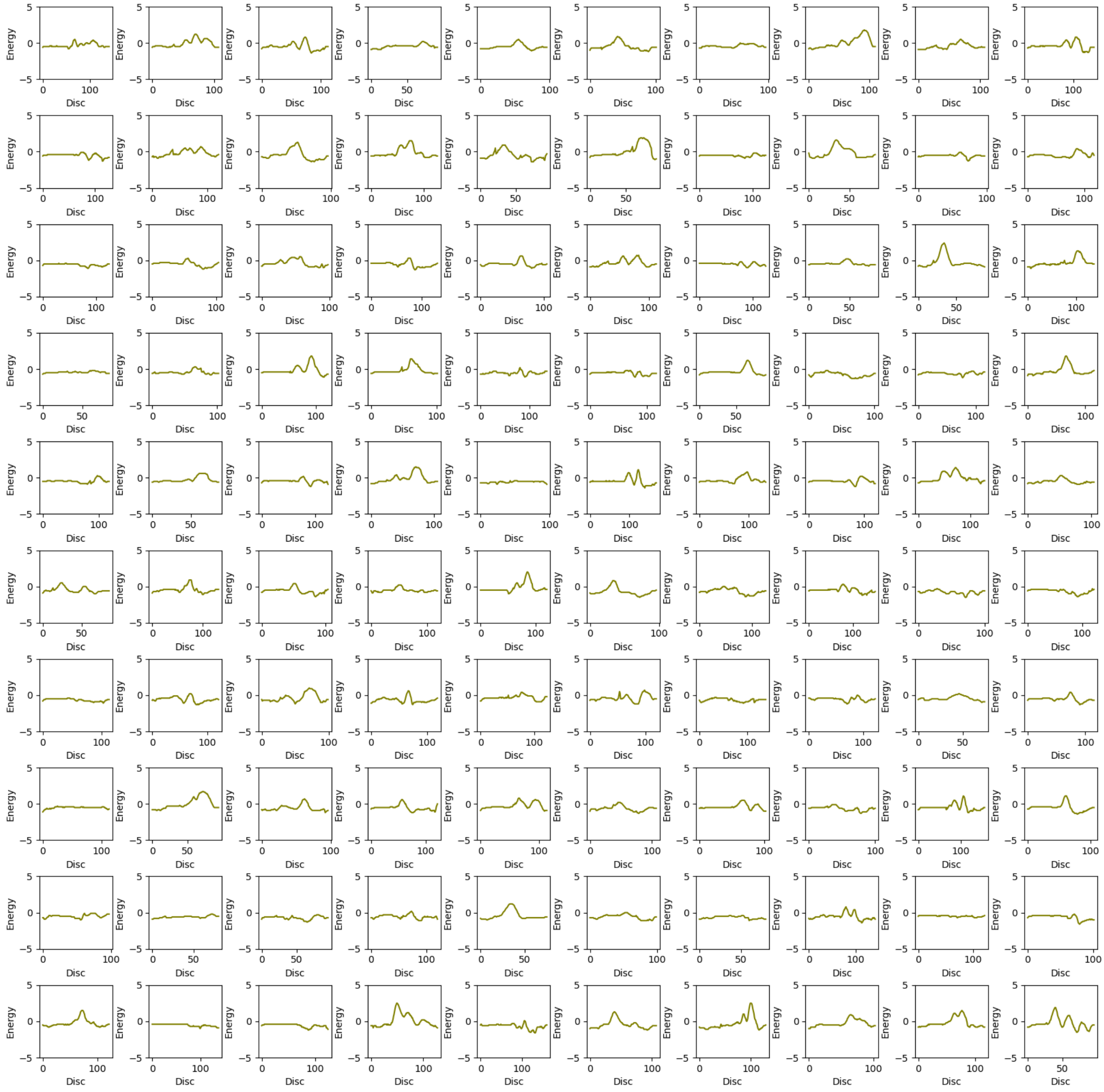

17. Upper Bound energy profile of p3-Closed with Bromide ion ($Br^-$) ligand. The X-axis represents upper bound energy (kcal/mol) and Y-axis represents length of the trajectory [Å], along the disc of tunnel.

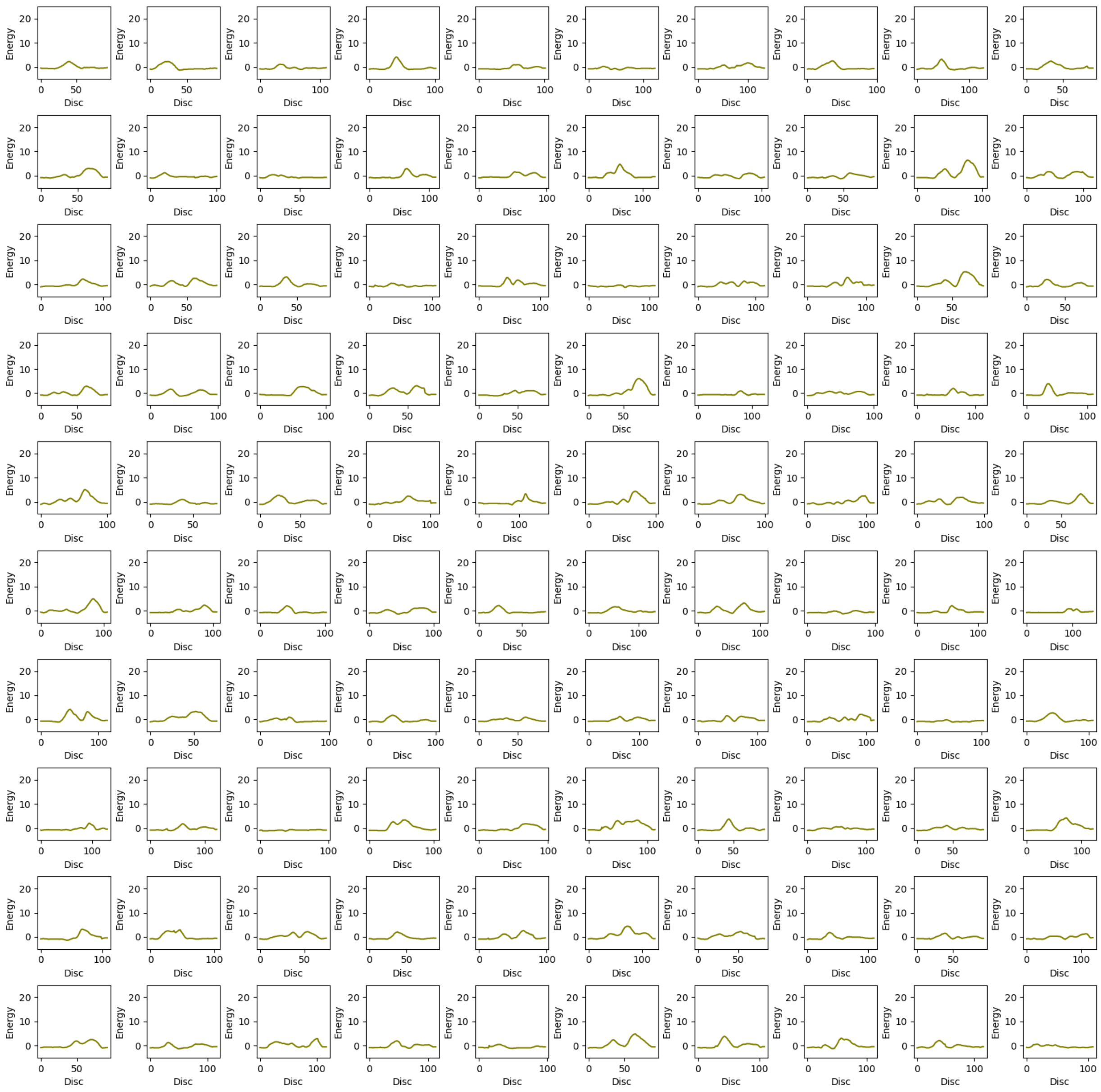

18. Upper Bound energy profile of p3-Closed with 2-Bromoethanol (be) ligand. The X-axis represents upper bound energy (kcal/mol) and Y-axis represents length of the trajectory [Å], along the disc of tunnel.

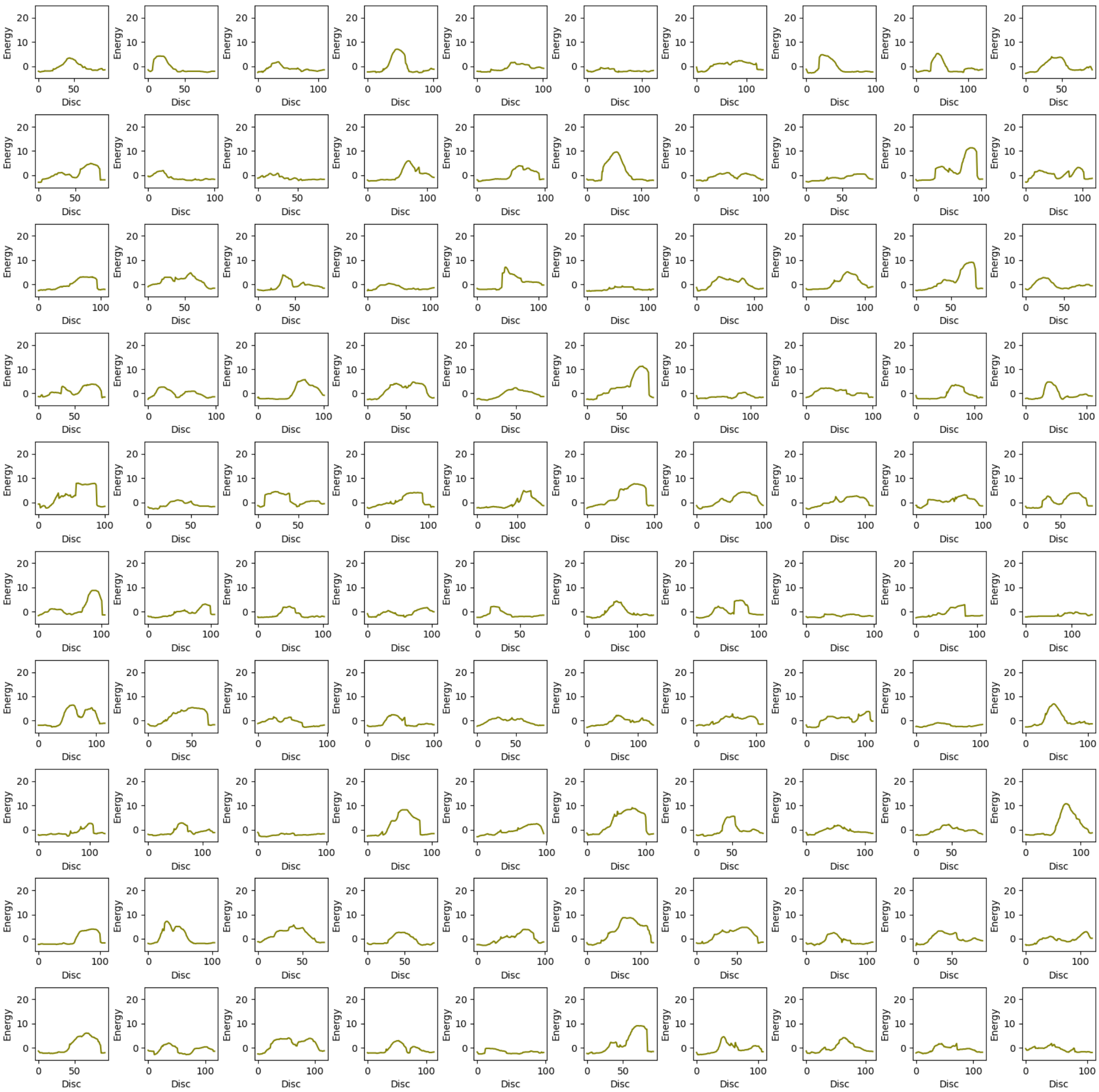

19. Upper Bound energy profile of p3-Closed with 1,2- Dibromoethane (dbe) ligand. The X-axis represents upper bound energy (kcal/mol) and Y-axis represents length of the trajectory [Å], along the disc of tunnel.

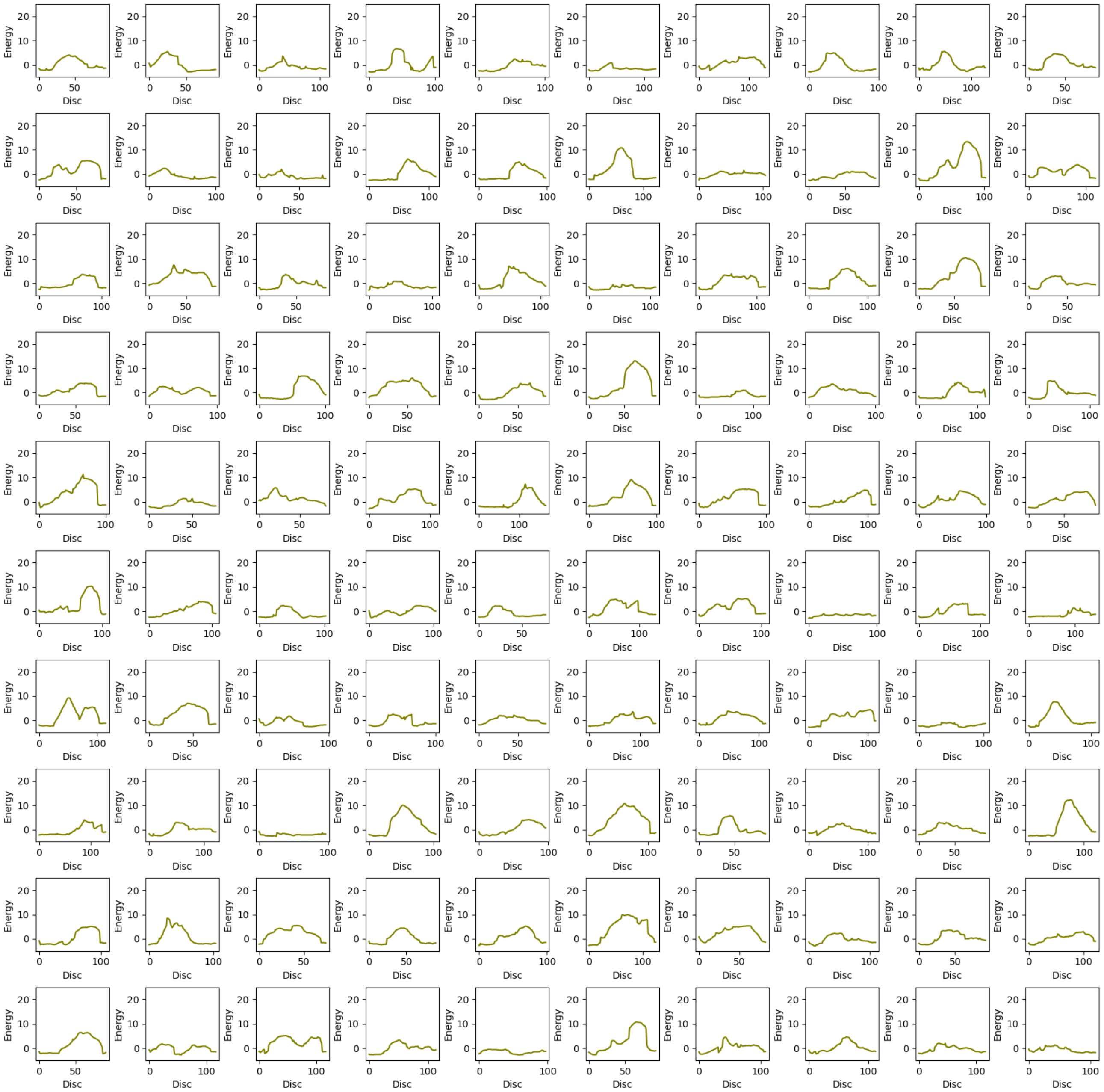

20. Upper Bound energy profile of p3-Closed with Water ($H_2O$) ligand. The X-axis represents upper bound energy (kcal/mol) and Y-axis represents length of the trajectory [Å], along the disc of tunnel.

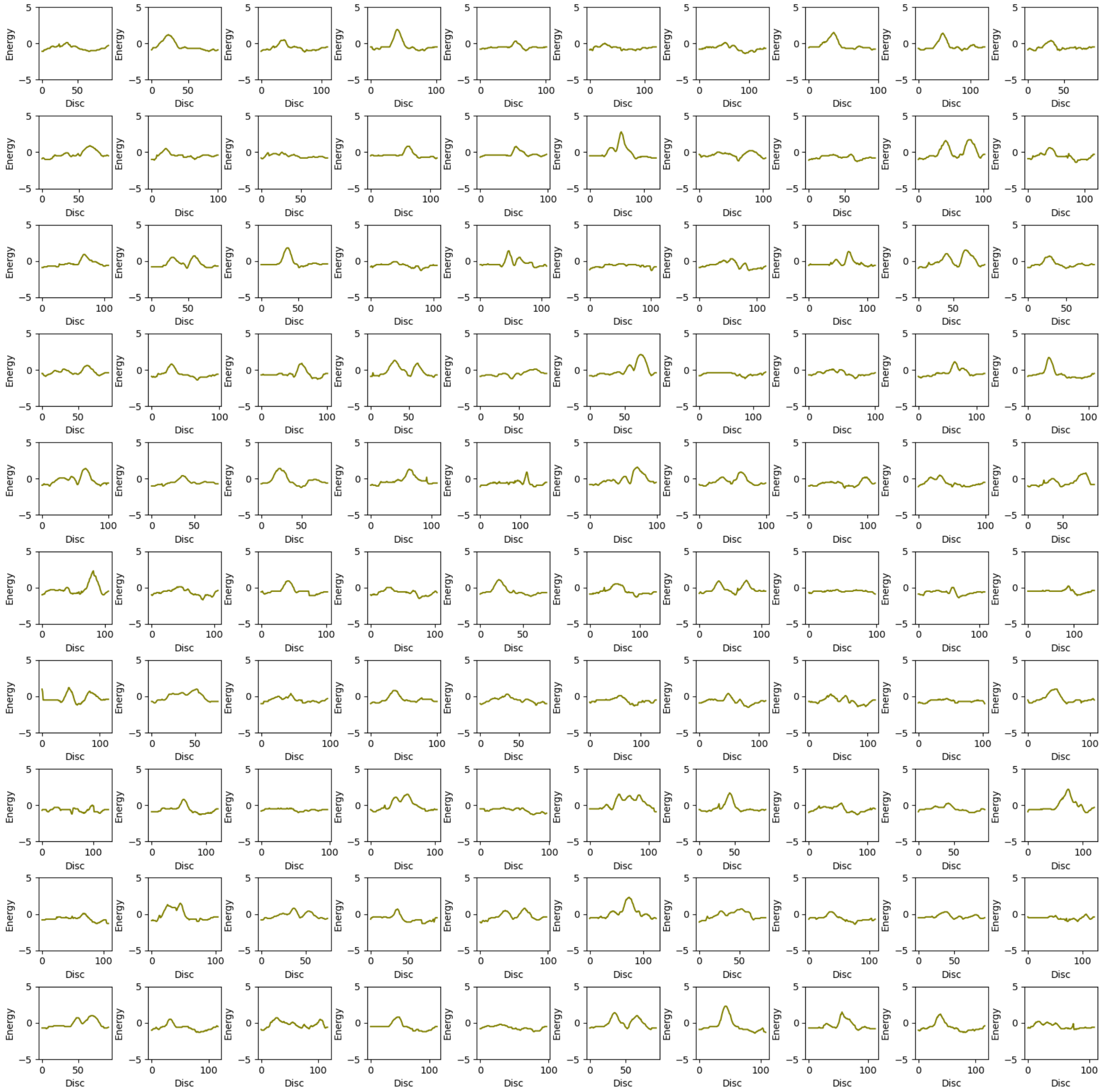

21. Upper Bound energy profile of p3-Open with Bromide ion ($Br^-$) ligand. The X-axis represents upper bound energy (kcal/mol) and Y-axis represents length of the trajectory [Å], along the disc of tunnel.

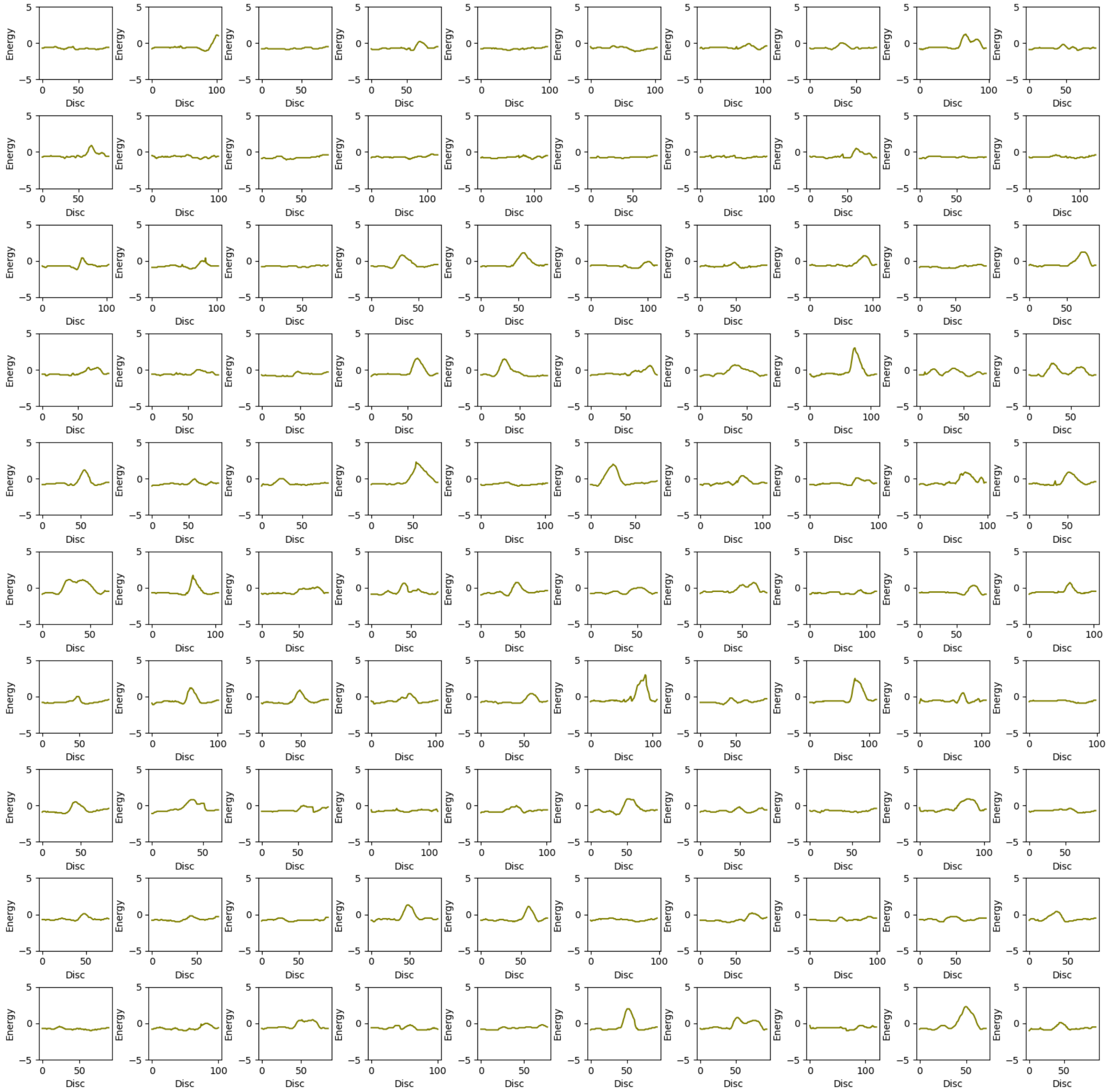

22. Upper Bound energy profile of p3-Open with 2-Bromoethanol (be) ligand. The X-axis represents upper bound energy (kcal/mol) and Y-axis represents length of the trajectory [Å], along the disc of tunnel.

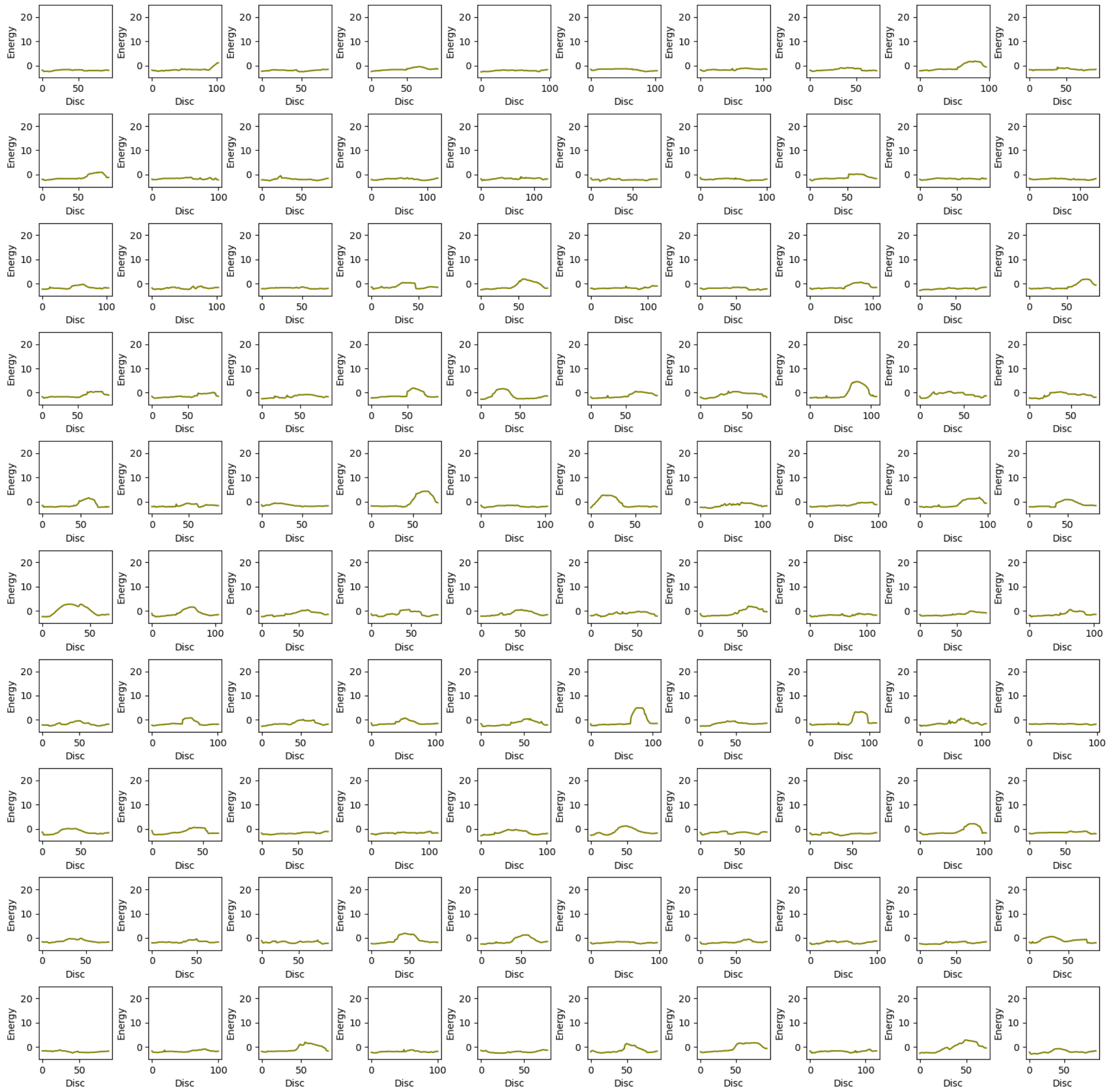

23. Upper Bound energy profile of p3-Open with 1,2- Dibromoethane (dbe) ligand. The X-axis represents upper bound energy (kcal/mol) and Y-axis represents length of the trajectory [Å], along the disc of tunnel.

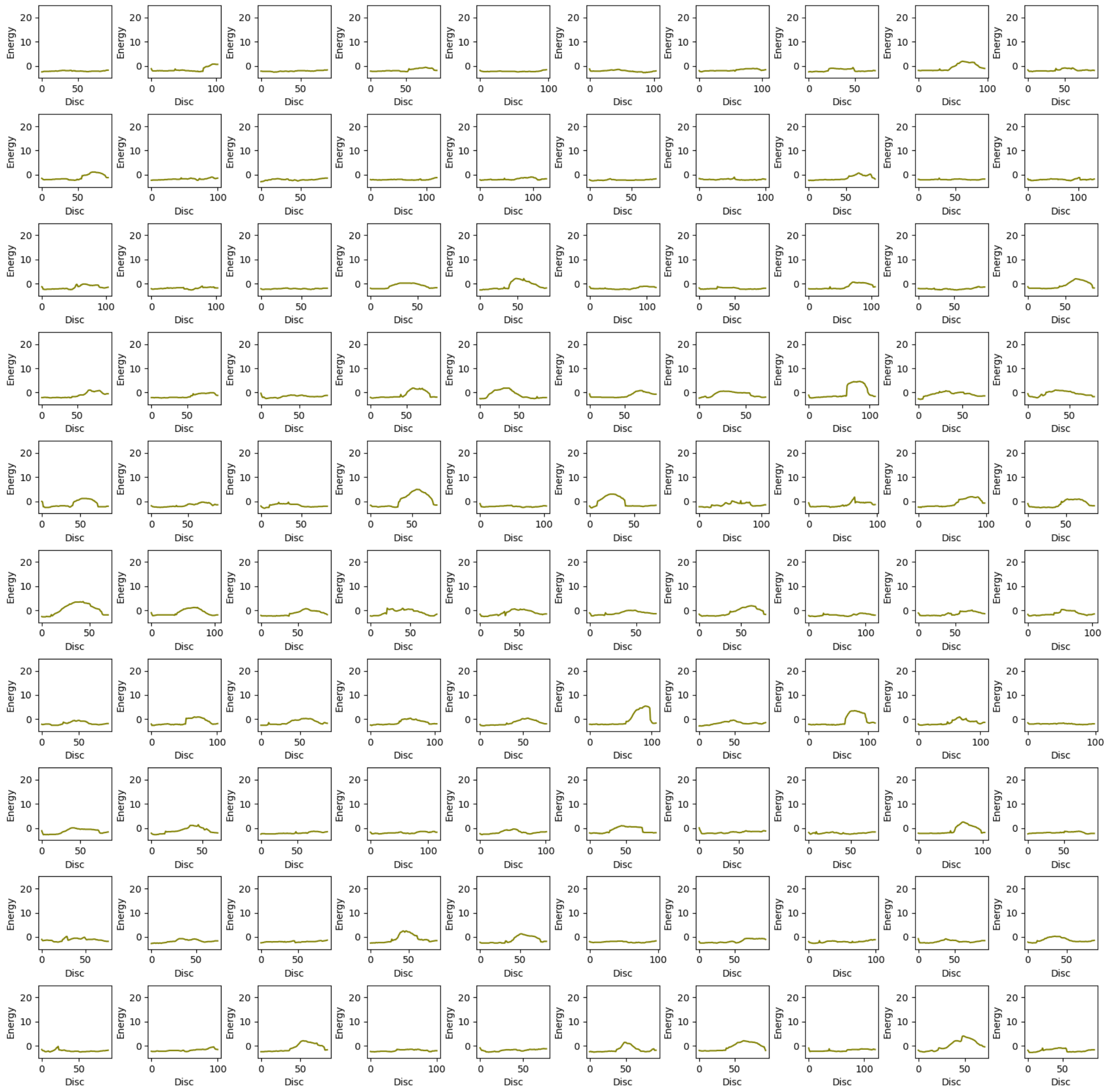

24. Upper Bound energy profile of p3-Open with Water ($H_2O$) ligand. The X-axis represents upper bound energy (kcal/mol) and Y-axis represents length of the trajectory [Å], along the disc of tunnel.

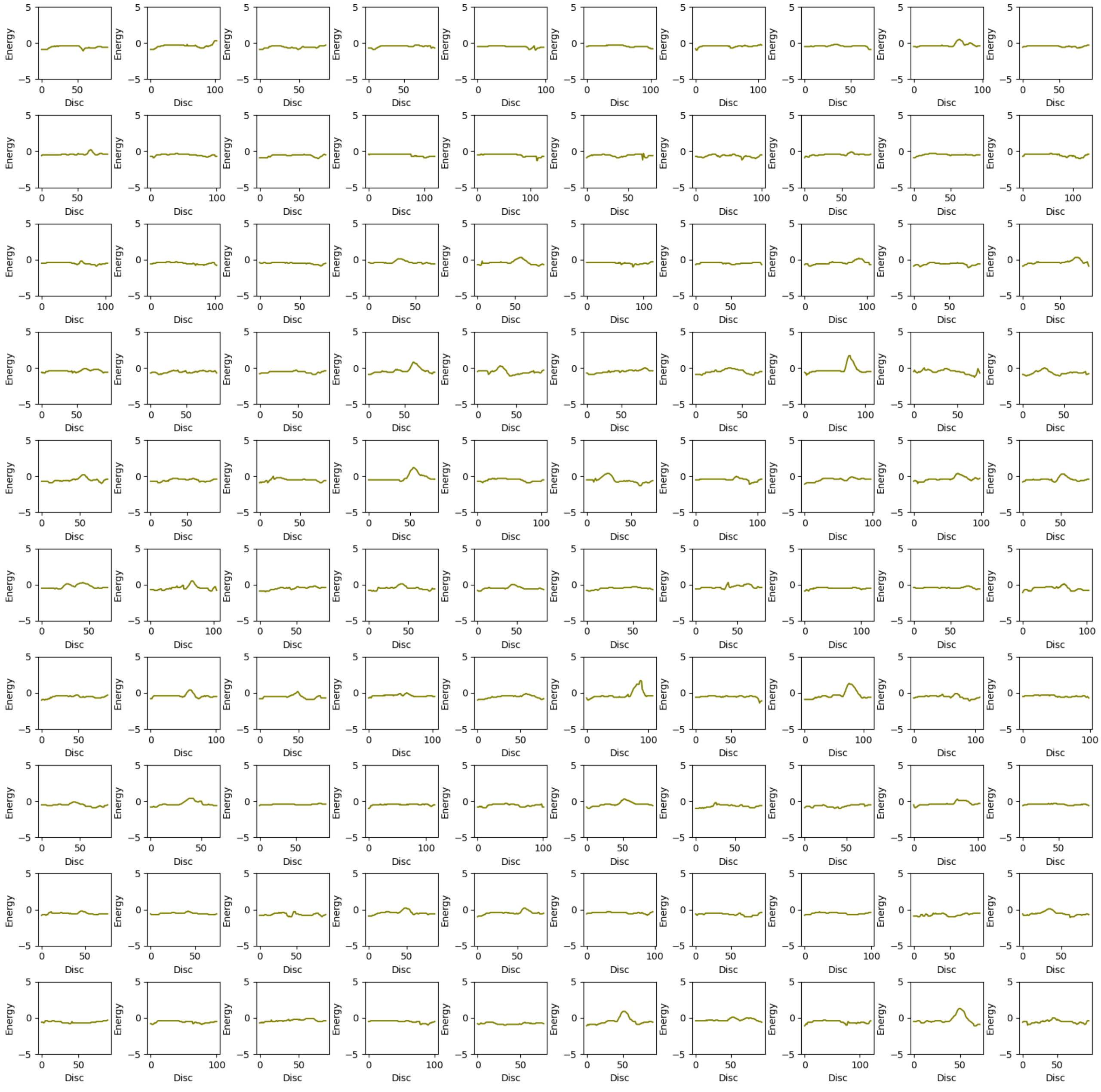

25. Upper Bound energy profile of ST-Wt with Bromide ion ($Br^-$) ligand. The X-axis represents upper bound energy (kcal/mol) and Y-axis represents length of the trajectory [Å], along the disc of tunnel.

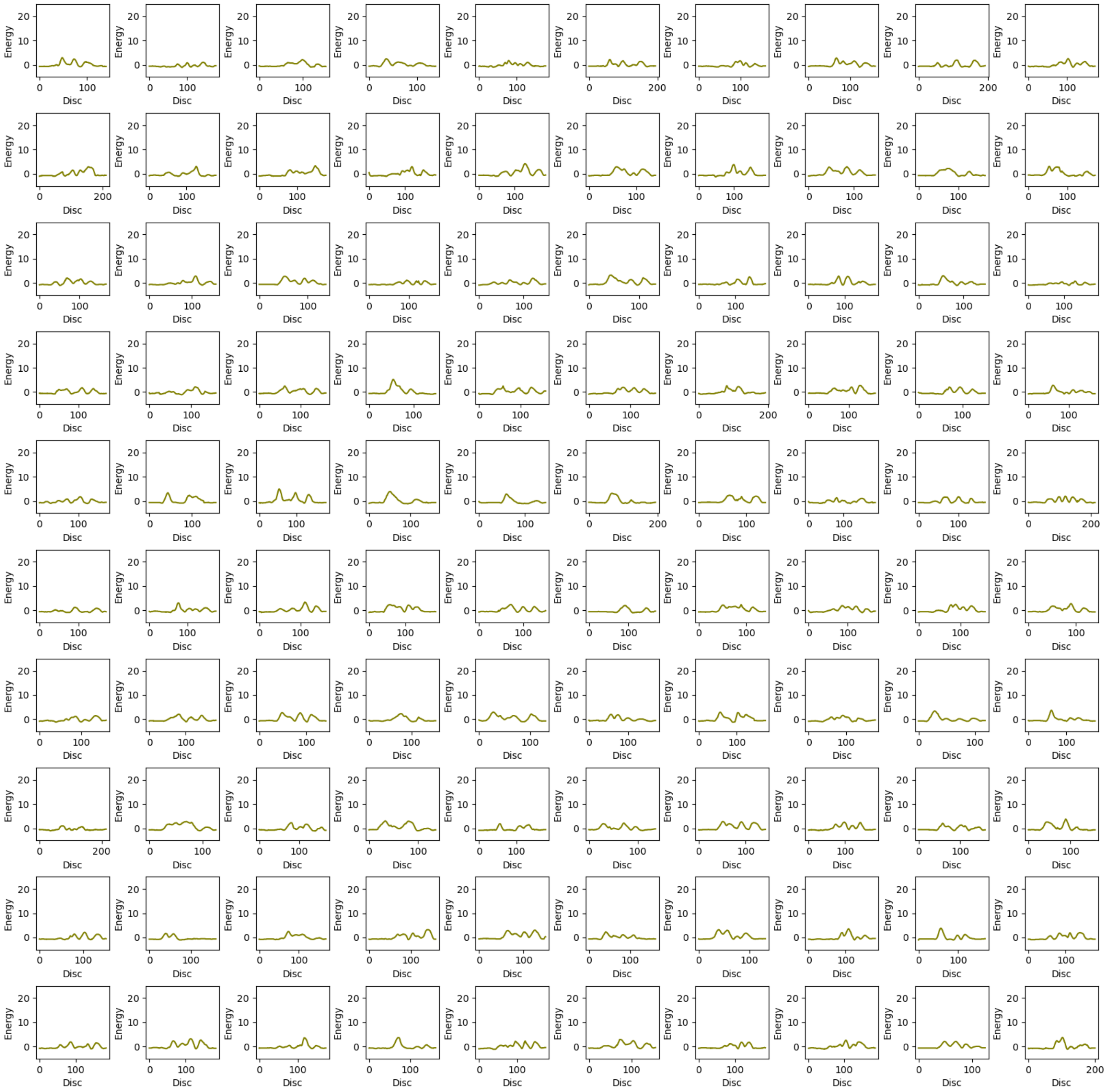

26. Upper Bound energy profile of ST-Wt with 2-Bromoethanol (be) ligand. The X-axis represents upper bound energy (kcal/mol) and Y-axis represents length of the trajectory [Å], along the disc of tunnel.

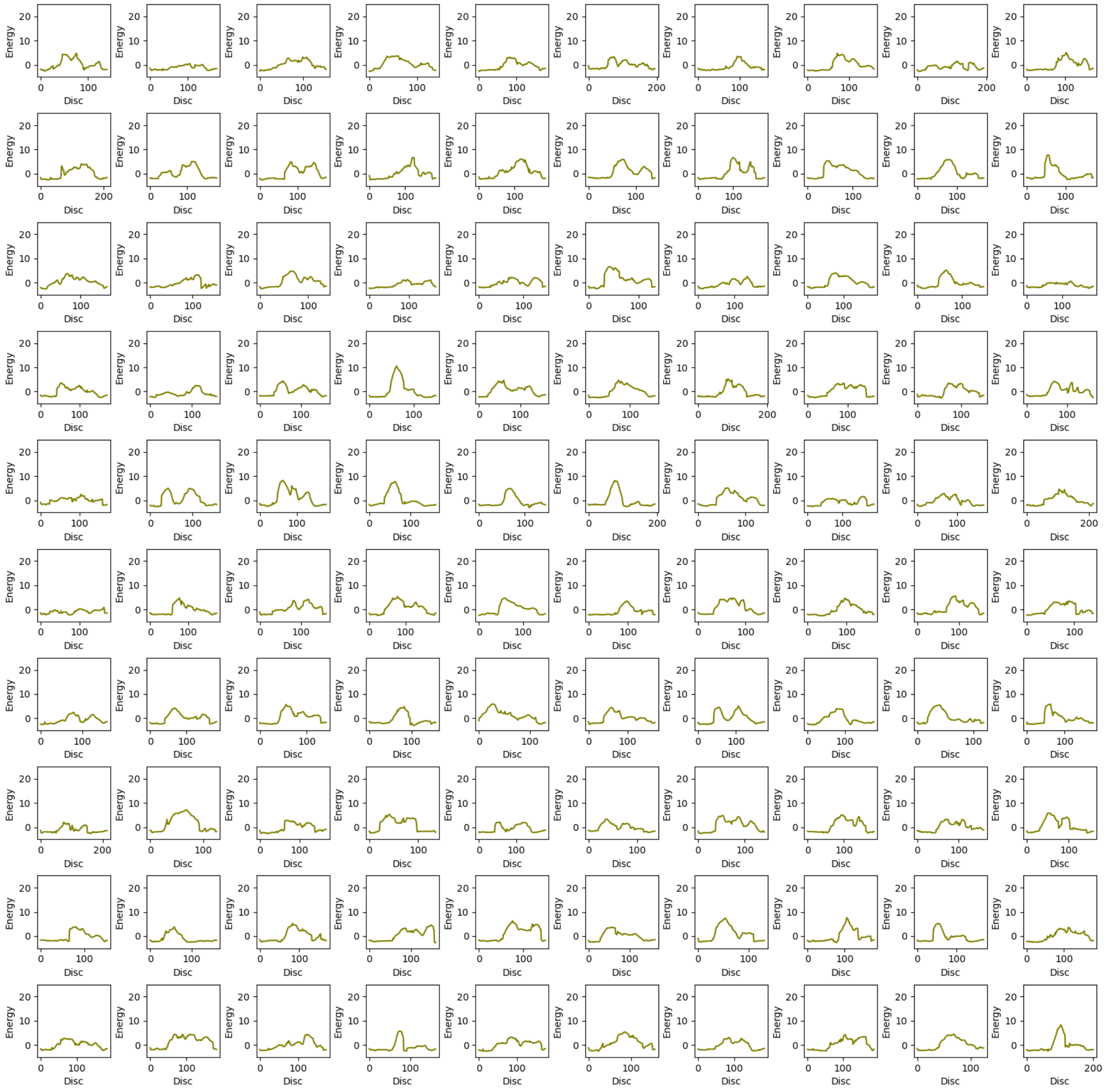

27. Upper Bound energy profile of ST-Wt with 1,2- Dibromoethane (dbe) ligand. The X-axis represents upper bound energy (kcal/mol) and Y-axis represents length of the trajectory [Å], along the disc of tunnel.

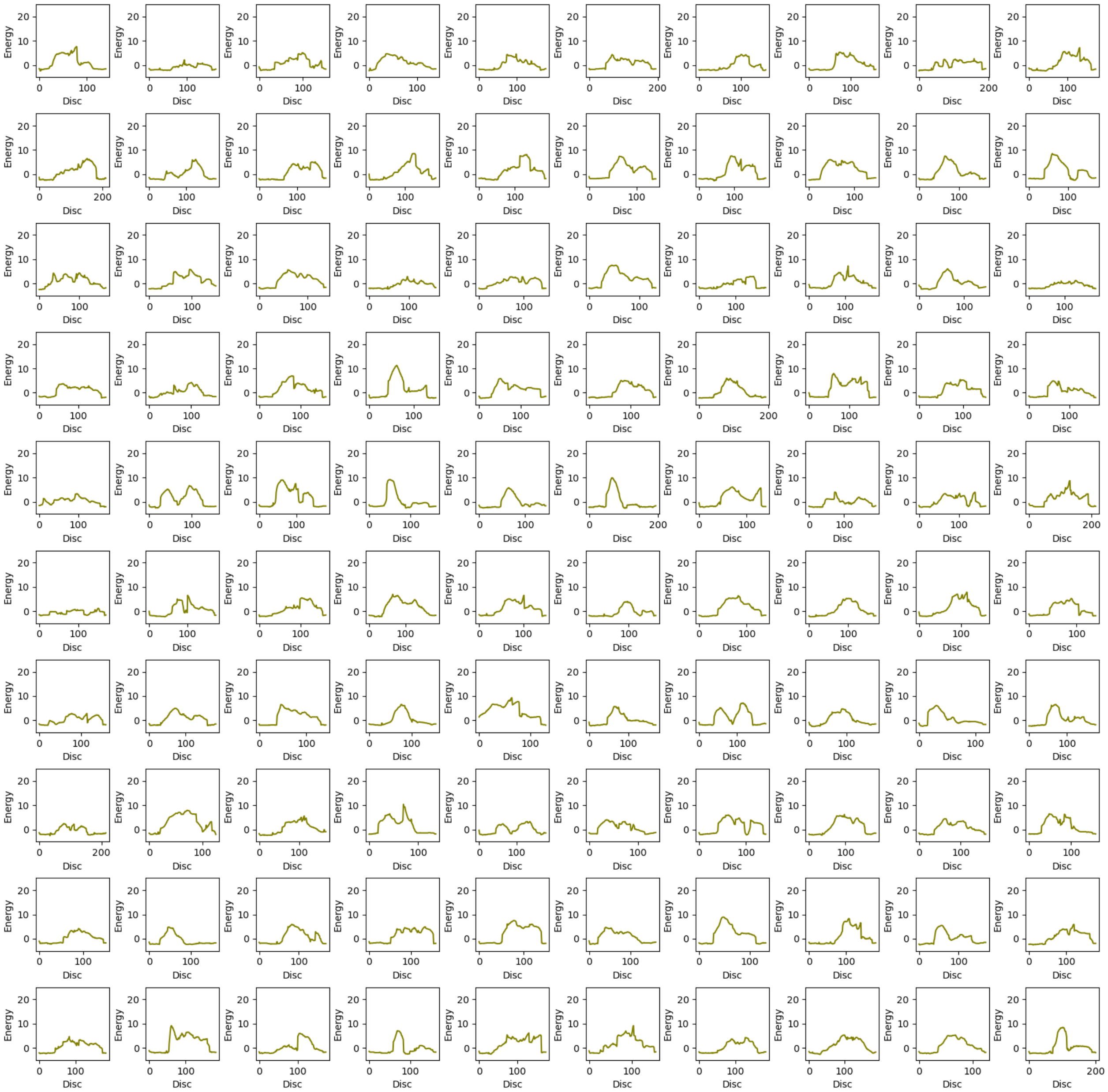

28. Upper Bound energy profile of ST-Wt with Water ($H_2O$) ligand. The X-axis represents upper bound energy (kcal/mol) and Y-axis represents length of the trajectory [Å], along the disc of tunnel.

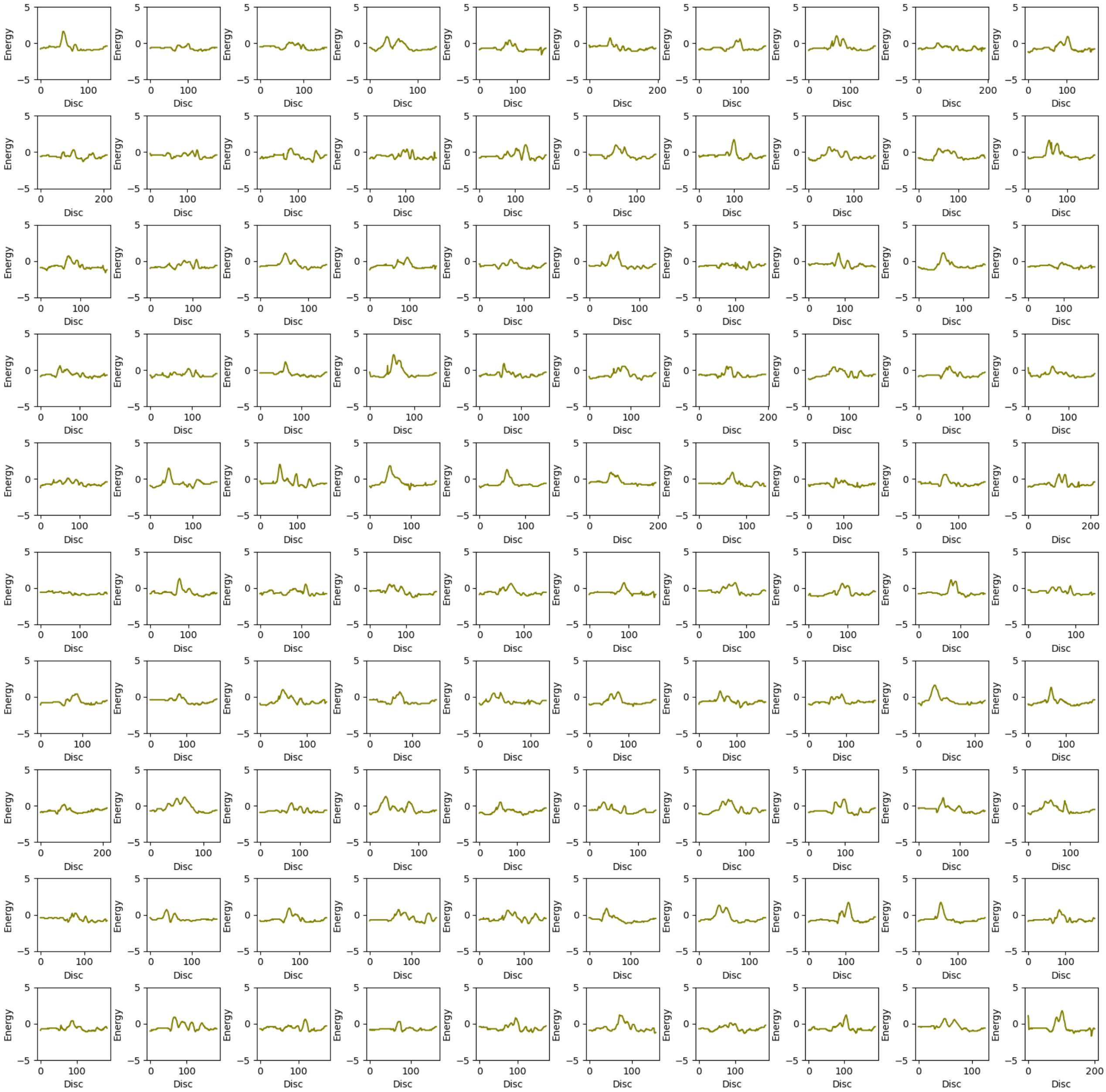

29. Upper Bound energy profile of ST-Closed with Bromide ion ($Br^-$) ligand. The X-axis represents upper bound energy (kcal/mol) and Y-axis represents length of the trajectory [Å], along the disc of tunnel.

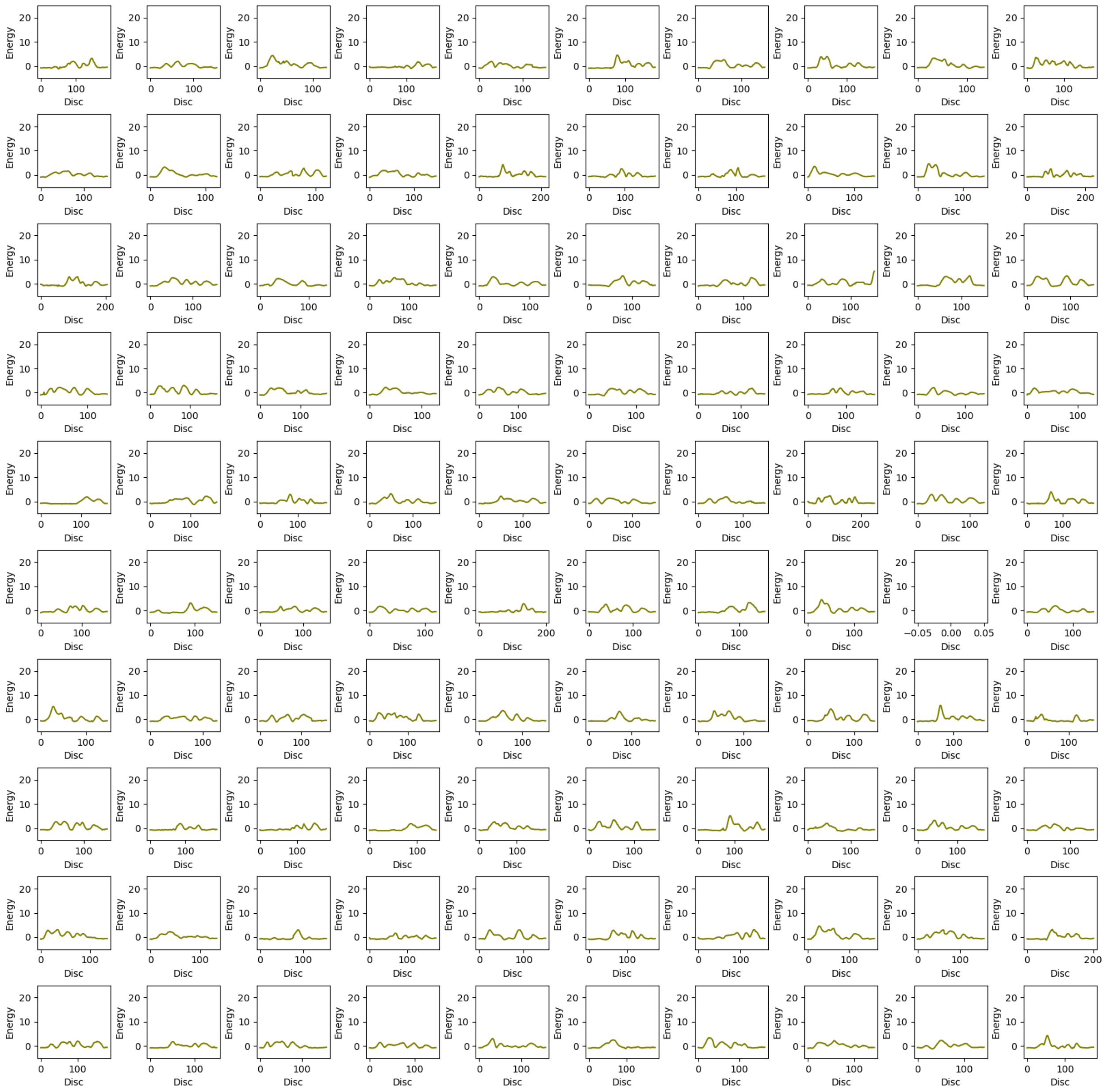

30. Upper Bound energy profile of ST-Closed with 2-Bromoethanol (be) ligand. The X-axis represents upper bound energy (kcal/mol) and Y-axis represents length of the trajectory [Å], along the disc of tunnel.

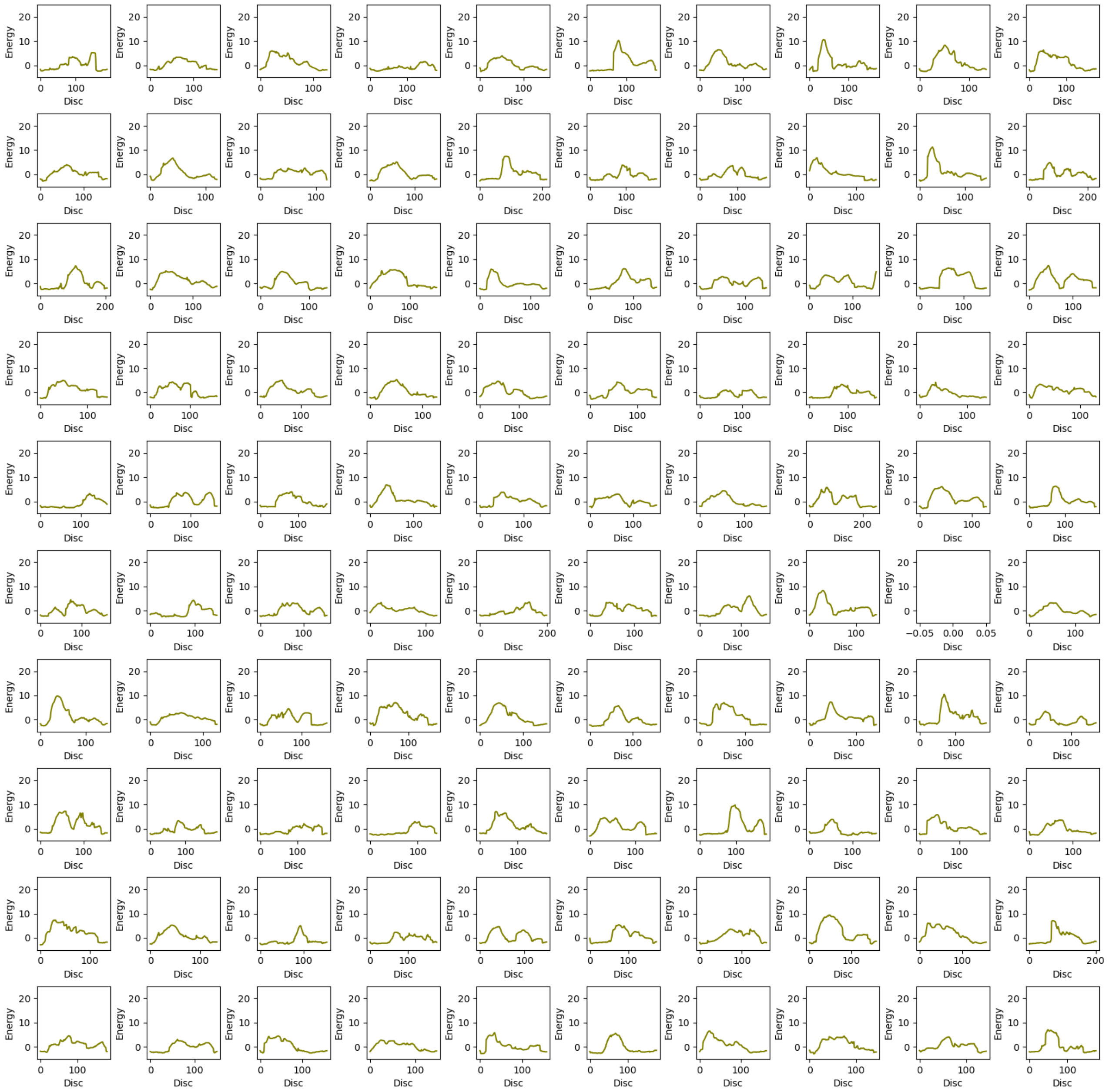

31. Upper Bound energy profile of ST-Closed with 1,2- Dibromoethane (dbe) ligand. The X-axis represents upper bound energy (kcal/mol) and Y-axis represents length of the trajectory [Å], along the disc of tunnel.

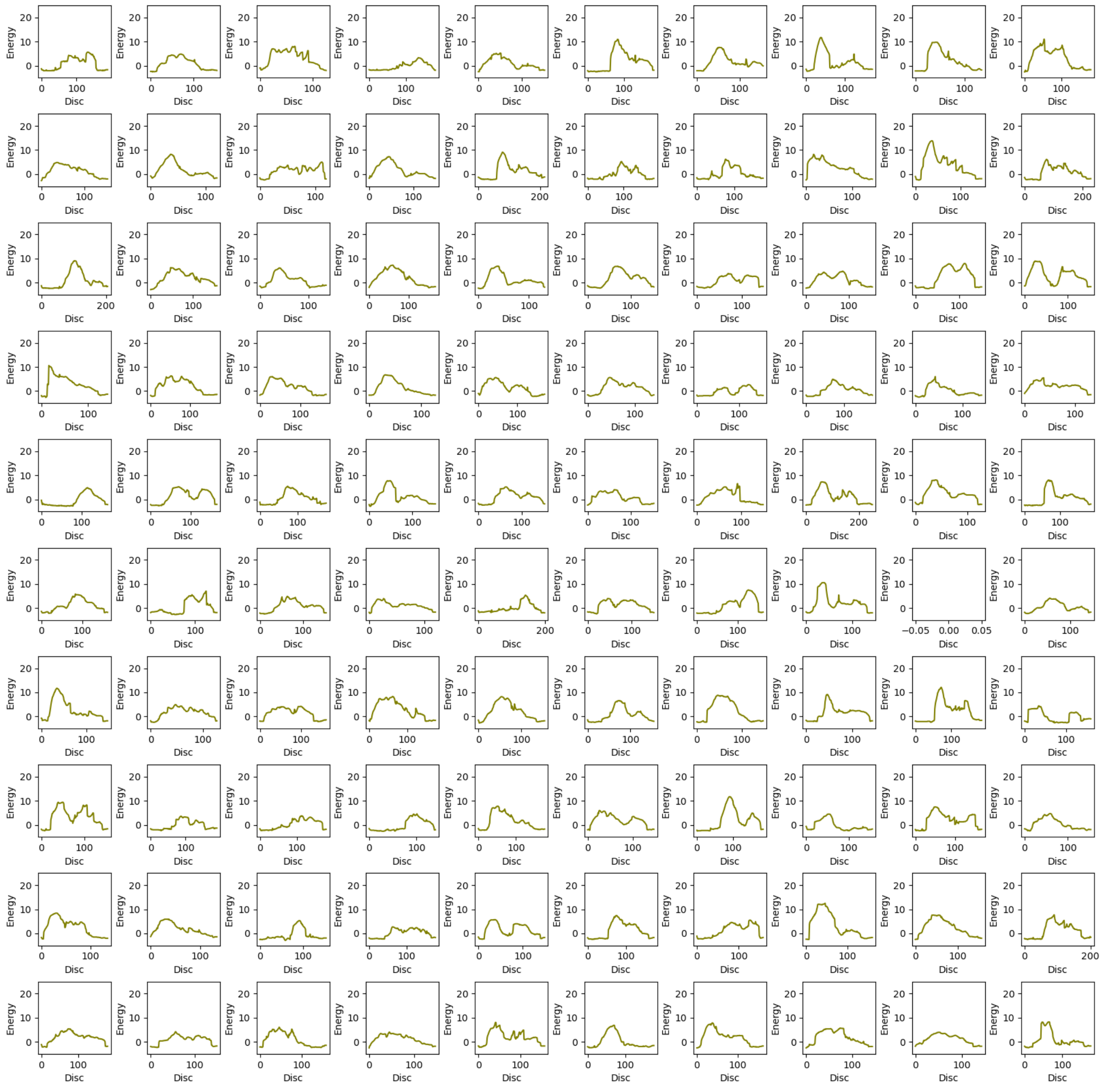

32. Upper Bound energy profile of ST-Closed with Water ($H_2O$) ligand. The X-axis represents upper bound energy (kcal/mol) and Y-axis represents length of the trajectory [Å], along the disc of tunnel.

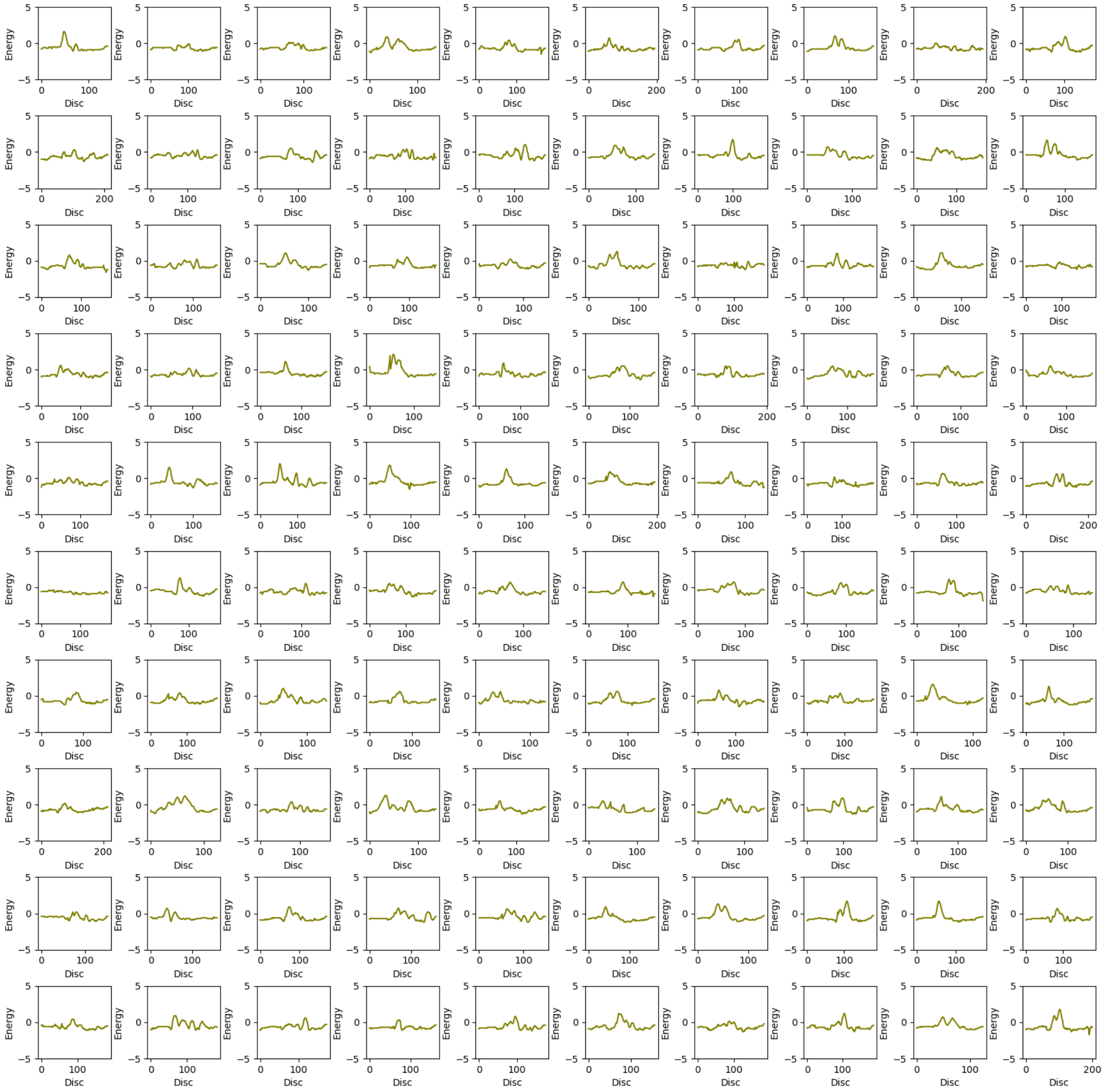

33. Upper Bound energy profile of ST-Open with Bromide ion ($Br^-$) ligand. The X-axis represents upper bound energy (kcal/mol) and Y-axis represents length of the trajectory [Å], along the disc of tunnel.

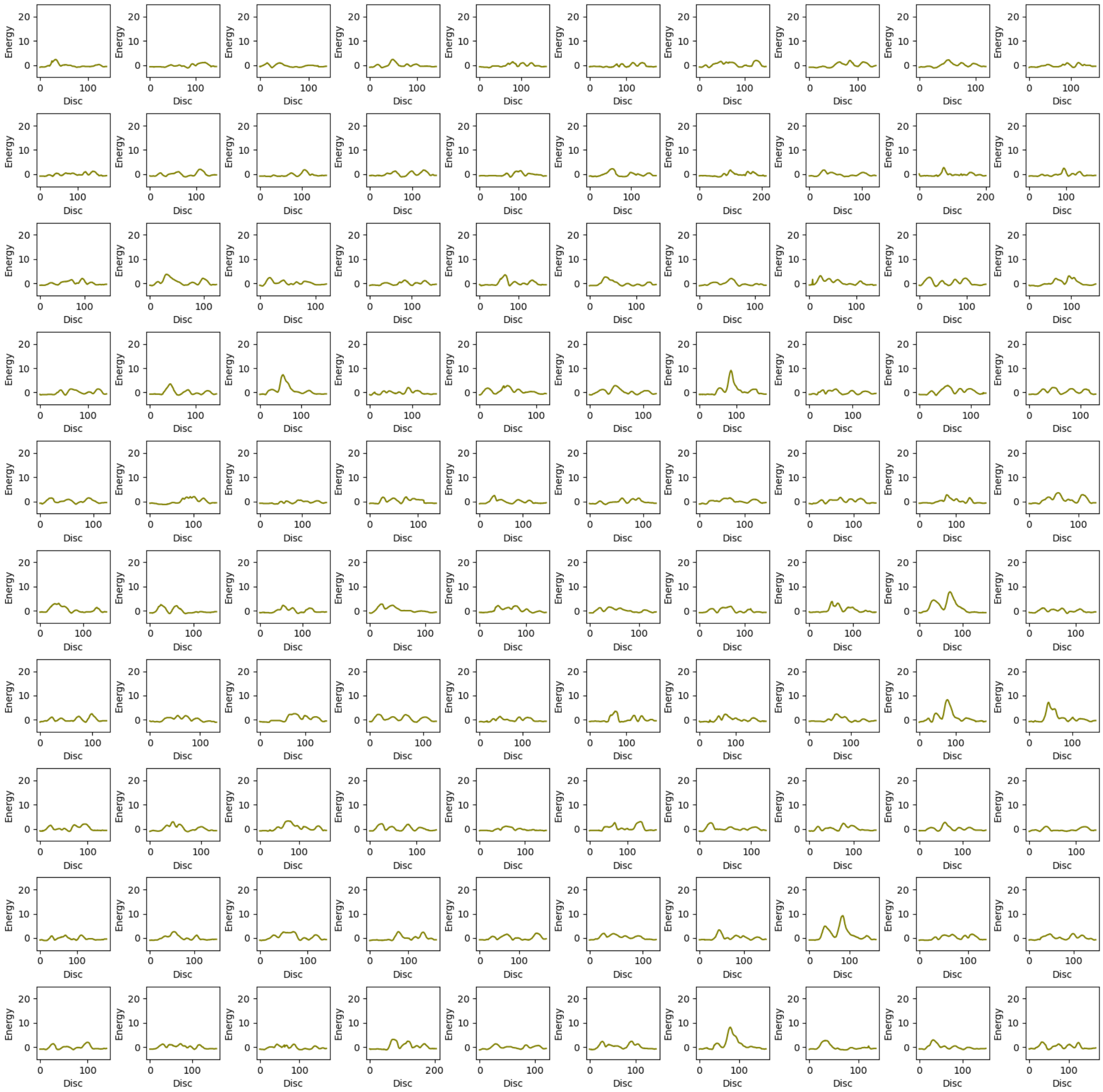

34. Upper Bound energy profile of ST-Open with 2-Bromoethanol (be) ligand. The X-axis represents upper bound energy (kcal/mol) and Y-axis represents length of the trajectory [Å], along the disc of tunnel.

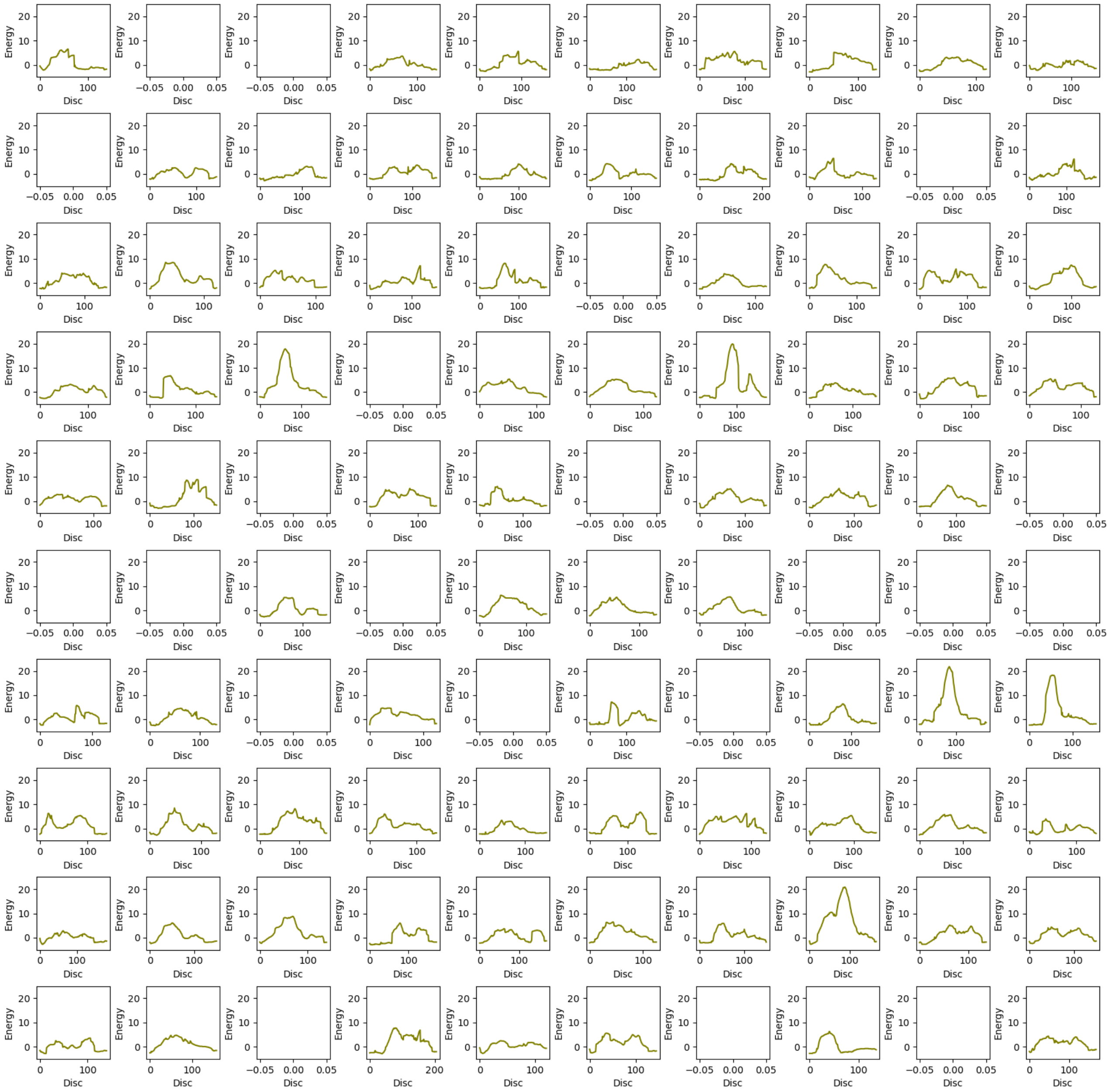

35. Upper Bound energy profile of ST-Open with 1,2- Dibromoethane (dbe) ligand. The X-axis represents upper bound energy (kcal/mol) and Y-axis represents length of the trajectory [Å], along the disc of tunnel.

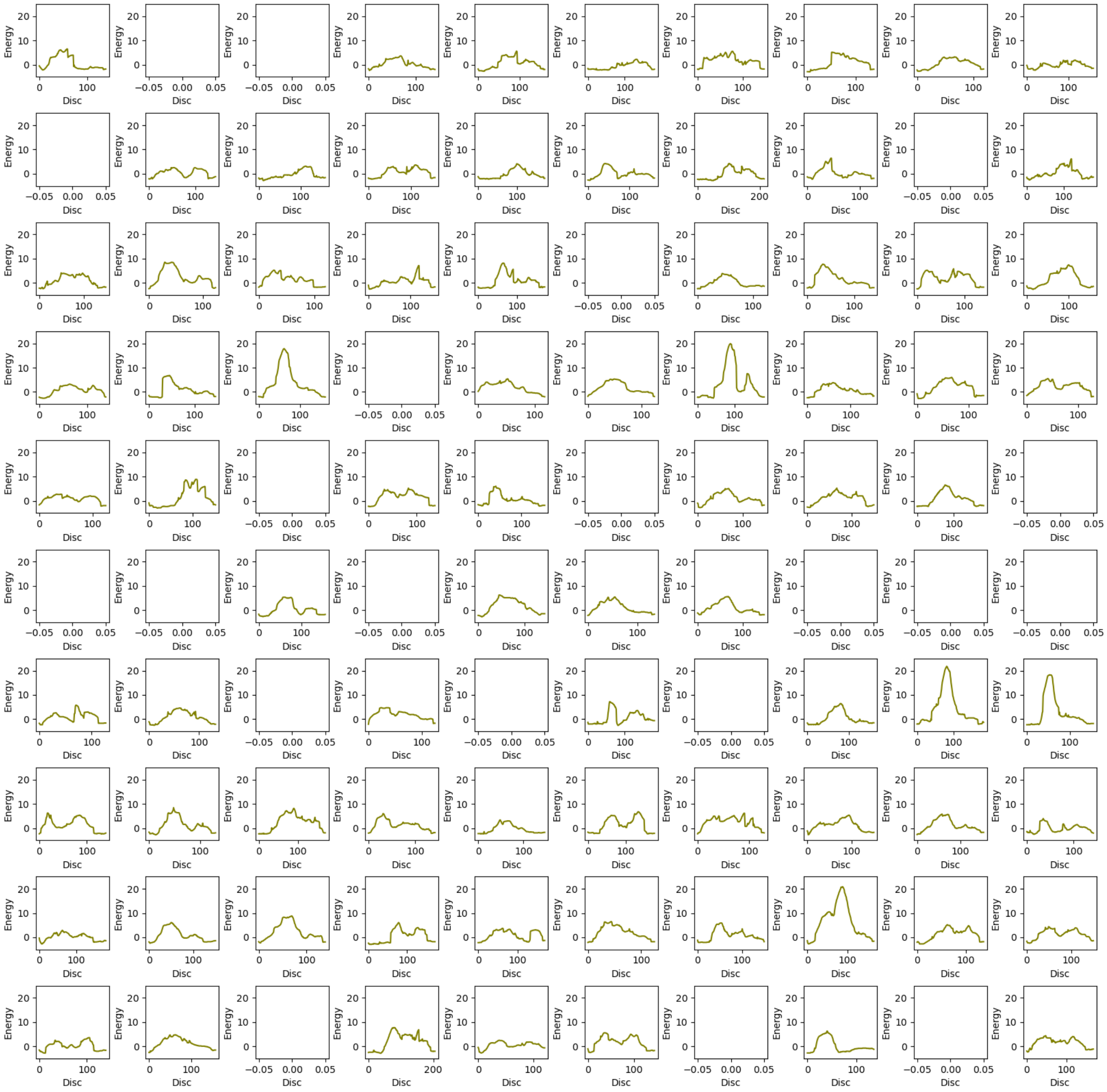

36. Upper Bound energy profile of ST-Open with Water ($H_2O$) ligand. The X-axis represents upper bound energy (kcal/mol) and Y-axis represents length of the trajectory [Å], along the disc of tunnel.

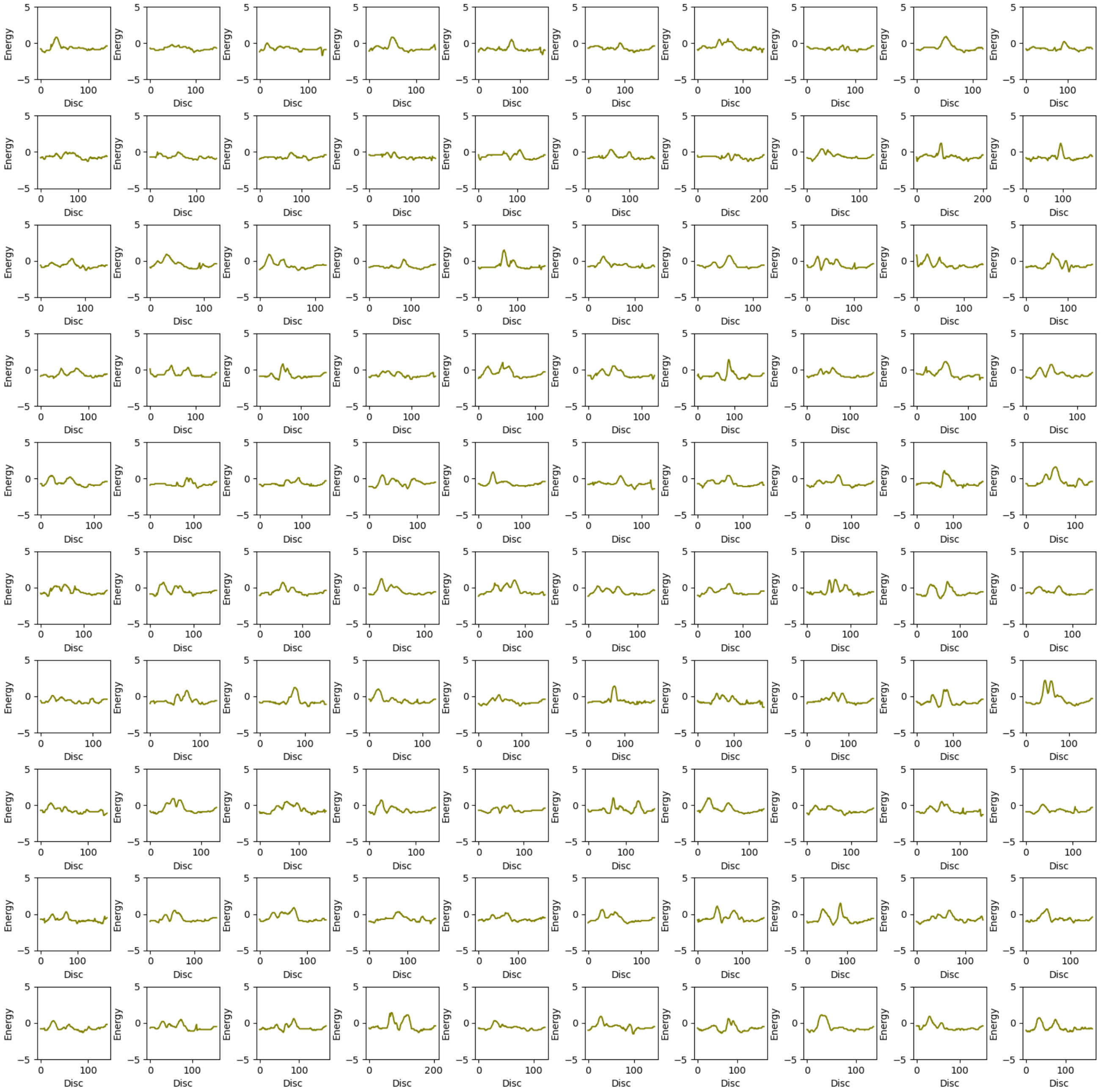
